# STORMM: Structure and TOpology Replica Molecular Mechanics for chemical simulations

**DOI:** 10.1101/2024.03.27.587048

**Authors:** David S. Cerutti, Rafal Wiewiora, Simon Boothroyd, Woody Sherman

## Abstract

The **S**tructure and **TO**pology **R**eplica **M**olecular **M**echanics (STORMM) code is a next-generation molecular simulation engine and associated libraries optimized for performance on fast, multicore central processor units (CPUs) and graphics processing units (GPUs) with independent memory and tens of thousands of threads. STORMM is built to run thousands of independent molecular mechanical calculations on a single GPU with novel implementations that optimize numerical precision, mathematical operations, throughput, and resource management. The libraries are built around accessible classes with detailed documentation, supporting fine-grained parallelism and algorithm development as well as macroscopic manipulations of groups of systems on and off of the GPU. A primary intention of the STORMM libraries is to provide developers of atomic simulation methods with access to a high-performance molecular mechanics engine with extensive facilities to prototype and develop bespoke tools aimed toward drug discovery applications. In its present state, STORMM delivers molecular dynamics simulations of small molecules and small proteins in implicit solvent with tens to hundreds of times the throughput of conventional codes. The engineering paradigm also transforms two of the most memory bandwidth-intensive aspects of condensed-phase dynamics, particle-mesh mapping and valence interactions, into compute-bound problems for several times the scalability of existing programs. Numerical methods for getting the most out of each bit of information present in stored coordinates and lookup tables are also presented, delivering improved accuracy over methods implemented in other molecular dynamics engines. The open-source code is released under the MIT license.

## 1 Introduction

Molecular simulations in drug discovery can provide unique insights and predictions related to statistical thermodynamic processes foundational to biology that are currently inaccessible with other methods. Applications include protein-ligand binding,[1, 2] allosteric modulation,[3, 4] membrane permeability,[5] and solubility[6] to name but a few. However, molecular simulations are intensive calculations in which the mutual interactions of all atoms in the system are computed and propagated motion over millions, billions, or even trillions of iterations to reach nanosecond, microsecond, and millisecond timescales, respectively. The scale, speed, accuracy, and applicability of the molecular simulations is a matter of the computational resources, algorithmic efficiency, multi-thread parallelism, and flexibility of the code. There is an intricate balance between the underlying computational hardware and the simulation engine – as computer architectures continue to evolve, it is important for molecular simulation packages to keep pace and leverage hardware advances for domain-specific applications.

Two types of hardware have come to dominate the high-performance computing space: general purpose central processing units (CPUs) and general purpose graphics processing units (GPUs). The term “general purpose” refers to the ease of programming each device in familiar languages with access to a breadth of common mathematical functions and matrix algebra, an important set of capabilities but not the entirety of what modern computers do. A “general purpose” CPU is not limited to operating as a single thread performing each operation in series: aside from the multicore nature of modern CPUs, each processor core has vectorized instructions to operate on aligned data. Expert programmers arrange their codes such that the compiler can identify and implement these optimizations. On the other hand, a “general purpose” GPU does not implement specialized classes like C++ maps or regular expressions. A GPU would not be useful for input parsing unless there are kernels to operate on bulk quantities of character arrays to apply a few common interpretations such as formatted numerical translation. The algorithmic advantages of each device (CPUs and GPUs) have supported a symbiosis in modern scientific computing while the common programming languages have enabled extreme performance enhancements in code bases with long histories.

The arithmetic units of CPUs and GPUs share much in common, but the recent success of GPUs in scientific computing owes just as much to a versatile memory model. While there is a marked difference in the abundance of data cache on each device, the memory hierarchies are so similar that “white board” strategies for solving problems often translate from one device to the other. GPUs offer a non-uniform memory access (NUMA) system, a common pool of memory that all processors can read and write, which has been a hallmark of many successful parallel computing systems dating to the 1990s. Optimizations in the compilers that allow GPUs to interlace arithmetic operations between high-latency memory accesses, “hiding” the most costly communication, have also been instrumental to their success. The memory in GPU devices is distinct from that accessed by CPUs on the host system, but the PCIE bus connecting the two is simple to utilize and the communication overhead can be negligible if there is enough arithmetic applied to the transferred data. The ease of communication, coupled with powerful and automated methods for dampening the impact of latency, has placed GPUs as the essential components of advanced compute resources.

The affordable nature of commodity gaming GPU cards facilitates a broad base of developers and users while the HPC market evolves to accommodate specialized applications that require changes in the underlying chip design. For example, neural network training workloads may lead to the advent of new and powerful “general purpose” tensor processing units in a the coming years. As a minor share of the GPU consumer base, the molecular simulation community has little sway the design decisions for the next generation of processor architectures and therefore we must tune our codes to reflect the changing architectures. The growing demand for artificial intelligence and generative models will likely continue to influence chip designs toward extreme performance for deep learning, yielding comparatively less benefit for simulation codes optimized for older architectures. Simultaneously, the gaming industry will continue to drive demand for commodity chips, which perform massive amounts of arithmetic operations useful to many scientific applications, albeit with a reduction or outright lack of high-precision floating point operations. Other possible developments such as CPU-adjacent Field Programmable Gate Arrays (FPGAs) or integrated general-purpose graphics processors could even make transitions to high-performance coding simpler by removing a partition that currently exists between the memory banks in graphics cards and those of the host computer.[7]

Despite significant changes in GPU architectures, the architectures of molecular simulation programs are based on codes developed fifteen years ago, when the first GPU-accelerated molecular dynamics (MD) codes became available.[8, 9] While chemical simulations with classical particles have reached time scales orders of magnitude higher than would be possible even with current CPU resources,[10] limits are being reached in terms of saturating modern GPUs to maximize performance. Other aspects of computational chemistry such as docking, coarse-grained simulations, and quantum mechanical calculations have not been as quick to adapt to the new architectures, although progress is taking place.[11, 12] There are many areas where high-performance computing could accelerate other problems in the field if they were posed in a manner amenable to the hardware. There is no universal software infrastructure with optimized GPU capabilities spanning the breadth of molecular simulations (minimization, molecular dynamics, docking, conformational search, etc.).

In order to facilitate the deployment of molecular simulation methods on modern hardware architectures and contribute to a community of developers to advance new methods for drug discovery, we present the STORMM (**S**tructure and **TO**pology **R**eplica **M**olecular **M**echanics) libraries, based on a collection of refined and interoperable C++ classes. These libraries provide a means to accumulate and organize a series of topologies and coordinate systems into collated arrays of parameters and chains of work units, enabling many independent problems to coexist in a single runtime process, thereby maximizing GPU utilization. The classes and libraries incorporate a custom, standalone facility for managing dynamic memory on the CPU and GPU, similar to the Thrust and RocThrust libraries provided by NVIDIA and AMD but lightweight and tuned to function as the building blocks of classes in molecular simulations. STORMM provides a means to develop objects in C++ and then manage equivalent memory layouts on both CPU and the GPU resources so that C++ prototyping can decrease time-to-solution with equivalent GPU kernels. In principle, STORMM is compatible with any of the major GPU computing libraries, but in development to date we have tailored kernels for NVIDIA’s CUDA language.

There is little conceptual difference between a multi-core CPU implementing vectorized instructions and a GPU. Both are built on the same lithographic processes, but in the former more of the silicon is devoted to fast “on chip” data storage and in the latter more of the silicon performs arithmetic. The key to effective GPU utilization is memory management and organization, which STORMM accomplishes through C++ classes that shed their formalism to communicate with CUDA kernels through a basic C language interface. In what follows, we will discuss design features of the STORMM libraries, its techniques for fixed-precision representations critical to parallel computing, methods for creating scalable work units in chemical simulations, and approximations for the most intensive molecular mechanics calculations. STORMM overcomes many scaling problems in GPUs by combining many small problems into one workload that can engage all the compute resources and optimize the communication bandwidth required by familiar, memory-throttled problems. We demonstrate how the STORMM algorithms turn force field valence term evaluation and particle-mesh charge density spreading into compute-bound processes. To close, we present performance data for parallel solution of small molecule energy minimization and dynamics problems in implicit solvent and discuss a path to implementing a complete molecular dynamics capability in condensed phase systems with periodic boundary conditions.

Psivant Therapeutics supports open science and releases STORMM as open-source software through the MIT license.

## 2 Guiding Principles

### 2.1 The durability of general-purpose serial programming and special-purpose vector programming

A program running in a single thread will execute a sequence of commands without consideration to other participating threads. This earliest model of programming in still useful for prototyping, as it translates into the simplest code and can be used in performant applications so long as the single CPU thread is managing a process that is yet slower, e.g. reading a file from disk and organizing its contents into data structures. An early model of parallelism from the 1990s, based on a message-passing interface (MPI) between threads that maintain separate memory allocations, is organizing many concurrent programs to run side-by-side. It arose in an era of cluster computing where each thread was running on its own physical machine. The advent of multicore processors with shared memory resources in the following decade and, on a grander scale, non-uniform memory access (NUMA) supercomputer systems, enabled a different model of parallelism, one in which multiple threads operate on the same memory addresses. This can, in principle, be more efficient (not least because each thread does not need to maintain its own image of the problem). The NUMA model is what GPUs have taken to an operational end point: a dense collection of vectorized processors controlled by a single scheduler to operate on a common pool of data, contained in a device that can be held in one hand. Instead of additional libraries that enhance single-threaded programs, GPU hardware has also arrived with specialized programming languages and compilers to broaden its application. With the end points established, the key is to develop mechanisms that improve the connectivity between the new and old computing models.

### 2.2 Unlimited topologies and coordinate sets in a single runtime instance

Unlike typical MD packages, STORMM begins with the premise that topologies and coordinates should be self-contained objects that can be copied, moved, arrayed, and reassigned at will. Furthermore, STORMM develops specialized coordinate and topology *syntheses* to concatenate many such objects into linear, indexed arrays of like quantities (e.g. Cartesian *y* coordinates of particles for all systems or the first atom targets of all distance restraints). In this manner, STORMM can pack a series of topologies and coordinate sets behind a single collection of pointers residing in the GPU’s constants cache, conserving critical register space in GPU kernels. The practical limit of this packing is the amount of global memory available in the GPU; however STORMM’s data structures and algorithms are designed to conserve RAM, whether on the CPU host or the GPU device. Modern desktop workstations and rack servers often have 64GB or more of main memory, whereas GPUs are available with more than 24GB of their own RAM (although still expensive as of the time of this publication). The tests that follow will show STORMM running collections of many individual systems which, combined, total over 2.6 million atoms. For some applications, a GPU with 10GB of memory can run problems comprising 7.5 million atoms. Planned improvements to STORMM include leveraging the existing coordinate copy and swap features to stage a “reservoir” of coordinate replicas in host RAM which can be cycled through a smaller array of system images allocated on the GPU, extending the number of atoms that STORMM might process with a single GPU into the tens of millions.

### 2.3 CPU prototyping, GPU execution

Of all the programming models, it is simplest to debug a serial program running in a single thread, both for the simplicity of the logic and the breadth of tools available, in particular the memory checking program valgrind[13]. STORMM develops classes based on a memory model with equivalent arrays in the CPU and the GPU random-access memory (RAM), as will be explained in detail later (see Figure 1). While it is not obligatory that the arrays contain the same information, for the purposes of prototyping a C++ function may be written to look at the same information in the same layout as the cognate GPU kernel. The C++ code can be subjected to unit testing and memory checking before serving as a model for the GPU kernel to emulate, leading to more rapid development and extending the complexity that would be possible in GPU implementations.

**Figure 1:**
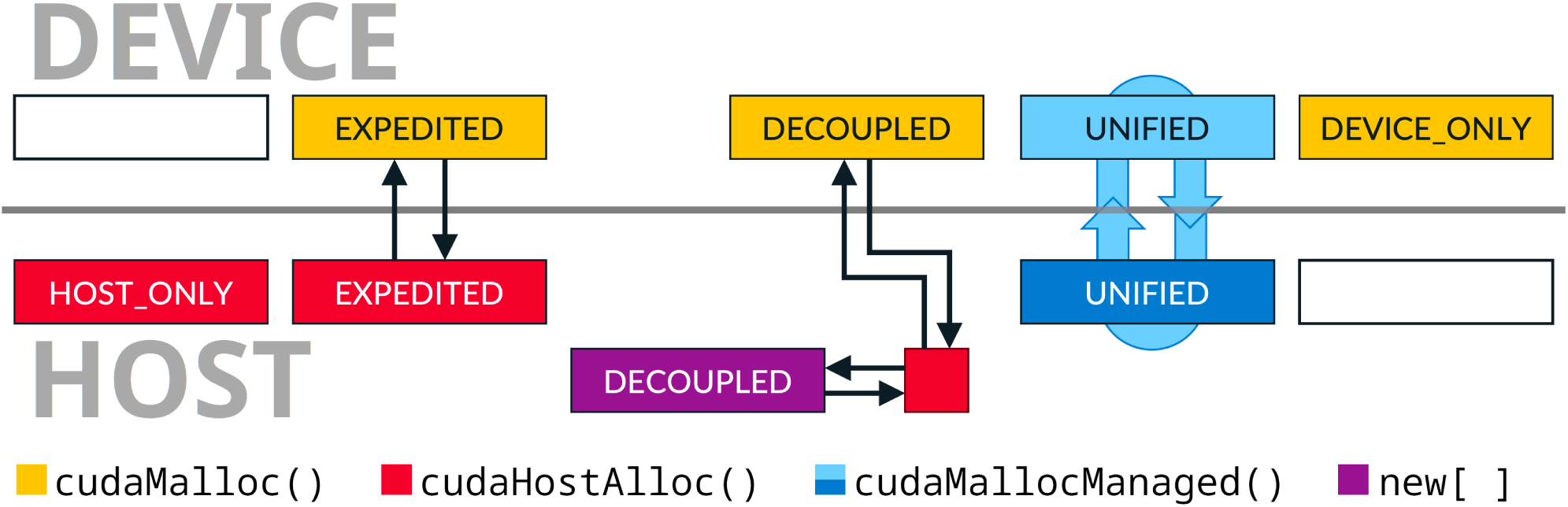
Allocation modes and memory transfer protocols in the Hybrid<T> class. The CUDA or C++ function responsible for allocating memory on the CPU host or GPU device is colored as shown in the legend. White boxes indicate no array on one side of the partition. For “DECOUPLED” allocations, upload and download of memory within an object of the class are mediated by a private buffer of page-locked memory managed by the CUDA driver itself. For “UNIFIED” allocations, the CUDA driver’s page migration engine manages automatic and obligatory synchronization. Otherwise, direct transfers are possible. While not supported by class methods, free functions may accept pointers and appropriate size bounds to transfer memory between separate Hybrid<T> objects.

In any GPU program, a great deal of the code is compiled on the C++ side to stage information for GPU operations. Aside from establishing a formal memory model between the two computing layers, STORMM leverages C++ code to develop refined work units with dense, often bit-packed instructions that minimize the necessary bandwidth between the GPU’s RAM and its streaming multiprocessors needed to direct computation. In the most optimized inner loops, it can be a helpful philosophy to write code such that “the GPU does not know what it is doing.” Rather, its threads are reading instructions that, in the space of 64 bits, say things like “access atoms 5, 89, 91, and 94 from the local storage array, compute the torsion angle, and scale the effect by parameter 571 before adding a non-bonded interaction between the first and last atoms screened by factor 23.” It is possible to trace where atoms 5, 89, 91, and 94 of the local cache came from by looking up the indices in a separate array, but debugging such code for all GPU threads would be intractable if there were not a corresponding function in C++ coupled with sufficient annotation. Also essential is extensive unit testing in the C++ code that writes such instructions. STORMM fosters a programming model that enables developers to write CPU code that plans GPU actions to reduce __global__ memory bandwidth to a degree that much more of the traditional computational chemistry problems become compute-bound, a sort of performance bedrock. This implies a degree of complexity and obfuscation of the information that would be right at hand in a serial C++ program written twenty years ago, but the complexity is encapsulated in a particular routine rather than forcing developers to offset memory bandwidth limitations with high-level techniques such as interleaving communication and compute-intensive processes, which can be sensitive to problem sizes and new hardware developments.

### 2.4 Mission Statement

STORMM seeks to internalize complexity while presenting outward simplicity and versatility. Compute-bound processes are valued above all else, as these are simplest to incorporate into any sequential workflow without adding pressure on developers to make special accommodations. While STORMM implementations are designed to be compatible with many programming methods, no assumptions are made to the effect that the implementations will be used in the context of high-level partitions of the GPU resources such as CUDA streams in order to optimize performance. Each specific process, such as particle density mapping to a grid or pairwise interactions between all atoms in a neighbor list, is written to be self-contained and separable from other stages of a workflow. This is perhaps best showcased by the fact that many STORMM classes and GPU kernels have specific benchmarking programs, distributed with the software, which can be used to reproduce a great deal of the results presented in this study.

STORMM routines are designed to exhibit elastic scaling, spreading out small problems to the greatest degree possible and condensing larger workloads to maximize throughput, gearing down as if it were “an automatic transmission.” Furthermore, most of the core objects are either templated or outfitted to support multiple precision modes, giving the developer a choice rather than imposing considerations about data types or numerical stability. To this end, many of STORMM’s innermost routines are rich in pre-processor directives that compile multiple branches of each kernel. Many existing codes provide a handful of precision modes to choose from, but STORMM compiles all variants of each routine into the same library in a way that provides users with finegrained control of phase space representations, force accumulation, and arithmetic in different stages of molecular mechanics calculations. The precision models of different processes are “mix and match,” e.g. valence and particle-particle interactions can be computed in float32_t (“single”) precision while particle-mesh interactions and FFTs are handled in float64_t (“double”) precision. Wrappers underneath the primary API will analyze the GPU to optimize workloads and select the appropriate kernel while various internal checks ensure that each participating object is constructed in a way that will support the requested calculation.

## 3 Core Capabilities

Pursuant to the design goals of interchangeable components, bandwidth-bound (rather than latency-bound) communication between host/device memory, CPU prototyping with GPU implementation, and CPU organization with GPU execution, STORMM builds most of its classes as compositions of a single, versatile array: the templated Hybrid. Within each Hybrid object there are two pointers, one to memory on the CPU host and the other to memory on the GPU device, and methods for transferring some or all of the memory between them.

Building on this fundamental mechanic, STORMM classes are written with methods, often named data() and taking an input argument to indicate whether the information on the host or device is of interest, to get a plain C-language struct with pointers, critical sizing constants, and possibly other pre-calculated values essential to the class’s operation. Switching modes from the C++ accessor / setter paradigm to a C-like manner of manipulating struct objects is at the heart of STORMM’s interface between C++ and high-performance computing (HPC) implementations, and STORMM adds variations on the theme where appropriate, in particular with templated objects. To further facilitate integration of the host and device code, STORMM offers a number of built-in facilities for unit testing, user interface design, and data display so that developers can build and test new programs with relatively simple APIs to leverage the powerful underlying features for mass molecular mechanics calculations.

### 3.1 The Hybrid Class

Since C++11, the preferred dynamic memory class in C++ has become the Standard Template Library vector, std::vector<T>, where T is the data type. In STORMM, the fundamental dynamic memory class is the Hybrid<T> array. The class object can be resized, moved, and copied as needed, and shares a handful of incrementing features inspired by std::vector<T>. One can be created from the other: Hybrid<T> has a constructor based on std::vector<T> as well as accessors to emit std::vector<T> based on a Hybrid<T> object’s data, whether on the CPU host or on the GPU device (individual elements can also be returned in the base data type, but doing this for data on the device would be an extreme example of latency-bound communication). The most common HPC languages, CUDA and HIP, each support their own wrapper library for the C++ Standard Template Library (Thrust and rocThrust, respectively). The Hybrid object is not nearly as complete, but offers some of the most essential functionality in a lightweight, built-in class that automatically binds arrays on the host and device. In most situations, declaring a Hybrid object is equivalent to declaring both a Thrust host vector and a device vector of equal size, with member functions for transmitting some or all of the data between the two.

While Hybrid<T> is designed to be a general-purpose array mechanism, it supports only plain data types and certain HPC tuples, e.g. float4, whereas std::vector<T> can be a container for C++ strings, user-defined types, even other std::vector<T> objects. Data types with their own methods tend to be harder for an HPC framework to utilize, although support for C++-like features in CUDA and other languages is broadening.

The exact layout of memory within each Hybrid<T> object is controlled by a C++ enumerator and fixed upon construction: Figure 1 illustrates various options. If STORMM is compiled for CPU operation alone, the sole option is HOST ONLY, enabling any code written with Hybrid objects to compile even without GPU support. When STORMM is compiled with GPU mode enabled, most objects are created in the default EXPEDITED mode, which allocates page-locked memory on the host and an equivalent block of memory on the device. In this format, GPU resources can upload and download information between the two arrays directly, with no intermediary managed by the graphics driver. The only limit on the amount of page-locked memory is that it must reside in the machine’s RAM, i.e. cannot be included in disk swap, but this is not a practical impediment in most situations. Even in GPU mode, STORMM can create Hybrid<T> objects with memory allocated only on the host, which might not have their own device memory but are nonetheless accessible to cards capable of writing host memory (which includes all of NVIDIA and AMD’s modern lineup). Other modes for the Hybrid<T> object include memory allocated only on the device and (when compiling with NVIDIA’s CUDA) unified virtual memory, which enforces synchronization of the two arrays by the driver’s internal page migration engine. However, unified memory is sub-optimal in most cases, intended to support the assumption that certain information is consistent between the host and the device, yet still limited by the PCIE bus and enforcing that data be maintained in two places at once. Judicious use of the Hybrid<T> object and the framework it offers allows developers, for example, to launch a GPU kernel to perform intense calculations on chemical structures in the memory on the device while staging new inputs in the corresponding space on the host, such that at the conclusion of kernel launch the CPU will have completed preparation of a new batch of structures that is immediately ready for upload after the GPU has transmitted its results somewhere else, forming an assembly line of sorts with a smaller memory footprint and high GPU uptime.

The Hybrid<T> class has a number of unique features, as well. Foremost, Hybrid<T> arrays are created as POINTERs or ARRAYs, reflecting an explicit pointer-array dichotomy reminiscent of C programming. A POINTER-kind Hybrid<T> array must target an ARRAY-kind Hybrid<T> in order to receive or dispense information, but when it does it can still have set bounds which will be applied even if the underlying array has more memory. A POINTER-kind Hybrid<T> array cannot be set to have bounds that would exceed the underlying target. Given the relative cost of cudaMalloc(), the primary means of allocating memory for the Hybrid<T>’s GPU, it is reasonable to include features that would not fit in a lighter, more agile class. The POINTER feature is one means by which STORMM reduces communication latency: many classes have a few ARRAY objects and multiple POINTERs targeting them. The AtomGraph, STORMM’s molecular topology class, contains more than eighty POINTERS all targeting one of four ARRAYs. Uploading or downloading the AtomGraph’s ARRAYs incurs much less latency than handling each of its POINTERs in series. However, if bandwidth is limiting, each valid POINTER may copy its data between the host and device, in whole or in part as needed.

Each Hybrid<T> array includes a C-string label and falls under a global memory ledger which tracks the total amount of memory allocated and the number of times that each array has been re-allocated. The labels will be included in various error messages related to allocating, freeing, or accessing data within a Hybrid<T> array, offering valuable clues when problems arise. When debugging accesses to a POINTER-kind array, the memory management system can be used to verify that the underlying array has been allocated the same number of times was recorded by the POINTER-kind array when it was first set, a strong indicator of whether the pointer is still valid. Future improvements may make use of the memory ledger to automatically update POINTER-kind Hybrid objects when the underlying ARRAYs are moved or destroyed.

### 3.2 The Abstract System

The abstract system shared by most STORMM classes creates a C-like means of working with the underlying data, whether in C++ code on the host or HPC languages on the device. Abstracts are all structs with no private members, accessors, or other member functions aside from the constructors, and most have some or all members initialized as const. They can be posted to the global constants cache of an HPC code unit if they do not change often, or passed as kernel arguments at minimal cost in launch latency (one clock cycle of the HPC device per 4 bytes of the struct’s overall size), giving each kernel launch the intuitive appearance of a C/C++ function call.

Some classes with many member variables may emit multiple abstracts, each specialized for a particular purpose. The AtomGraph topology emits separate abstracts for valence interactions, non-bonded interactions, constraints, and virtual site manipulation, with minimal overlap in their contents. In some situations, a class will have methods for emitting abstracts specific for 32-bit or 64-bit floating point calculations, to conserve the HPC constants cache and reduce the latency of kernel launches and function calls. In general, mutable class objects will emit *writer* abstracts with non-const pointers into their internal arrays and const objects will emit *readers* by different overloads of their data method.

Most templated classes will emit abstracts in their native type. However, there is a barrier for communicating templated information between C++-compiled and HPC-compiled *executable* objects (various files often ending in “.o” produced by the compiler, distinct from *class* objects). The two compilers can often work together, but require that each implementation necessary for code they compile be available at compile time, which will not be the case for templated classes in the executable objects produced by the other compiler. To solve this limitation and continue applying C++ and HPC compilers to their respective implementation files, templated STORMM classes intended for HPC operation also emit “template-less” abstracts through special methods, where all template-dependent pointers are cast to void*∗* , the C language implementation of a generic pointer. One key requirement is that the abstract have only templated *pointers*, not templated constants such as T atom count with T changing identities from short to long int in different class objects. At the same time, templated free functions (using the overloaded name restoreType()) will accept one of the template-less abstracts and recast the void*∗* pointers to the developer-specified type. In situations where it is hard to anticipate what type will be coming and going, STORMM keeps an extensive list of the type ID numbers set by std::type index(typeid(T)).hash code(), and the appropriate 64-bit code for <T> is just a size t that can be transmitted alongside the template-less abstract. While the C++ and HPC compilers may not be able to assemble executables with each other’s templated functions, they can each compile the same code and header files for themselves, and thus know how to strip away or restore typed character to an abstract. The recommended approach is to cast away templated character on the C++ or HPC side, transmit the abstract to code built by the other compiler, and restore the templated type on the other side. This is another way in which the abstract system connects C++ and HPC implementations: the abstracts, containing only pointers, can pass through the C++:HPC barrier, while the original objects cannot have their internal arrays and very identities recast to <VOID>, even by deep copy.

In general, the data member of each class accepts an indicator as to whether the abstract should collect pointers valid for memory on the CPU host or on the GPU device. Most modern GPUs are able to take pointers to page-locked host memory and write directly to them. (This could be a common mistake among new developers, who may find their code running slow because they have unwittingly created an abstract for data on the host and submitted it to an HPC kernel.) However, for safety as well as complete coverage, STORMM classes that may be engineered to have the GPU write directly to host RAM will do so via the deviceViewToHostData() method, which constructs the abstract with pointers to host data guaranteed to be valid on the GPU device.

### 3.3 Built-in Unit Testing Functions

While multiple unit testing frameworks are available, the extra dependencies can complicate and slow down compilation. Instead, STORMM comes with its own lightweight library of unit testing features specialized for assertions and vectorized comparisons relevant to molecular simulations.

The library functions can evaluate the truth of a statement, evaluate equalities or inequalities according to a given tolerance, and replicate such assertions for all elements of two vectors. To facilitate updates to unit tests as methods change or if existing tests are found to be incorrect, there is a -snapshot command line argument for storing lengthy streams of numbers in a developer-specified file which becomes part of the distribution. Unit testing programs can read the file in order to compare the current code to the established result, or take the command line option -snapshot as an instruction to rewrite the file according to the current code’s output. As another fundamental capability, STORMM’s unit testing library has macros to wrap code that is expected to fail in C++ try / catch statements, permitting checks on input traps or other situations that are intended to raise exceptions.

STORMM’s framework will organize tests into categories according to developer specifications, so that testing programs can report successes and failures as part of broader groups. The unit testing library can also identify whether it is possible to run a test, skip it if required inputs are missing or inaccessible, and report that result along with other successes or failures.

### 3.4 User Interface Development

STORMM offers a facility for creating command line inputs to new programs and command files based on an enhanced Fortran &namelist syntax. In STORMM, namelist keyword / value pairs can be separated with or without commas and linking keywords to values with an = sign is likewise optional. At the developer’s discretion, keywords can be set to accept multiple entries, with the first entry either adding to an obligatory default value or replacing it. The input libraries keep records of whether each parameter retains a default setting, has been set by the user (even if the user’s input specified the default value), or is missing from the input (and lacking a default). In addition to the familiar real-valued, integer-valued, and string-value keywords, in STORMM &namelist keywords can also be structs with real-, integral-, and string-valued subkeys. The input libraries let the developer specify help messages for each new keyword, where to begin scanning a command file for a particular &namelist (and whether to wrap back to the beginning if it is not found), and provide methods for command line input to have a table of all keywords and descriptions for one of the program’s relevant &namelists displayed in the terminal.

### 3.5 Data Display Features

Because one of the underlying goals is to help developers gain more control over the numerical aspects of molecular simulations, STORMM has a number of features for tabulating results in human-readable format which are also amenable to popular matrix algebra packages such as Matlab or GNU Octave, Python tools such as NumPy and MatPlotLib, and the mesh format OpenDX. Where possible, STORMM attempts to make use of the output format’s comment syntax to label columns of tables and provide narrative for each input section, creating files that can be read like the output summaries of typical molecular simulations engines but also fed as scripts the preferred input package to display the data in graphics or have the data available for further manipulations.

### 3.6 Chemical Features Detection

While the RDKit software[14] provides excellent capabilities for exploring chemical space, it requires certain inputs that may be lacking in molecular dynamics force fields or topologies, such as chemical bond orders and formal charges. These two descriptors are best calculated in conjunction with one another. As a stepping stone to its planned integration of RDKit, STORMM offers an implementation of the Indigo scoring function.[15]

The Indigo developers’ original approach applied the A* algorithm[16] with graph libraries written in Python and Boost C++. In STORMM, the modular implementation builds on the software’s built-in topology and coordinate objects to create instances of the ChemicalFeatures class. Rather than A* or generalized methods in graph theory, STORMM reduces the complexity of the problem by first identifying tetrahedral carbon or nitrogen atoms: the octet rule posited by the Indigo scoring function implies that these atoms must have single bonds connecting them to each of their substituent atoms. This and other logical deductions in the STORMM implementation subdivide the atoms into smaller groups, allowing bond orders and formal charges for even large drug-like molecules to be resolved in milliseconds. Lewis structures of entire proteins can be drawn in tens of milliseconds, on the order of the time it takes to read their respective topology files from disk. There remain a handful of cases prone to combinatorial explosion, e.g. a graphene sheet, but these can be solved with additional improvements to the divide-and-conquer strategy.

Furthermore, the implementation in STORMM identifies all iso-energetic solutions of system under the Indigo scoring function and stores real-valued bond orders and partial formal charges, e.g. bond orders of 1.5 around a benzene ring and partial charges of 0.5 atomic units for both carboxylate oxygen atoms. This allows STORMM to store resonance states and correctly identify some atoms as having the same “chemical rank” where RDKit fails to recognize their equivalence.

The ChemicalFeatures class plays a central role in resolving atom mask selections in STORMM, as well as identifying ring systems and rotatable bonds for conformer generation. It also enables STORMM to read a molecular mechanics topology, containing elements and connectivity, and produce a complete Biovia MDL MOL file (Dassault Systemes, Paris, France), the core of the industry-standard .sdf format. In the future, it will be critical for the interface between STORMM and RDKit, and could be instrumental in identifying force field terms needed to develop molecular models in pursuit of high-throughput three dimensional model rendering based on generative modeling in chemical space.[17]

### 3.7 Supported Chemical Models

STORMM is intended to handle a breadth of molecular models. While electronic polarization and potentials based on deep learning are outside the present scope of the project, there is already support for all common AMBER and CHARMM force field terms, including harmonic bonds and bond angle terms, Urey-Bradley potentials, Fourier cosine series dihedral and improper dihedral potentials, harmonic improper dihedrals, Correction Map (CMAP) terms for two dihedral terms sharing three atoms in common, and flat-bottom harmonic restraints (further description in Section 4.4). In terms of non-bonded potentials, STORMM implements Lennard-Jones potentials and will identify geometric, Lorentz-Berthelot, or pair-specific Lennard-Jones combining rules (the analysis is performed on any input topology and all rules run under a common implementation). Cutoff-based Lennard-Jones interactions are supported, and a dispersion calculation based on the Particle Mesh Ewald method[18] (see Section 5.4) is planned. For more elaborate and accurate monopole distributions, STORMM implements all virtual site forms found in the GROMACS[19] palette of massless particles. Implicit solvent calculations on small molecules or small proteins are possible with any of the AMBER software’s models.[20, 21, 22]

## 4 Coordinate and Topology Compilation

### 4.1 Specialized Coordinate Representations

STORMM has five unique coordinate objects, including an object that only stores Cartesian *x*, *y*, and *z* particle positions with box dimensions (if applicable), a simple representation of particle positions, velocities and forces that is suitable for managing molecular dynamics of a single system, and an extensible array of coordinates for a series of snapshots from a single system. While all box dimensions are held in float64_t (“double” in C++), the data types of coordinates in each object can vary based on its purpose.

Traffic between all coordinate objects is managed by a versatile API, which includes coordCopy() to copy coordinates between any two coordinate objects, conserving as much bitwise detail as possible and altering only memory on the CPU host or GPU device as specified in the call, e.g. coordinates may be taken directly from the GPU-based memory of object A to the CPU host memory of object B, without altering the host memory content of A or the GPU-based content of B. For like coordinate objects each containing multiple structures, the coordSwap() function can swap one or more systems from one to the other, to and from memory on either the host or device, with a single kernel launch to maximize communication bandwidth. Table 1 lists specifications of the available classes while Table 2 indicates their possible applications.

**Table 1:**
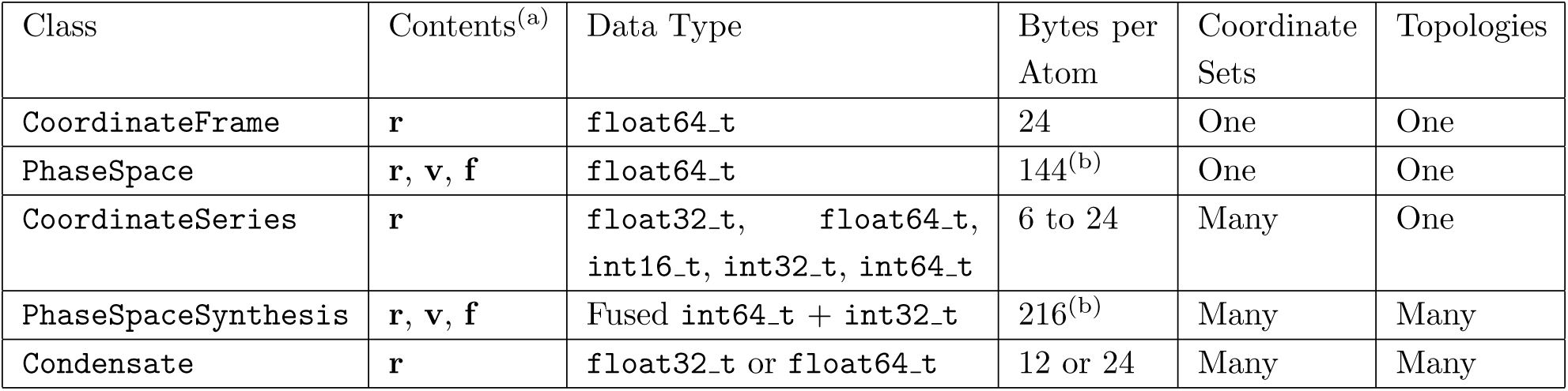
Properties of STORMM coordinate classes. **(a)** Cartesian data: **r** = particle positions, **v** = velocities, **f** = forces **(b)** The PhaseSpace and PhaseSpaceSynthesis objects have twice the expected size for their data types due to holding two copies of all coordinates: one for holding the current or reference state, the other for constructing the forthcoming state during dynamics or energy minimization.

**Table 2:**
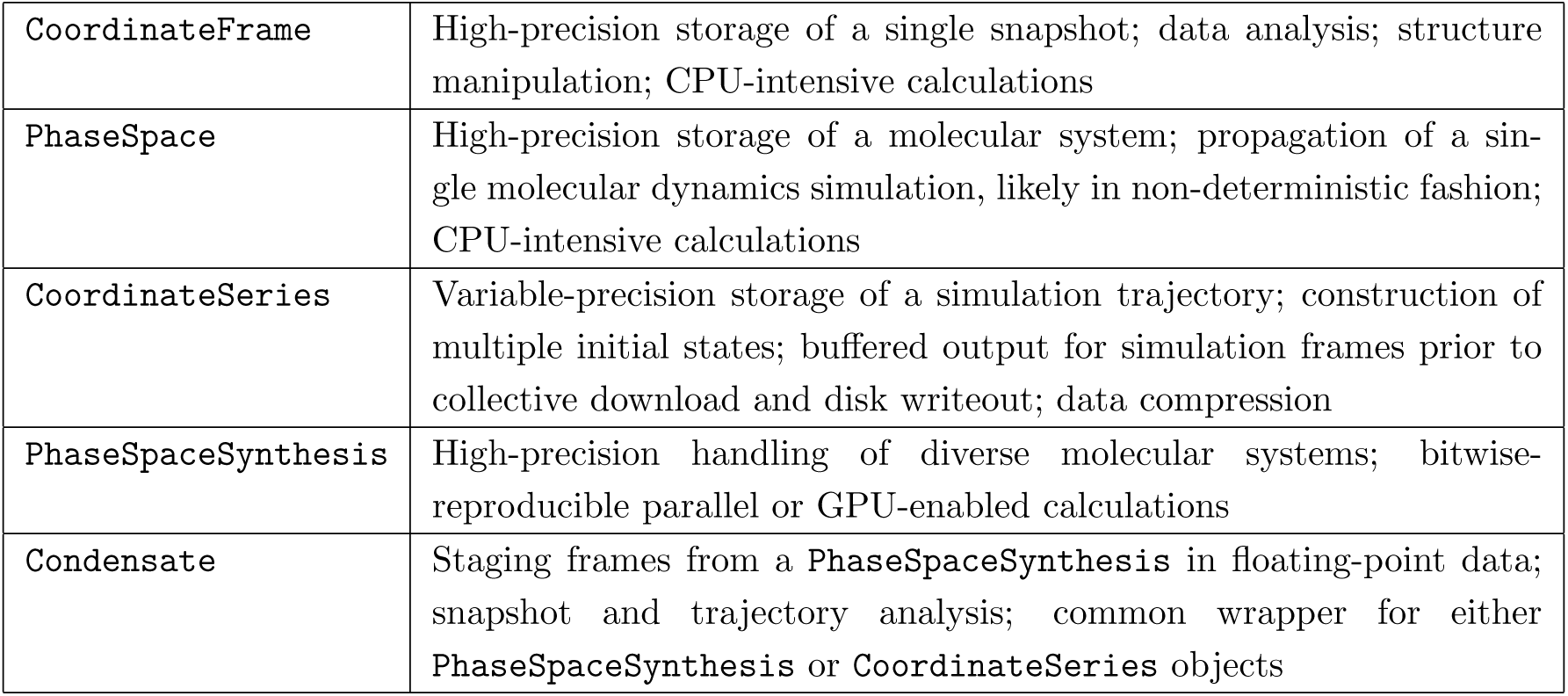
Uses of STORMM coordinate classes.

### 4.2 Many Coordinate Systems in One: The PhaseSpaceSynthesis

The most complex, memory-intensive, and secure coordinate object is the PhaseSpaceSynthesis. Like the primitive PhaseSpace object, it stores positions, velocities, and forces, but it concatenates like pieces of the information for each system, e.g. the particle positions along the Cartesian *x* axis, and stores arrays of limits to track the demarcations between each system. Designed for GPU computing with bitwise reproducibility, the PhaseSpaceSynthesis stores all coordinates in fixed-precision format with combined int64 t and int32 t data. The length of the fixed precision representation for each coordinate type may be specified with the creation of the object, but any aspect of the phase space and forces for each system may be specified to a precision of up to one part in 2^72^ of the internal units of Angstrom (Å), Åper femtosecond, and kcal/mol-Å. For convenience, the PhaseSpaceSynthesis object also stores pointers to the original topologies, the AtomGraph class in STORMM, for each of the systems in its collection. Various constructors are available to create the PhaseSpaceSynthesis from an array of existing PhaseSpace objects, replication, and other indexing or tiling methods. There is no restriction on disparate system sizes present in any one PhaseSpaceSynthesis, but there is a restriction that either all systems have periodic boundary conditions or that no system has any boundary conditions.

### 4.3 Many Topologies in One: The AtomGraphSynthesis

Just as the PhaseSpaceSynthesis compiles multiple coordinate systems by concatenating their component arrays, the topology synthesis, AtomGraphSynthesis, compiles multiple AtomGraph topologies describing different systems into a single object, concatenating arrays such as atomic partial charges, Lennard-Jones tables, and bond term atom indices in the process. In the interest of conserving local cache space, STORMM will make consensus tables for valence terms such as dihedral parameters so that all systems can work from common information which is likely to be relevant even if the molecular systems themselves differ. However, Lennard-Jones pairwise interaction matrices, which would grow as the square of the number of unique atom types, are only used for replicas of the same system: otherwise, the Lennard-Jones pairwise matrices for each unique topology are concatenated into a common array. Tracking systems and interactions in the AtomGraphSynthesis is a complicated task, but this is handled by single-threaded C++ code for easy debugging. The C++ code reconfigures atom indices and condenses parameter tables as needed, then constructs work units with compact instruction sets encoded in bit strings so that GPU kernels can perform cycles of work, one of which is detailed in Section 5.3.

The AtomGraphSynthesis emits several abstracts for handling valence interactions, non-bonded interactions, virtual site manipulation, and other calculations depending on the work of a particular kernel.

### 4.4 Supplemental Restraints: the RestraintApparatus

While a complete chemical model can express interactions among particles in the form of the STORMM AtomGraph and AtomGraphSynthesis, computational experiments often rely on controlled perturbations that can be subtracted from the simulation in post-analysis. STORMM offers the RestraintApparatus to flatbottom, harmonic restraints similar to AMBER’s NMR potentials for a chemical system. Formally, the information in restraints is separate from the topology (the unitary AtomGraph), but it can be incorporated into the AtomGraphSynthesis.

Due to the difficulty of enumerating restraints over the large numbers of systems that STORMM is designed to handle, users can specify groups of restraints with a single command, e.g. “all heavy atom dihedrals in this atom selection mask” or “distance restraints that penalize intra-molecular hydrogen bonding.” Restraints can also be applied to a particular structure, a labeled group of structures, or all systems in the runtime instance.

## 5 Unique Numerical Implementations

### 5.1 Split Fixed-Precision Representations with atomicAdd()

One of the most significant numerical innovations in STORMM is the use of multiple accumulators to store and accumulate fixed-precision representations of numbers, in multiple integers. Fixed-precision representations work by multiplying floating point numbers by some power of 2, common choices ranging from 20 to 40, and recasting them as integers to accumulate the result. The rounding technique is a matter of choice, although the goal is often to provide enough information in the fixed-precision number to capture all information in the mantissa of most floating-point contributions. Fixed-precision accumulation can be used to extend the precision of calculations taking place within well-bounded ranges, when catastrophic cancellation is a problem. More often, it is used to achieve bitwise reproducible results in parallel programming, where sequential uncertainty can combine with the non-associativity of floating point addition.

In many situations, it is sufficient to accumulate quantities such as the representation of a force with a fixed-precision scaling of perhaps 2^24^, such that most accumulations will not go outside the range [*−*2^31^, 2^31^). However, if an accumulation does stray from this range, even on occasion, the result can be catastrophic: in the case of forces, a large negative force flips to become a large positive force. The solution is to make the fixed-precision representation able to accommodate numbers so large that the calculation, e.g. force in a molecular dynamics simulation, would itself be broken if a legitimate value of such magnitude were to appear. The insurance policy can be expensive: if 32 bit integers are sufficient to hold 99.99% of the accumulated forces, then a 64-bit representation which can hold 100% of the results, while necessary to ensure a stable simulation, is transmitting twice the useful information in all but one case out of 10,000.

A *split* fixed precision representation makes the insurance policy more adaptable: accumulation can go on in the “primary” accumulator, a 32-bit accumulator in the example above, while holding a second 32-bit accumulator in reserve in case of emergencies. Two conditions must be identified: first, when a specific contribution or initialization would overflow the primary accumulator, and second, when an incremental accumulation out of a series causes the primary accumulator to overflow. In each case, the overflow of the primary accumulator must be contributed to the secondary accumulator. STORMM implements the scheme illustrated in Figure 2, combining two 32-bit signed integers to support a 63 bit range for accumulating results from 32-bit floating point math or one 64-bit primary accumulator and a 32-bit overflow accumulator to support a 95 bit range for accumulating results from 64-bit floating point math.

**Figure 2:**
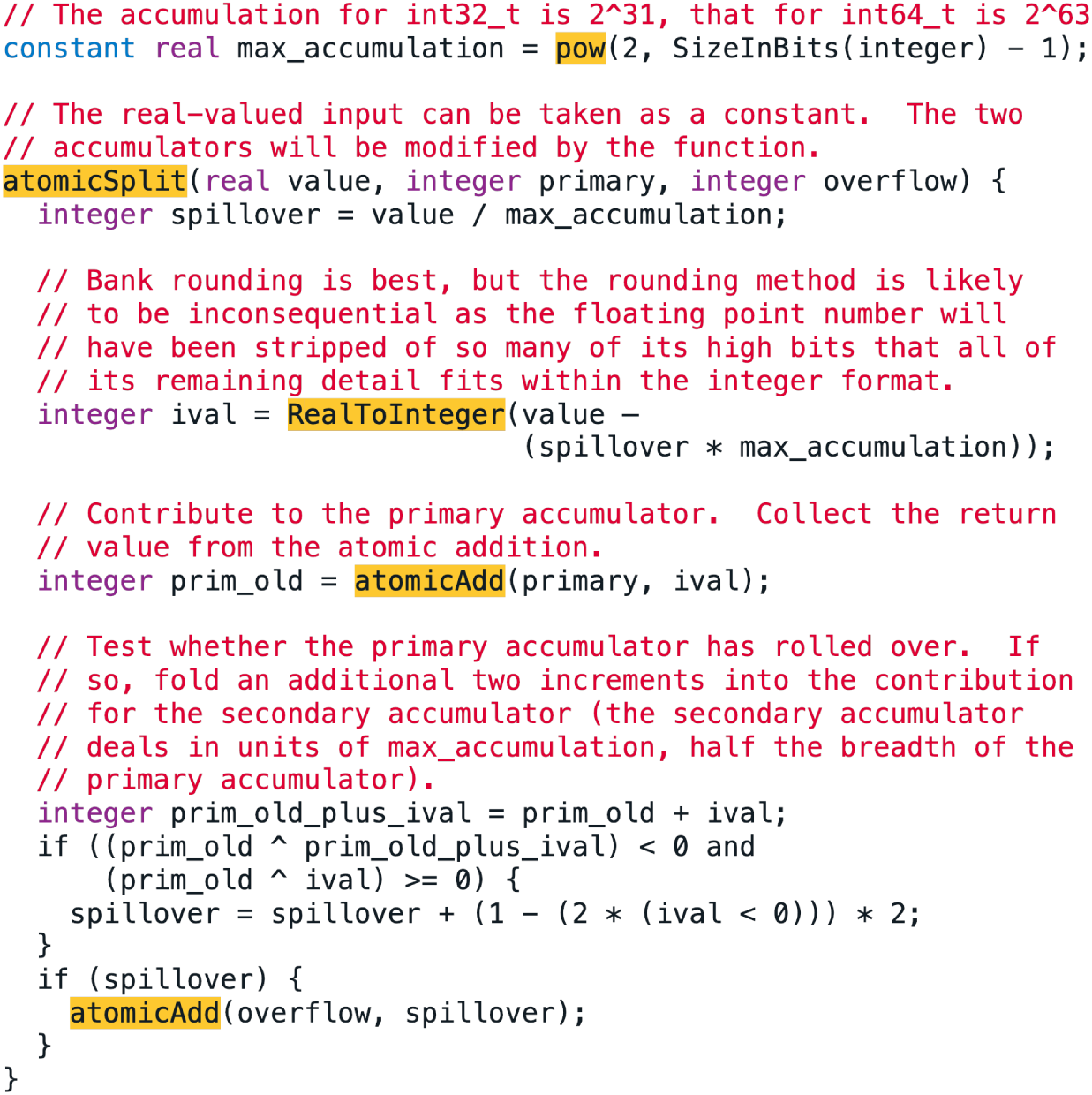
Pseudo-code for split fixed-precision accumulation. Function names are highlighted in gold, variable types written in purple. Comments appear in red. The carat (^) indicates bitwise exclusive OR (XOR).

As split accumulation is used throughout the STORMM code base, the performance enhancements it confers will be presented in context. For reference, a handful of benchmarking kernels were created that perform a minimal amount of math to vary float 32t counters with a median value of 45.0 *±* 17.0, then scale them by different fixed-precision factors and accumulate them using atomic operations. In these cases, the vast majority of the work lies in the atomic accumulation and, depending on the depth of the fixed precision accumulation, the primary accumulator overflows with different probabilities, triggering the secondary atomic operation. Figure 3 shows the significant advantages that dual int32 t accumulators can have over a single uint64 t, particularly on commodity cards when the accumulators reside in a __shared__ memory partition of the chip’s local cache.

**Figure 3:**
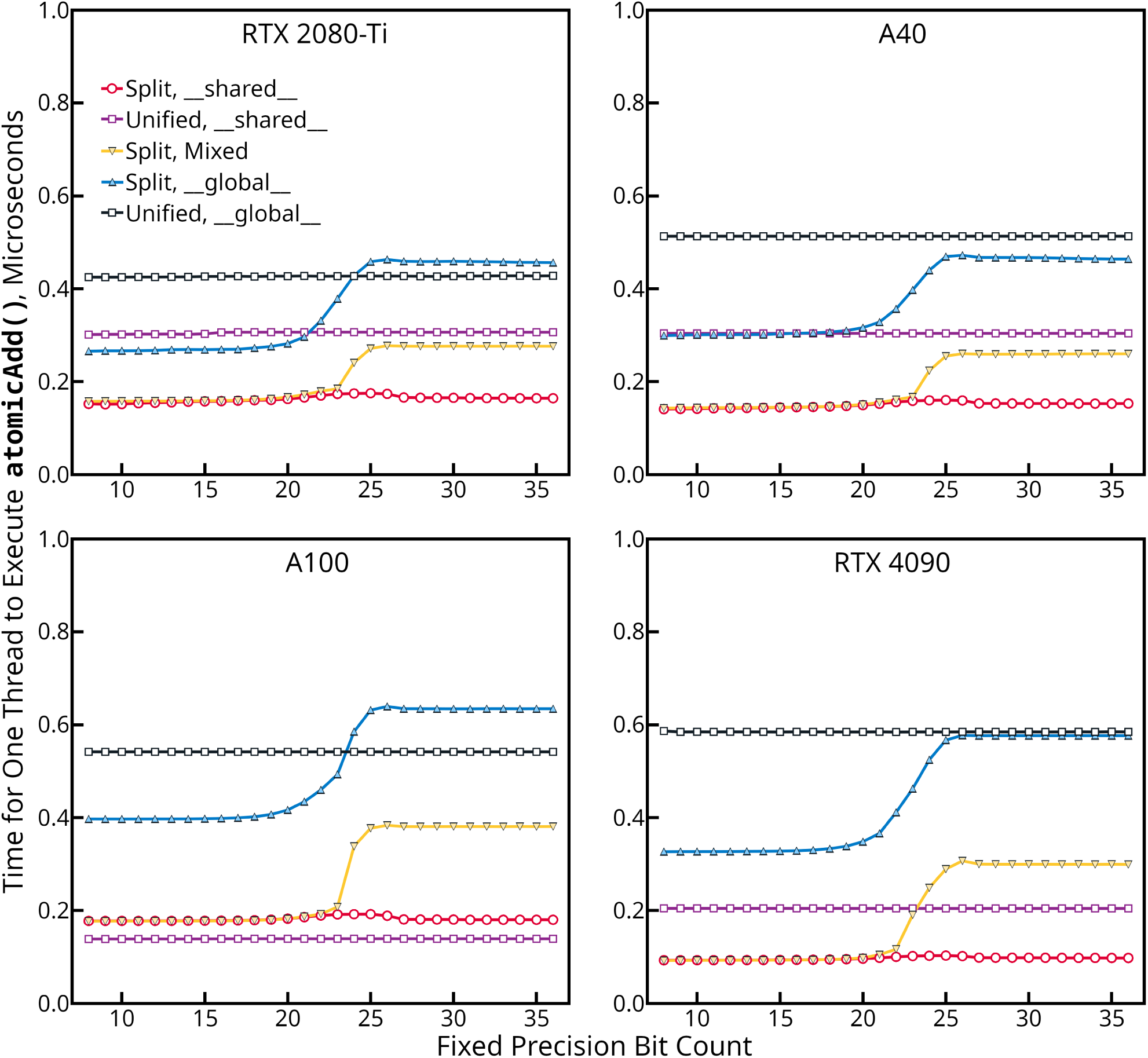
Timings for coalesced, non-competing atomics. In each operation, a float32_t number is scaled by a power of two indicated by the number of fixed precision bits, converted to integer format, and added to accumulators within the local or global cache. The kernel launch grid involved 1024 threads per streaming multiprocessor on each GPU. The overall throughput of any card is a function of the speed with which one thread can execute each atomic and the total number of threads active across all streaming multiprocessors. The inset legend in the upper left panel applies to all panels.

When the sum rarely spills into the secondary accumulator, the pair of int32 t accumulators can work up to twice as fast as a unified 64-bit accumulator depending on the card model. However, for the professional-grade A100 GPUs the performance of 64-bit accumulation in __shared__ resources appears to be as fast as solitary 32-bit accumulation, supplanting split accumulation by a small amount. On all cards, the cost of each atomic reaches a plateau as the accumulations approach 2^31^, the maximum amount that the primary accumulator can accept in a single operation and also the most likely to trigger incremental overflow. The split accumulation carries an advantage over direct 64-bit accumulation in __global__ memory when the secondary accumulators reside there (atomic operations to global memory addresses take place in the global L2 cache), and even, in the case of the A40 GPU, when both accumulators reside there. Keeping the primary accumulator in __shared__ while storing the secondary in __global__ is a way to conserve the most valuable chip cache space while still protecting against catastrophic overflow. Of course, the split accumulators can be used to carry out fixed-precision calculations on cards that do not support 64-bit atomic operations, for which the OpenMM program has adopted a similar method.[23]

Some notable properties of the GPUs presented in Figure 3 are listed in Table 3. While the speed of split atomics to __shared__ appears to be a function of the cards’ clock speeds, the trend with more impact may be the steady decline in the speed of 32-bit as well as 64-bit atomicAdd() instructions to __global__ memory addresses. In a given amount of time, the RTX 4090 can issue more instructions to L2 than an RTX 2080-Ti because the 4090 has almost twice as many streaming multiprocessors, but the instruction bandwidth falls relative to the computing capacity from one generation to the next. The benchmarking program used to collect the data presented in Figure 3 is distributed with the STORMM code base, as are other small programs used to carry out most of the studies in this paper.

**Table 3:** Critical parameters for NVIDIA GPUs used in this study. **(a)** The compute capability of the architecture, as codified by NVIDIA **(b)** Streaming multiprocessors (SMs) are contained compute units within the graphics card. Each SM has its own L1 cache resource and arithmetic units. **(c)** The cards’ base / boost clock speeds are displayed **(d)** The total L1 capacity is taken as the sum of texture and __shared__ configurable resources.

| Card Line | CC<sup>(a)</sup> | SM<sup>(b)</sup><br>Count | Clock<br>Speed,<br>GHz<sup>(c)</sup> | Global<br>Memory,<br>GB | Global<br>Memory<br>Bandwidth,<br>GB/s | Total L2<br>Cache,<br>MB | Total L1<br>Cache per<br>SM<sup>(d)</sup>, kB | Power<br>Draw<br>(TDP) |
| --- | --- | --- | --- | --- | --- | --- | --- | --- |
| RTX 2080-Ti | 7.5 | 68 | 1.35 / 1.55 | 11 | 616 | 5.5 | 96 | 250W |
| A40 | 8.6 | 84 | 1.31 / 1.74 | 48 | 696 | 6 | 128 | 300W |
| A100 | 8.0 | 112 | 1.07 / 1.41 | 80 | 1935 | 80 | 192 | 300W |
| RTX 4090 | 8.9 | 128 | 2.23 / 2.52 | 24 | 1008 | 72 | 128 | 450W |

The precision model in STORMM is intended to be very flexible and, with sensible default parameters, in the user’s control. Rather than simply applying “single” or “double” precision variants of the program, users will be able to apply a &precision namelist to their molecular simulations to toggle between single and double precision in various aspects of the calculations and also control the fixed precision bits used to accumulate forces or represent the phase space. Continued development in STORMM will keep all options open in kernels that rely on a great deal of communication, with the goal of fulfilling any requested precision model in the most efficient and safe manner possible.

### 5.2 Logarithmic Indexing for Tabulated Functional Forms

In AMBER18[24], the pmemd.cuda code introduced a novel method for approximating a function of one variable and applied it to the particle-particle interaction of Essmann’s Smooth Particle Mesh Ewald method (PME).[18] PME is a special case of the particle-particle, particle-mesh (P3M) method where the splitting function that divides the interactions between a form which vanishes in real space and one which converges in Fourier space is inspired by a Gaussian and, hence, incorporates the error function. The real space interaction takes the form:

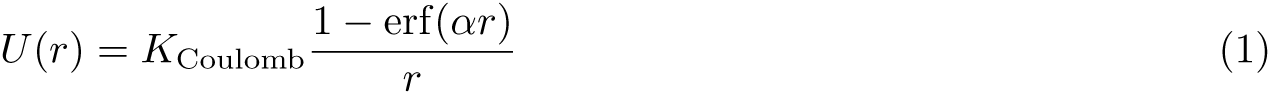

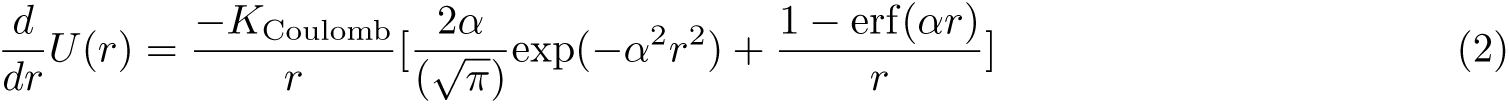

This form is laborious to compute, and numerically challenging although library functions for exponentiation and the complementary error function based on 32-bit arithmetic, expf() and erfcf(), are available on both CPU and GPU platforms. The logarithmic table index works well for functions with exponential decay, which are most challenging to approximate at short range and quite smooth at longer range. The method in pmemd.cuda interpreted the high 14 bits of the *squared* distance between two particles as an index into a table of spline functions. In effect, this consumes the exponent of the 32-bit floating point number (and the sign bit, which is always zero for a squared displacement) and the first five bits of the mantissa to index a table with 2^5^ = 32 spline segments on any interval [2*^n−^*^1^, 2*^n^*) for some power *n* represented in the format. The table density increases in proportion to 2 raised to (1*/r*^2^) as the squared displacement approaches zero, and while the overall size of a table with these parameters is 128kB only a narrow range of spline segments, covering a range of about [0.5, 100.0), would be used in a typical simulation for a total cache footprint of about 2.5kB. This is the basis of the method’s speed advantage: re-use of the cached coefficients many times over the course of the real-space particle-particle inner loops. While the original publication states “cubic spline” the form was, and remains:

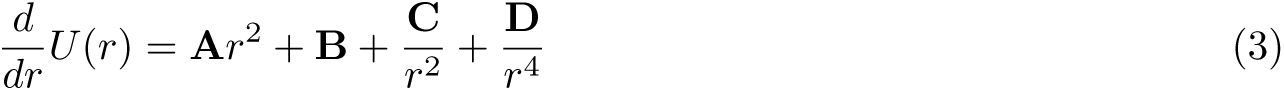

The series based on positive and negative powers of *r*^2^ uses quantities that were either already available or needed to be calculated for the Lennard-Jones evaluation. Because the spline coefficients **A**, **B**, **C**, and **D** can be fitted in 64-bit floating point arithmetic before converting to 32-bit floating point forms, the splines roll together a much larger number of arithmetic operations with greater numerical stability, a benefit which can be most apparent in the case of “excluded” interactions, where a raw Coulomb interaction is often subtracted from the particle-particle interaction to account for the fact that particles with covalent connections do not interact in their nearest image. In such a case, analytic calculation must contend with catastrophic cancellation in 32-bit precision. A simpler and more accurate way to handle the excluded interactions is to compute:

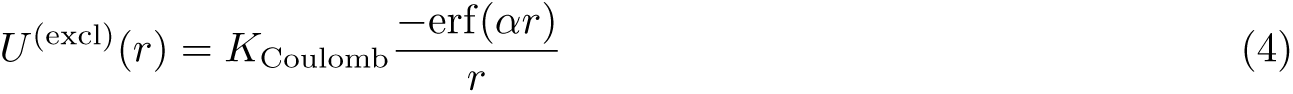

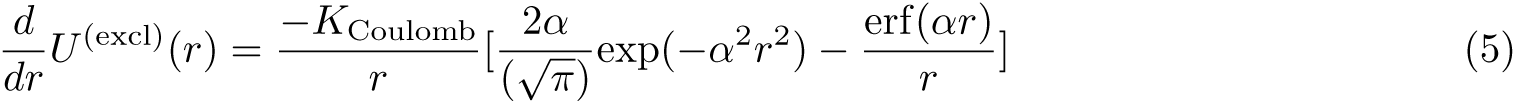

However, this is inefficient on GPUs because it bifurcates the parallel warp between two distinct and expensive calculations, one involving erfcf() and the other involving erff(). To avoid the bifurcation in favor of erff() produces the same catastrophic error cancellation, this time in the case of the more numerous non-excluded interactions. In contrast, splines offer rapid interpolation for either target function in the precision of 64-bit benchmark calculations, and clever indexing by introducing an offset in the table lookup based on whether the interaction is an exclusion can even avoid the warp bifurcation.

In STORMM, the approach is generalized in a first-class, templated C++ object for evaluating spline coefficients. Like other objects, the LogScaleSpline stores its critical coefficient tables in a Hybrid array to transmit them to the GPU, and emits an abstract with critical pointers, sizing constants, and bit masks specific to the implementation. The number of bits taken out of the mantissa in the corresponding floating point number is an input parameter to the object’s constructor, as is the form of the function to approximate, with options for the PME real space interaction potential and force, with or without exclusions, as well as other potentials and even a custom mode where the developer can provide a series of (argument, value) pairs to be translated into a spline table with logarithmic indexing. Furthermore, the numerical approach was revised to include a different polynomial. This was done using bitwise AND (&) to isolate the low bits of the mantissa, all bits of the original floating point number not used for indexing, left-shifting them to the front of the mantissa, then adding an exponent value of 127 (for 32-bit floating point numbers) or 1023 (for 64-bit floating point numbers) to form a new floating point number in the range [1, 2). The corresponding spline coefficients are fitted by solving:

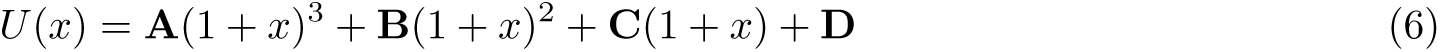

where *x* ranges from zero to one and is obtained from the progress of the spline indexing parameter, e.g. *r*^2^ where *r* is the argument to the distance-based function of interest. The STORMM implementation solves for the polynomial coefficients by sampling the target function at eight points spread evenly over the applicable spline interval, setting up an over-determined system of eight equations and four unknowns tractable to the method of least squares, which we implement by QR decomposition.

This framework combines the logarithmic indexing with a more traditional cubic spline, whose segments are shifted to work in a relative coordinate system with improved numerical stability. Another, more profound, interpretation of the strategy is that it recognizes that the high “indexing bits” of the original argument, hitherto the squared particle-particle displacement, are used to determine which element of the table to use, and cannot therefore contribute anything meaningful to the form evaluated using the chosen spline coefficients. The spline form is fitted to translate the nuances in the remainder of the original argument, the low bits of the mantissa, into a result. Jogging those “detail” bits forward to the front of the mantissa helps protect them from a degree of rounding error when computing the square or cube. The choice of exponent is arbitrary–equivalent results could be obtained with any other power of two if the cube stays within the floating point format range. The new implementation offers a more intuitive way to think about the information present in any given floating point number, and how this can be partitioned to useful effect. (Astute readers will note that the low bits of the mantissa could be jogged forward one more place to take advantage of the “hidden bit” in the IEEE floating point format. However, this complicates the implementation and would be more laborious to compute for a modest improvement in the accuracy.)

Error in estimates of the PME real space interaction for two protons is plotted in Figure 4, with and without the effect of a local, excluded interaction. The tests in panels A and B submit particular values of the squared displacement to be evaluated by the spline table or analytic computations, indicating the splines’ advantage in numerical precision. The picture is incomplete, however: in a real problem there are many ways in which two particles can be separated by a given distance, three Cartesian displacements in many combinations. In order to test the accuracy of either calculation in a more realistic setting, 32 trials were conducted for any given separation of particles, placing two points at random locations on the interval [*−*5, 5] along *x*, *y*, and *z*, then normalizing the distance between them to the desired value while keeping the midpoint constant. It was important to keep the particles near the origin of the coordinate axes, because calculations of any difference are more accurate for smaller numbers. (The forthcoming STORMM neighbor list decomposition follows a similar strategy of localizing the coordinate axes to conserve precision.) The coordinates were then converted to 32-bit floats so that the squared displacements and real space interactions calculated from this setup could be compared to the results of a framework using 64-bit floats throughout. In the 64-bit floating point framework, deviations in the magnitude of the interaction computed for different pairs of particles juxtaposed at the same distance are negligible, but panels C and D show that in the lower precision model it is the computation of the displacement which contributes the most error to the real space interaction between the particles. In turn, the lion’s share of this error is accrued through rounding when summing the squared displacements along three directions, which can have great variations in size. The improved accuracy of the splines is almost indiscernible against that level of noise, but it remains evident for excluded interactions. The fact that the splines are, by themselves, more accurate than the equivalent 32-bit computations is a compelling case for the approach, but in certain contexts, e.g. the presence of much less accurate (and obligatory) calculations and infrequent use preventing effective caching of the spline coefficients, direct 32-bit computations might be preferable.

**Figure 4:**
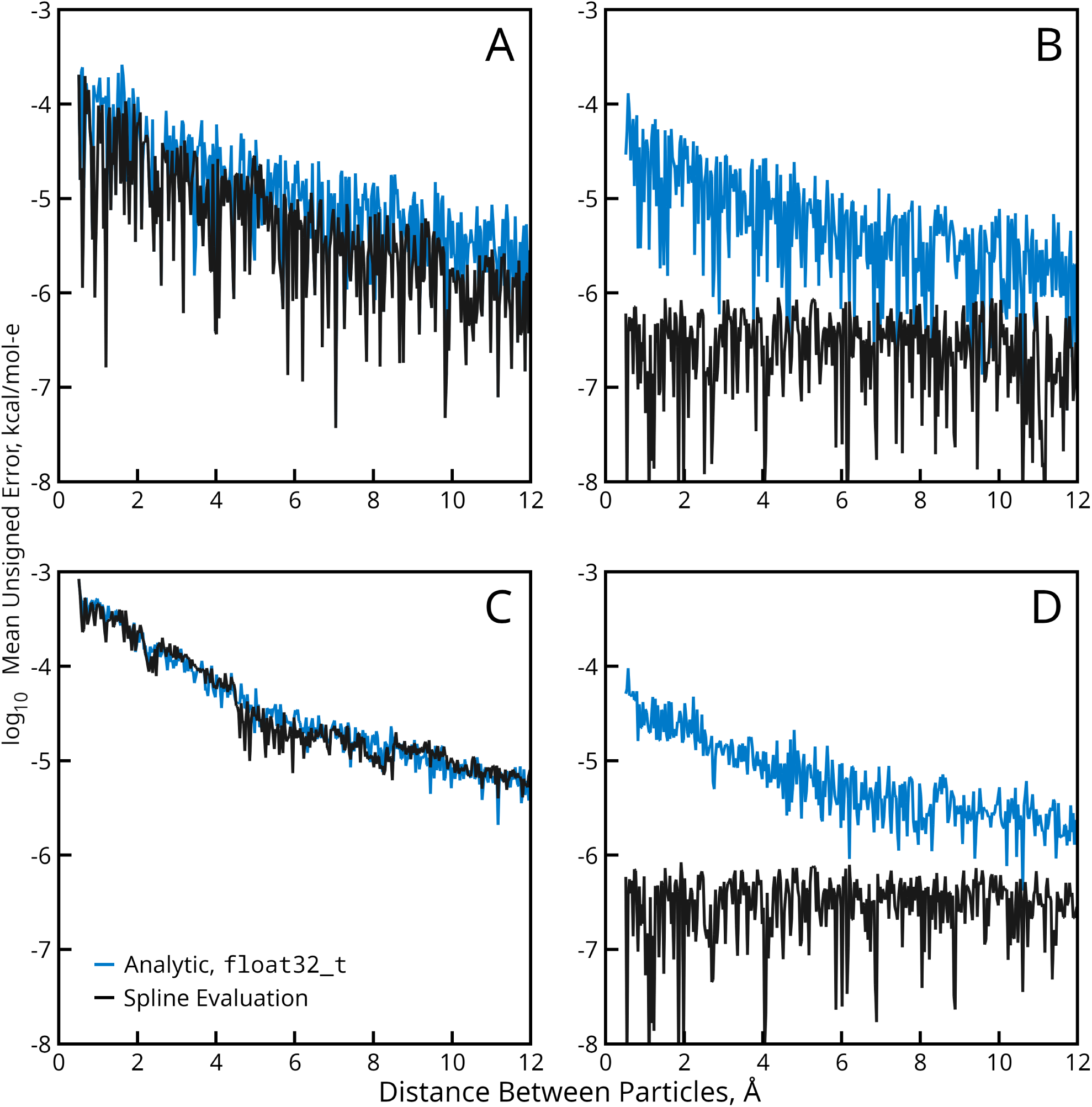
Error in calculation of the PME force between two particles with one atomic unit of charge each. Errors are computed relative to a standard computed in 64-bit floating-point arithmetic when the inter-particle distance is known to 64-bit precision. Panels A and B present error in the force with and without a pairwise exclusion of the Coulomb interaction in the home image, respectively. Panels C and D present the error inherent in such calculations when the intended inter-particle distance is known exactly but the displacement is computed in 32-bit floating-point precision. The inset legend of panel C applies to all plots: blue for direct computation and black for spline-based interpolation.

Table 4 demonstrates the effect of changing the number of bits taken out of the mantissa to serve in the exponent, and of indexing the table by the squared displacement or the absolute magnitude of the displacement. The results represent a weighted average over the applicable spline range, increasing as the square of the inter-particle distance to reflect the likelihood of any given range being sampled in a simulation. For comparison, STORMM also implements the older AMBER method of building the splines based on equation 3. The choice of devoting five bits out of the mantissa in AMBER arose from trying sets of 16, 32, and 64 spline segments per increment of the exponent, and finding that five bits (32 segments) was the best balance of accuracy and cache performance. In fact, for the old AMBER formulation there is hardly any difference among these choices. Table 4 shows that the new polynomial formulation is overall more accurate, and makes table indexing by the target function’s argument competitive with table indexing by the square. Even for the improved polynomial formulation, the accuracy hits a wall after five bits, and the discussion above provides a foundation for understanding why: in a 32-bit float with a mantissa of 23 bits, every bit used to index into the table is one that cannot be used to provide detail in the number evaluated by the spline basis functions. Doubling the number of splines comes at the expense of halving the accuracy in the value of *x* in Equation 6, and the leading bits of the mantissa are, at best, something that the spline coefficients fitted in the older AMBER method (Equation 3) can work around. Five bits is the best choice when working in 32-bit floating point numbers. For extreme cache conservation, indexing with four bits of the mantissa and the square of the target function might be viable, but indexing by five bits of the mantissa and the target function’s unmodified argument would produce slightly higher accuracy for the same overall table size.

**Table 4:** Mean accuracy for spline-based interpolation of the PME particle-particle interaction force with different spline basis functions, spline arguments, and indexing bit counts. All values are given in units of 10*^−^*^6^ kcal/mol-Å. **(a)** The number of bits of the floating-point input argument’s mantissa used in conjunction with its exponent to determine the spline table index **(b)** Splines are defined by the polynomial in equation 6 **(c)** Splines are defined by the fractional power series in equation 3

| Indexing Bits<sup>(a)</sup> | Polynomial Spline Basis<sup>(b)</sup> |  | AMBER Spline Basis<sup>(c)</sup> |  |
| --- | --- | --- | --- | --- |
| | $r$ | $r^2$ | $r$ | $r^2$ |
| 4 | 27.15 | 4.51 | 20.82 | 4.00 |
| 5 | 2.75 | 1.58 | 19.72 | 3.93 |
| 6 | 1.52 | 1.43 | 19.37 | 3.88 |
| 7 | 1.40 | 1.42 | 19.66 | 3.89 |

In AMBER, moderate additional improvement in the accuracy of the splines was obtained by optimizing the 32-bit floating point coefficients such that, when evaluated for many points in the relevant range with 32-bit floating point math as they would be in a “real world” scenario, the outcomes fell as close as possible to the 64-bit benchmark. Similar code was added to STORMM, allowing this optimization to be quantified in terms of improvement per “units of least place” that each coefficient was allowed to wander. As shown in Figure 5, the optimization is more effective for splines based on the older form (inverse powers of the input argument), but even before refinement the new method based on a traditional cubic spline of the mantissa’s detail bits is more accurate.

**Figure 5:**
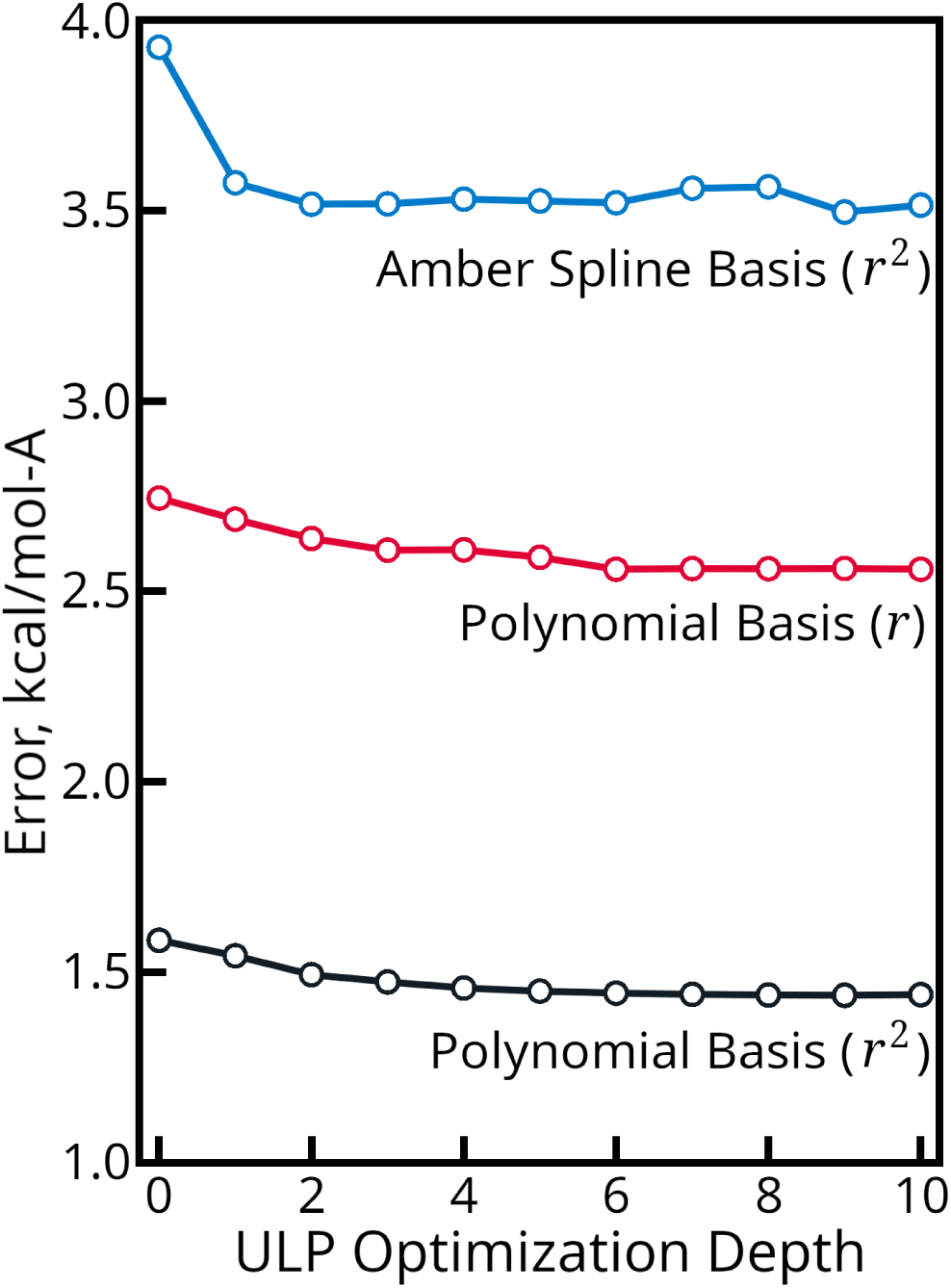
Error in spline estimates of the electrostatic PME particle-particle force after coefficient optimization. The Units of Least Place (ULPs), the lowest bits in each coefficient, are optimized by perturbing them within the stated limits and searching all combinations to find perturbations that yield more accurate results when evaluating the splines in 32-bit arithmetic. Curves for different spline basis sets are indicated on the figure, along with the indexing method. Splines based on the AMBER basis functions and indexed by the function argument are ten times less accurate and off the plot scale.

For interactions that will never sample a range near zero in any practical situation, it is appropriate to index the table starting from an infinitesimal value as the vast majority of the indices will be ignored. However, for target functions that are well enough behaved and could be sampled near zero, rolling an offset into the indexing parameter can be very helpful as it puts a lower bound on the width of each spline segment as the distance argument approaches zero. An offset of 0.5 does not preclude interpolation for values below 0.5, it ensures that there is a single group of, say, 32 spline segments with width 0.015625 spanning the range [0, 0.5). This can conserve cache as well as avoid problems when fitting a functional form over infinitesimal separations becomes unwieldy for the machine precision. In general, adding an offset to the indexing parameter takes a toll on accuracy at short range, e.g. 0.5 Å to 4 Å depending on the severity of the offset, but the effects on a function like the PME direct space sum are minimal for small offsets, and splines tailored to smoother functions exhibit little appreciable increase in error until the offset becomes much larger.

As mentioned above, STORMM implements the LogScaleSpline object as a template. Splines implemented in 32-bit floats approximate a target function calculated in 64-bit math, but splines implemented as 64-bit floats can approximate a benchmark computed in yet higher precision (“long double” in C/C++ syntax). While the results for 64-bit spline coefficients are outside the scope of this presentation, all of the same effects can be seen with spline tables indexed by 8, 10, or even 12 bits into the mantissa. Interpolated results can likewise exceed the accuracy of direct 64-bit computations. The main drawback to double-precision spline tables is the memory requirements of the table, which can be tens or hundreds of megabytes, and the caching requirements of the most useful segments, which scale as 32 bytes per spline (up from 16 for 32-bit floats) times 2**^M^**, where **M** is the number of high bits in the mantissa used for indexing. However, this technique could still be a useful replacement to some existing tabulated functions, on CPU resources with copious L2 and L3 cache.

The utility of the LogScaleSpline object can be extended not just to other target functions but also to other spline forms. Any expression **A***f* (*x*) + **B***g*(*x*) + **C***h*(*x*) + **D** could be acceptable, and the precise range of *x* obtained from the low bits of the original argument’s mantissa opens many possibilities. It is often advantageous to use simple basis functions that have forms similar to the target function, and depending on the context some forms may be more convenient based on quantities that must be computed for other purposes.

### 5.3 Advanced Work Units for Valence Interactions

In a CPU-based simulation, particularly running at low parallelism, the valence interactions are a trivial calculation, about 2-4% of the overall wall time. Here, “valence interactions” denote all bond, bond angle, dihedral, CHARMM Correction Map (CMAP)[25], and attenuated non-bonded interactions (usually between atoms separated by three molecular bonds). The number of interactions scales in proportion to the number of atoms connected by such features. Water, the most common component of many simulations, has no dihedral angles (the most numerous valence interaction, and one of the most laborious to compute) and often no flexible bonds. However, each of the terms can involve a large number of memory transactions to obtain coordinates and parameters, a fact which, in naive implementations, favors CPUs over GPUs. The ease with which CPUs can accomplish valence interactions has led some GPU programs[19] to support computing paradigms whereby all coordinates are transferred to the host RAM for these and other memory-intensive calculations, e.g. constraint algorithms, to proceed. However, even naive GPU implementations can be competitive given the latency and bandwidth requirements of downloading coordinates, then uploading forces, through the machine’s PCIe bus.

Optimizations in the AMBER code[24] included a set of work units, created by several thousand lines of C code running on the host after the topology was first loaded, to improve the performance of this aspect of MD simulations. The strategy involved stepping across the lists of valence interactions and adding atoms as needed to a short list (maximum 128 atoms). As more atoms were added to fulfill one interaction, the list was checked for other interactions that might also be fulfilled. As the buffer of atoms filled up, additional interactions seeded new lists. A “work unit” then consisted of a collection of atoms to import into local chip cache on some GPU streaming multiprocessor followed by a series of instructions for the valence interaction type, its parameter indices, and the local indices of cached atoms it applied to (see Figure 6). Evaluating all of the covered interactions would accumulate forces and energies, also in the local cache, for a buffered write back to the __global__ force tallies at the end of the work unit assignment. The strategy was found to expedite dynamics in AMBER18 and beyond, although the benefits were greatest in large systems and small protein-in-water simulations such as the dihydrofolate reductase benchmark (23,558 atoms) did not gain (or lose) much speed. Even with the caching optimizations, the kernel was found to be memory-bound, though not in as severe a way as the earlier, naive method in AMBER16 (direct queries of all atoms and parameters for each interaction).

**Figure 6:**
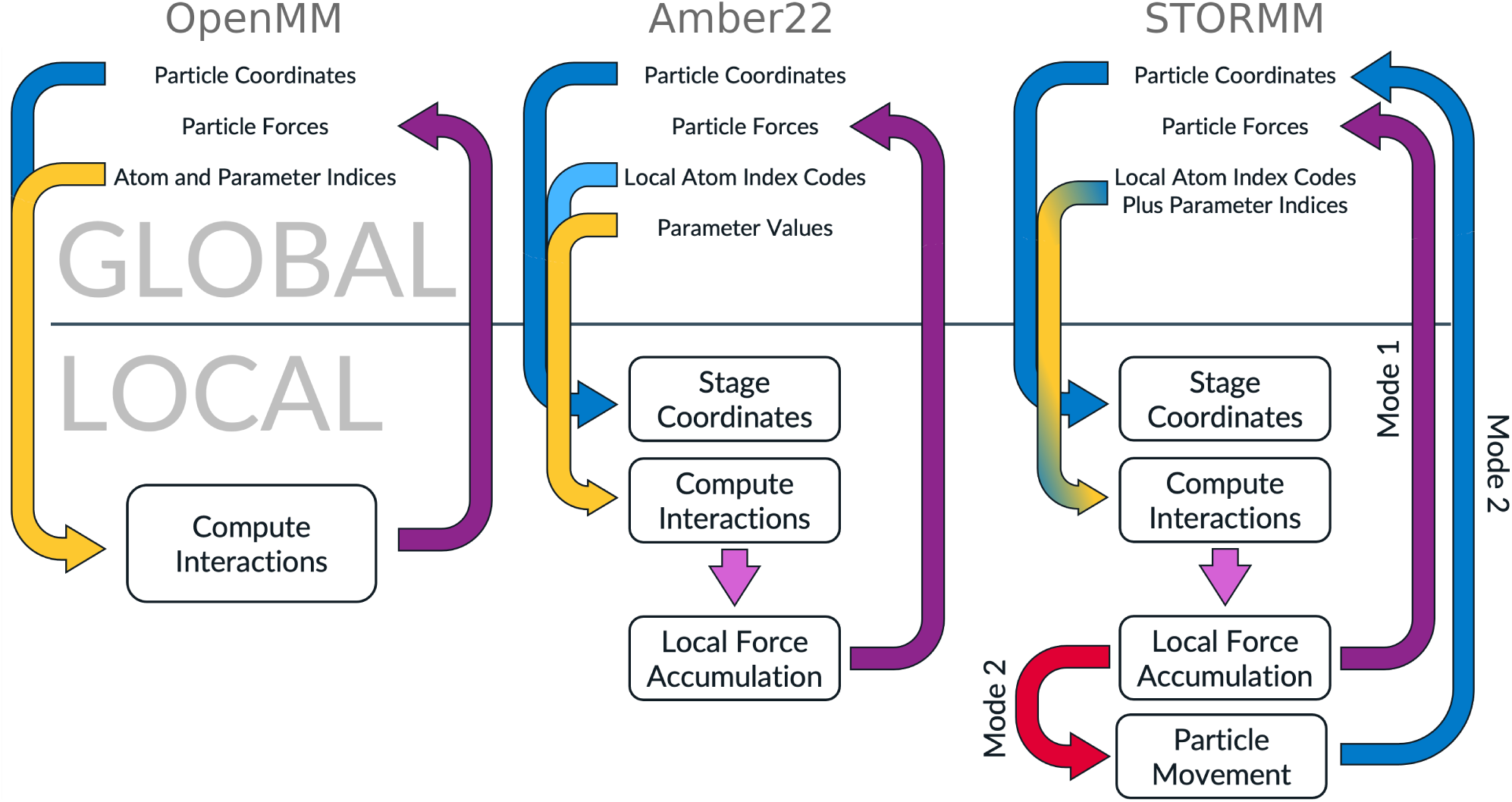
Workflow for valence calculations in OpenMM, AMBER22, and STORMM. Transfers of coordinate data are indicated by blue arrows, parameter data by yellow arrows. Purple arrows connote atomic accumulation of forces. Locations of various actions and data arrays are divided between local (on chip) and “global” memory. Staging coordinates and local force accumulation are, broadly speaking, examples of explicit data caching. Implicit caching effects are important to OpenMM with coordinates, and to both OpenMM and STORMM with parameter tables, although neither is shown on the chart. STORMM’s valence work units can function in either of two modes to serve different molecular mechanics calculations.

Table 5 shows that the AMBER code still spends a considerable amount of time processing valence interactions, and on systems of 100,000 atoms or more the modern GPUs continue to perform in proportion to the bandwidth of their __global__ memory buses. Timings were obtained by forcing each code to repeat the valence calculation many times, inserting the extra work after forces had been applied to move atoms and just before the buffers were cleared to ensure that the dynamics remained stable. The cost of the particular kernel was isolated from the differences in wall clock time needed for the original and modified codes to run a given number of steps, a technique that will be used again in Section 5.4.

**Table 5:**
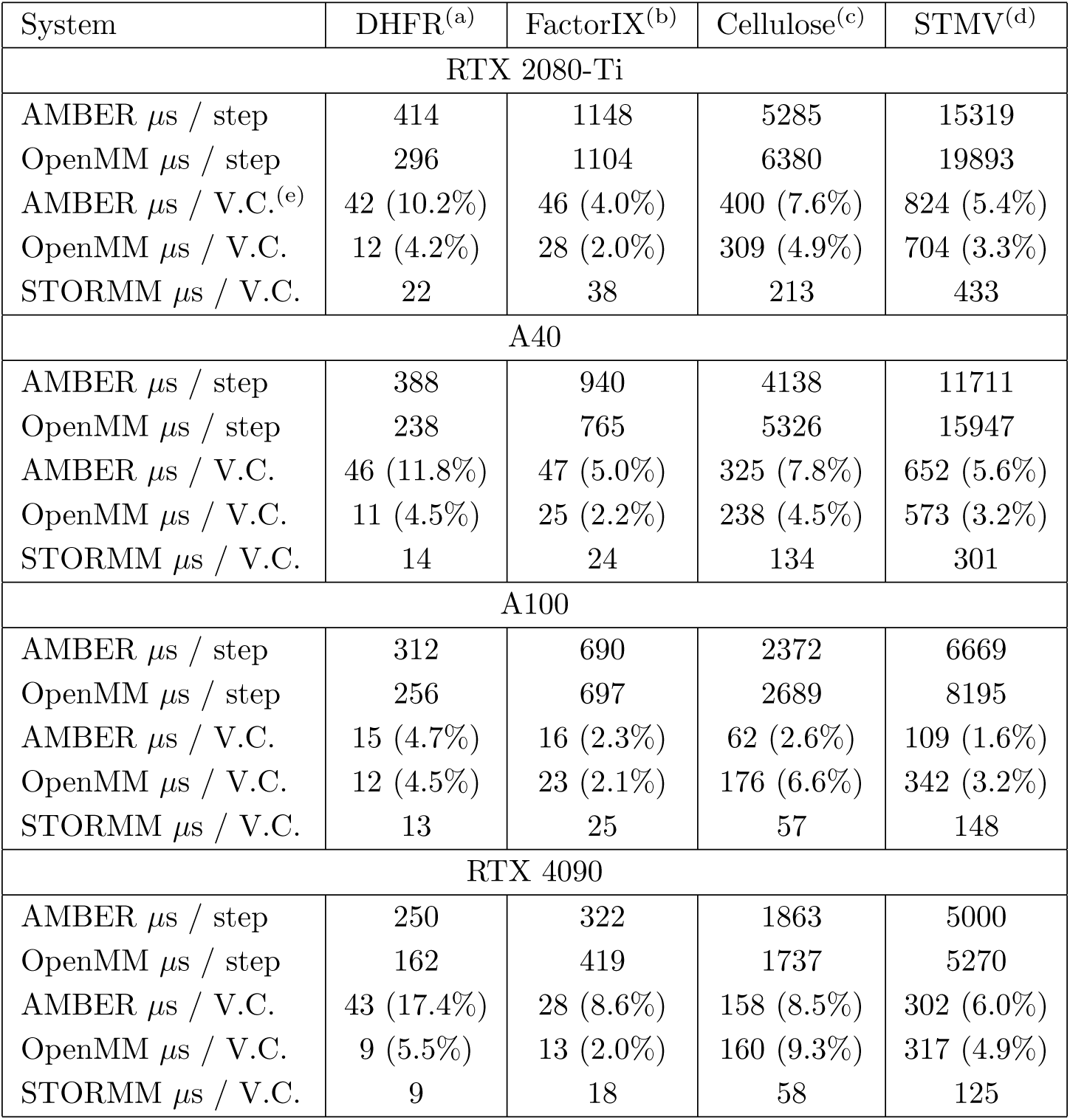
Timings for AMBER and OpenMM 8.1 running four AMBER benchmark MD systems. Run conditions: 4fs time step, constant volume, temperature maintained at 300K by Langevin thermostat, 9.0 Å cutoff on short-ranged electrostatic and Lennard-Jones interactions, TIP3P water. PME was applied only to electrostatics, with each program’s default behavior determining the grid density. Overall timings for each time step were compiled with a post-release modification to 8.1 (see Supporting Information, where timings in ns/day are availble), but the valence calculation timings will be consistent with the 8.1 release version. **(a)** Dihydrofolate reductase, 23558 atoms (including 21069 in water) **(b)** Factor IX, 90906 atoms (including 85074 in water) **(c)** Cellulose fibers (in water), 408609 atoms (including 317565 in water) **(d)** Satellite Tobacco Mosaic Virus, 1067095 atoms (including 900159 in water) **(e)** Each code’s cost of valence interaction computation is likewise given in microseconds. The percentage of time spent in this section of the code is given in parentheses for complete MD packages.

STORMM uses an approach inspired by the AMBER design, with critical refinements. The work units are laid out based on graphs of contiguous atoms, not passes over specific valence terms. Figure 6 compares the approaches. In AMBER, each work unit is expected to contribute its accumulated forces back to__ global__ tallies, but in STORMM they may either follow a similar track or become the capstone of the force calculation, reading the results of prior non-bonded computations and then updating the positions of the atoms they have already cached. Constraints, virtual site placement, and even the thermostat may thus fuse with the valence interaction kernel, saving launch latency and eliminating the need for particles to make multiple round trips between the card’s RAM and the streaming multiprocessors. In order to cover all valence interactions, there is overlap between work units: each work unit comes with a mask of atoms which it is responsible for moving. In the most complex cases, such as virtual sites with frame atoms subject to their own constraints, each work unit understands an order of operations and comprises all atoms whose positions and total force contributions must be known in order to complete its position update assignments. Some valence interactions are computed by more than one work unit in STORMM, in the interest of having the results for force transmission from virtual particles and the final coordinate update. If required, energies for such redundant calculations are likewise tallied only by one work unit responsible for doing so, according to the plan laid out on the CPU after loading the topology.

As is evident in Table 5, the STORMM valence work units are superior to the AMBER implementation, as is OpenMM’s implementation except when AMBER can work with a strong memory bus. AMBER’s implementation lessened but did not eliminate the memory bottleneck associated with valence evaluations. While it caches atoms and performs local accumulations of their forces, the process of computing each dihedral term still involves reading a 64-bit instruction containing the atom indices followed by copies of the force constants (amplitude, periodicity, and phase angle). The second-most numerous type of valence interactions, attenuated “1:4” non-bonded interactions, likewise involve import of new values for each pair. In contrast, STORMM’s 64-bit instructions (the dihedral is illustrated as an example in Figure 7) compress the data much further and use additional indices into parameter tables which the kernels take care to conserve at various levels of cache. STORMM exploits the strong association between dihedral terms and 1:4 attenuated non-bonded interactions to reduce the number of instructions and atomicAdd() operations they entail. Furthermore, STORMM exploits the fact that many dihedral terms apply to the same quartet of atoms, differing only in amplitude, periodicity, and perhaps the phase angle. In such cases, or to accommodate bespoke force fields with more than 65535 unique dihedral parameters, STORMM’s dihedral instructions can refer to a secondary 32-bit instruction encoding another cosine series term. With 8 bytes read from__ global__ (and not cached, as it is used once per time step), STORMM obtains information that may take 48 bytes to convey in the AMBER work units, or 32 bytes in OpenMM’s scheme. In its extended 12 byte dihedral instructions, STORMM transmits information that would take 76 bytes in AMBER and 52 bytes in OpenMM.

**Figure 7:**
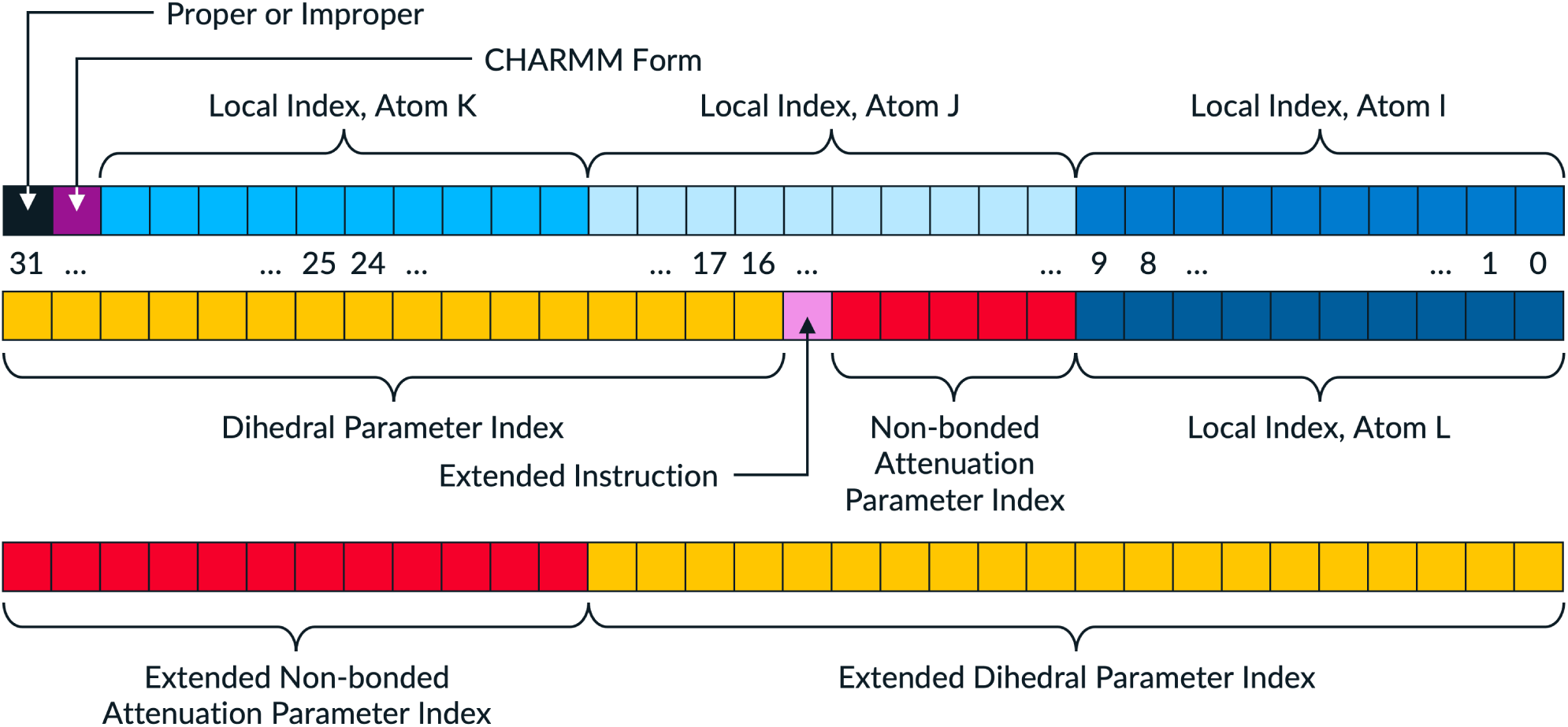
Map of the bitmask in a STORMM dihedral instruction. The basic dihedral instruction consists of 64 bits, including 40 for atom indices (I, J, K, L) in the local cache, 16 for the term’s parameter index (a code of 0xffff in these bits indicates zero amplitude), and 5 for the I:L non-bonded attenuation strength (a code of 0x0 in these bits indicates no non-bonded interaction). Other bits are used to differentiate proper and improper dihedrals (STORMM logs these energy components separately), and whether an additional cosine term should be applied to the same four atoms (dihedrals with three or more terms in the cosine series would entail new instructions targeting the same atoms). The extended instruction accommodates up to a million unique dihedral parameters and 4096 unique non-bonded attenuations.

OpenMM compiles a unique kernel for evaluating bonded interactions in the simulation at hand, although only the overall numbers of each term are hard-wired. (The benefit is that only those terms present in the actual model are then compiled, minimizing register pressure by omitting terms with complex expressions when they are not used.) This is by far the simplest approach, as shown in Figure 6. Atom indices and term parameters are still imported from data arrays in__ global__, making it much like AMBER’s original GPU implementation, although it lessens memory traffic by referring to parameter sets by their unique indices and leaning on global caching effects for atomic coordinates. It may be impossible to exceed the speed of this approach for very small systems with any model based on work units, which require various synchronizations across the thread block in order to have local force accumulation, even if the overall throughput benefits at higher atom counts. AMBER’s work unit scheme relies on a feed of raw parameters for each term, e.g. two float64_t for the a bond’s equilibrium and stiffness constants, rather than exploiting indices and parameter tables. This was one of the major changes in STORMM that compressed the data stream. In that sense, OpenMM’s implementation has features of both AMBER and STORMM, and the timings reflect it.

While all codes do the bulk of their work in float32_t arithmetic and accumulate in some sort of fixed-point scheme, the precision models differ. This carries consequences for the required memory bandwidth or floating-point arithmetic throughput and the overall speed of the kernel. Table 6 compares the methods in each code. Both the AMBER and OpenMM codes are able to produce microcanonical (“NVE”) dynamics with very low energy drift over long simulations, but for its combination of precise displacements, high-precision accumulation throughout, and safeguards against instabilities in acosf() for dihedral angles near 0 or *π*, STORMM is likely to be the most stable of the three codes in this respect.

**Table 6:**
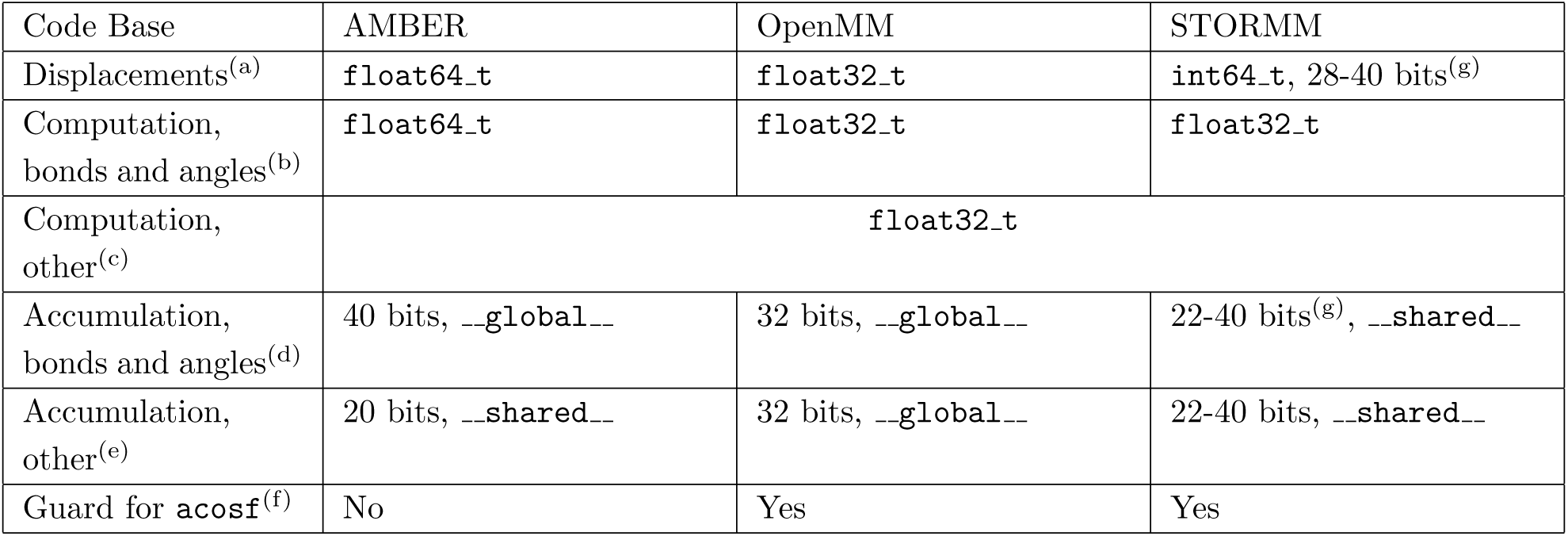
Features of the valence interaction computation in three code bases. Each code includes a run mode focused on float32_t “single” precision. These features apply to that mode; all code bases also include a “double” precision mode which raises accuracy and cost of most facets of the calculation. **(a)** The precision in which the displacement between particles is computed. Once the small displacement is computed from two (possibly much larger) numbers in each precision model, all codes convert the result to float32_t or float64_t, as appropriate to the specific interaction. **(b)** Floating-point arithmetic used in computations of bond or bond angle terms **(c)** Floating-point arithmetic used in computations of dihedral, CMAP, attenuated non-bonded interactions, and various restraint forces **(d)** The number of bits in the number’s fraction is given, followed by the location of the accumulator. Following its local force sums, STORMM is running in a mode whereby the valence interaction contributions are added back to __global__ accumulators stored in int64_t. **(e)** The method of accumulating forces emerging from all other interactions. **(f)** The arccosine function becomes unstable in float32_t computation for arguments leading to angles near 0 or *π*. The remedy in OpenMM and STORMM is to detect such situations and evaluate (or approximate) the angle with arcsine. **(g)** STORMM allows the user to select the fixed-precision model for coordinates and force accumulation, within what are deemed to be safe limits.

The advance in overall speed across a variety of systems is a combination of bandwidth optimization as well as flexible sizing in the work units themselves. While all of STORMM’s valence work units must have one consistent maximum size for a given kernel launch, the size is set based on the workload and can vary between 68 and 544 atoms. Likewise, the thread blocks that process each work unit can vary in size between 64 and 512 threads. STORMM will query the GPU and decide how best to cut the problem into pieces that will keep as much of the card occupied as possible. The benefits also extend to STORMM’s ability to run multiple systems in a single runtime instance. By design, each valence work unit tracks only one set of energy terms and takes a single offset for atom indexing into the topology and coordinate syntheses. Therefore, each work unit applies to at most one system, even if there remains space for many more atoms, but in the case of tens of thousands of small molecule problems, the smallest work unit division can still achieve good thread utilization. When multiple work units combine to evaluate a single problem, the overlap between them is modest, and with very large problems the largest work units can have overlaps as low as 1-2% of their overall content (data not shown).

Despite the results in Table 5, STORMM is working against a major disadvantage: its valence work units cover all atoms in each simulation, even rigid water molecules, which have no meaningful bond or bond angle terms and are therefore skipped by AMBER’s work units and OpenMM’s bonded interactions kernel. STORMM is anticipating that these atoms will be subject to constraints if the molecules themselves are not flexible, or that the atoms may need to be moved (both processes fall within the purview of the kernels that evaluate valence interactions). To evaluate each code in strict terms of valence interaction throughput, a basic unit of di-, tri-, and tetra-peptides, 2000 atoms in all, was assembled and compacted into a cubic box about 27 Å on a side at one atmosphere pressure. This protein-like fluid was tiled to varying degrees to create systems with up to 500,000 atoms bearing the sort of valence interactions found in typical biomolecules. No atom movement or constraint applications were requested in the STORMM kernel’s calculations, to be consistent with the established codes’ kernels: STORMM was set to operate in “Mode 1” illustrated in Figure 6. The same experiment behind Table5 was repeated for the range of systems to time each implementation. Results are presented in Figure 8: when measured as a function of the total number of protein atoms, STORMM shows the strongest performance overall, rivaled only by AMBER’s on the A100. While OpenMM’s wall time rebounds on some lower-bandwidth cards as the system size gets very large, given the logarithmic scale of the plots it is unlikely that AMBER’s implementation will ever surpass it on such hardware (the largest of the tiled systems contains three times more “protein” atoms than the mammoth STMV simulation in Table 5). STORMM’s approach also comes close to equalizing the older graphics-oriented GPUs and the A100, despite the disparity in memory bandwidth, and can muster the tremendous compute power of the RTX 4090 to produce even higher throughput (the time to compute valence interactions in a large system is *∼*0.5 *µ*s per 2000 atoms on an RTX 4090, versus *∼*0.7 *µ*s on an A100, and *∼*1.2 *µ*s on an A40). The split fixed-precision accumulation detailed in Section 5.1 shows consistent benefits, even on the A100 (on this card, the split accumulation was not as fast as the unified int64 t approach in the benchmark presented in Figure 3, perhaps because in the prior test no threads were competing for the same memory).

**Figure 8:**
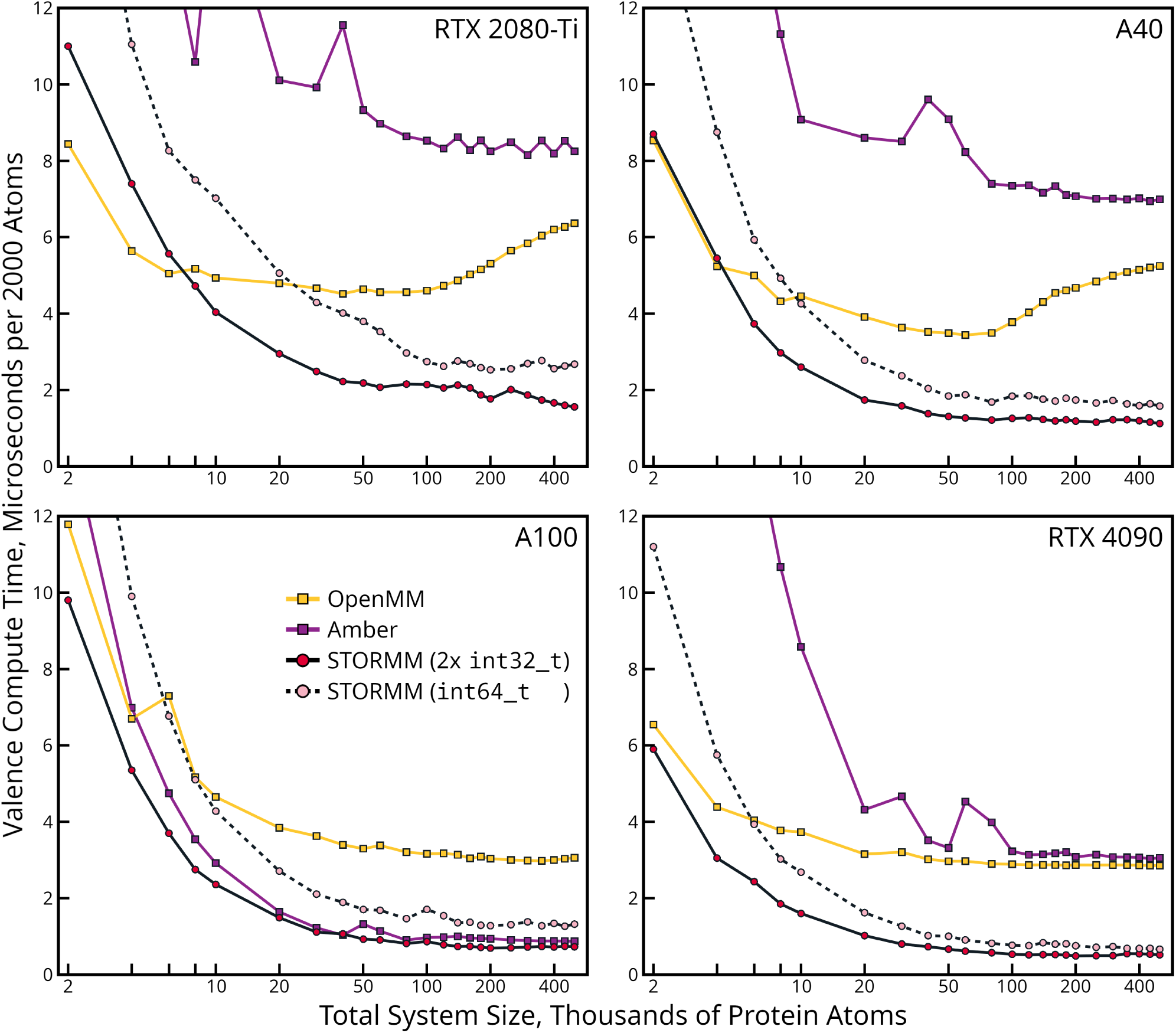
Comparison of three implementations for valence interactions as a function of total atom count. A molecular system consisting of 2000 atoms in various peptides was tiled as indicated on the *x*-axis to create specific numbers of “protein” atoms bearing valence interactions typical of biomolecular simulations. Each code’s valence interaction computation time is plotted per tile of the basic unit to show its scaling with overall system size. The inset legend in the lower left panel applies to all panels. The *y*-axis works in time per 2000 atoms as this is about the smallest meaningful increment for measuring the cards’ performance: the minimum kernel execution time in any of the codes is around 6-10*µ*s, and any of the codes would process 1000 atoms in the same time it takes to process 2000.

While it is clear that STORMM’s implementation depends less on memory bandwidth, the result for A100 in Figure 8 calls into question other “macroscopic” features of each approach. AMBER is launching three 128-threaded blocks per streaming multiprocessor (SM), a total of 384 threads. Up to Volta, it was found that AMBER’s memory-bound kernel did not improve in performance when launching more threads, although the late-generation A100 may provide yet higher returns if it launched 512 or 768 threads per SM. In contrast, STORMM benefits from launching 1024 threads per SM, the maximum possible number given the 2^16^ available registers, yet it is barely faster than AMBER if memory bandwidth is plentiful. The simplest explanation is that STORMM is doing more with more. STORMM’s check for numerical stability in acosf() and processing of redundant terms for the sake of having complete forces on particular atoms no doubt take a toll. More important is that while AMBER is accumulating the bulk of its calculations in int32 t with coarse 20-bit fixed precision, using “fire-and-forget” atomics with no requirement to check the return value, STORMM is using somewhat higher precision and checking for catastrophic overflow with each atomic addition. As will be seen in the following section, waiting on the return value of atomicAdd() is costly, albeit necessary to provide freedom in the precision model. The fact that split accumulation makes such a dramatic improvement over direct accumulation in int64 t (comparable to the effect in Figure 3) shows how much of the process lies in keeping the tallies.

Valence work should not be a major effort in biomolecular simulations, and the problem is compacted to an appropriate degree in most OpenMM simulations. AMBER’s work units, while a step in the right direction, remain dogged by excessive __global__ traffic due to the layout of some basic parameter arrays. In applications such as membrane simulations, as much as 25% of the total atoms may be found in lipid or protein molecules. It pays to optimize this facet of the calculation lest the code become reliant on the strength of the card’s memory bus. In STORMM, special emphasis has been placed on this design to scale small molecule calculations, where as will be shown in Section 6.1 the valence interactions can account for more than half the GPU wall time. The versatility of the result will benefit much larger condensed-phase simulations as well.

### 5.4 Particle-Mesh Interactions in Periodic Simulations

The complete periodic simulation capability in STORMM is unfinished, but a strong functionality for one of the most memory-intensive operations is prepared and, like the valence interactions, showcases a powerful strategy in having the CPU examine the workload at hand in order to subdivide it into balanced work units that can achieve high thread utilization on the GPU. The splitting between particle-particle and particle-mesh interactions was alluded to in Section 5.2. The particle-mesh interactions in a molecular simulation follow the same steps as particle simulations in other branches of computational physics[26]:

- Interpolate particle density from the particles’ actual positions to regular points on a three-dimensional mesh.
- Perform a “forward” Fast Fourier Transform (FFT) to render the particle density in frequency space.
- Carry out another forward transform of the influence function of one mesh point on all others. This step can be skipped if the influence function has an analytic form in frequency space, as it does in the case of PME.
- In frequency space, multiply all elements of the particle density with the mesh influence potential.
- Complete the convolution with a reverse FFT of the above product, obtaining the mesh potential based on the density at every point projecting its influence onto all others.
- Map the mesh-based potential back to the actual particles based on the interpolation used in the first step. Derivatives of the interpolant indicate the forces on each particle along the mesh axes, which can be transformed into Cartesian space using chain rules.

Of these steps, the FFT is oft reviled as the most difficult to scale. In principle, this is where the global communication occurs, but the FFT accomplishes a tremendous amount of work at an astonishing economy in arithmetic. The influence of one point on the lattice can be extended to all others by computations involving all points directly to the left or right, then all points directly forward or behind, and last all points directly above or below: all-to-all interactions processed between every pair of *N* ^3^ points for communicating just 3*N* pieces of information (twice). While it is desirable to minimize global communication requirements, the only way in which this can happen is to reduce the density of the mesh on which global interactions occur, and many alternatives to PME[27, 28, 29] still perform FFTs on a mesh spanning the entire periodic simulation cell. On GPUs, the greatest obstacle to reducing the global mesh density in canonical PME is the communication required to map particle density to the mesh . Each particle transmits *M* ^3^ density contributions for an interpolation order of *M* (the reverse process, force retrieval, in principle involves the same communication, but it consists of reads rather than atomic writes). PME and other approaches use stencils based on *B* -splines to project the particle density. Details may be found in literature.[18]

As we show in Figure 9 (compare with Figure 11, presented later in this discussion), this mapping cost can be greater than the combined cost of the forward and reverse FFTs. In that Figure, a kinase simulation of 52,889 atoms (orthorhombic unit cell dimensions 85.2 Å by 90.6 Å by 81.9 Å) is tiled (taking the most compact arrangement with the given number of replicas) to explore the effect of system size on particle-mesh mapping in two popular, GPU-enabled MD codes, AMBER22 and OpenMM 8.1. In each case, the particle-mesh mapping cost was measured by adding a loop inside the main dynamics loop to reset the density mesh accumulators to zero, launch the accumulation kernel, then re-arrange and (if needed) convert the mesh to float32_t suitable for FFTs. Repeating this loop ten times and then comparing the total run time of the adjusted code with that of the original code over 5000-100000 time steps yielded an estimate of the average particle-mesh mapping cost. The OpenMM code also includes a step to sort all particles based on their mesh alignments, which is reused in the force interpolation at the end of the particle-mesh workflow. This sorting cost was measured in the same manner as before, and half was assigned to OpenMM’s overall mapping procedure. These timings can be compared to STORMM’s two options for mapping particle density to the PME mesh. Both are based on a neighbor list that locates particles to a spatial decomposition grid subdivided by half the particle-particle cutoff distance. Another unique aspect of STORMM is that the PME mesh is aligned to the neighbor list decomposition grid: an integer number of mesh spacings (e.g. 4 or 5) must span each neighbor list decomposition cell in all directions. The first option, “global accumulation,” loops over all atoms issuing atomicAdd() instructions to the global (L2) cache. In the second (and more advanced) option, a kernel loops over “local work units” spanning the spatial decomposition cells and works within __shared__ memory resources, a partition of the local (L1) cache on each GPU multi-processor. As with the valence interaction work units, the particle-mesh mapping instructions are drawn by C++ routines to optimize cache utilization and load balance. The “local work unit” kernel has nearly the same breadth of applicability and choice of interpolation order as the “global accumulation” kernel, unless an extreme PME mesh discretization like 8 or 10 per neighbor list cell is chosen.

**Figure 9:**
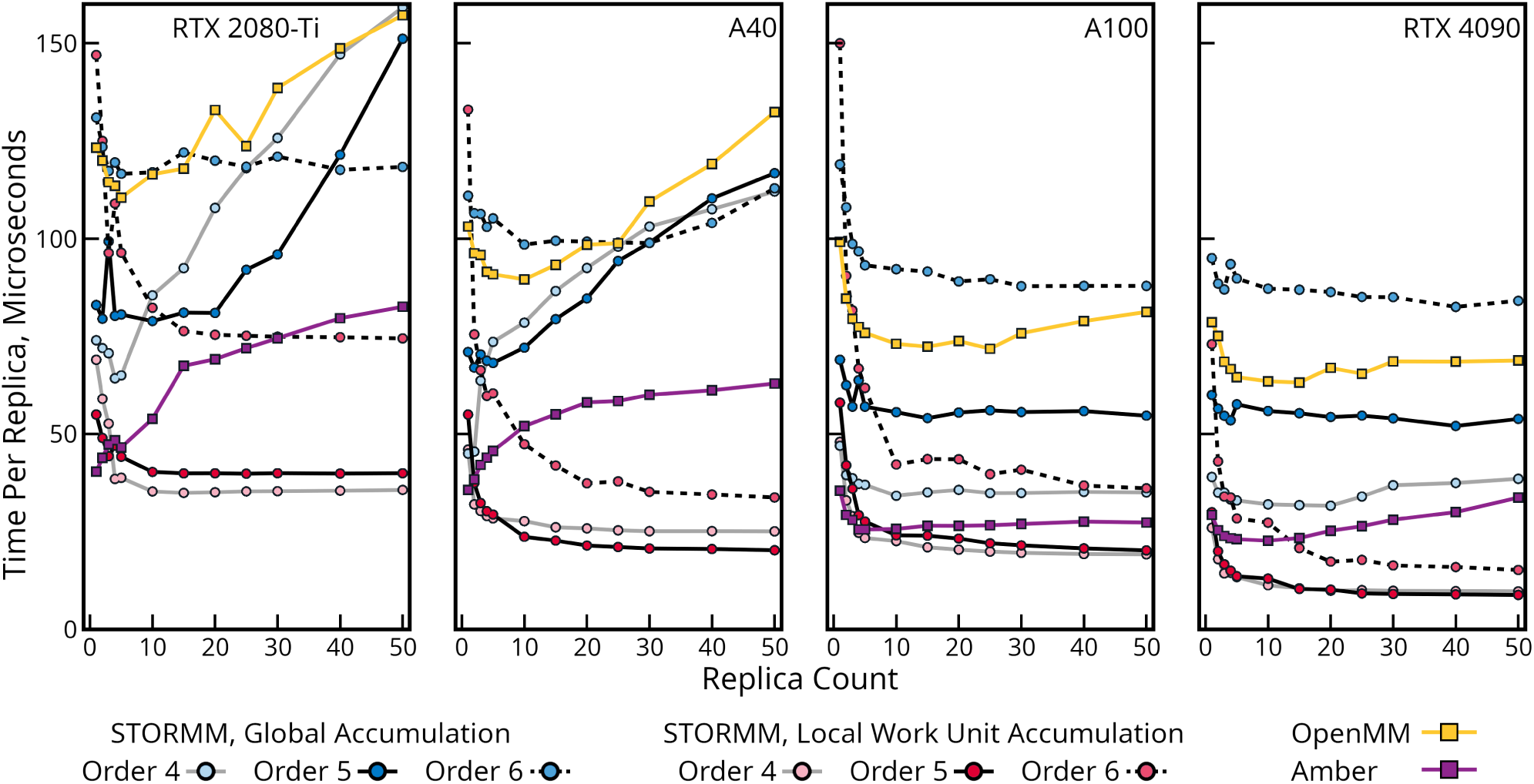
Time needed to map charge density to PME meshs. Two STORMM kernels (one working in L2 to perform “global accumulation”, the other working out of a __shared__ partition of “local” L1, based on CPU-designed work units) as well as the equivalent code in AMBER and OpenMM 8.1. STORMM kernels can work in a range of interpolation orders, as shown in the legend, and in this test are performing fixed-precision accumulation with 28 bits after the point. AMBER performs fixed-precision accumulation on a similar scale but is restricted to 4^th^ order interpolation. OpenMM runs 5^th^ order interpolation, and for this test is configured in non-deterministic mode, accumulating in float32_t to approximate a similar bandwidth requirement to STORMM and AMBER. The system is a 52,889 atom kinase in water. Mesh discretizations in STORMM were set such that voxels of the simulation close to 4.5 Å in width were filled with 125 points in 4^th^ order interpolation, 64 points in 5^th^ order interpolation, or 27 points in 6^th^ order interpolation, which approximate the mesh densities of equivalent calculations in AMBER and OpenMM. STORMM treats each replica of the system as a separate problem while AMBER and OpenMM are tasked to simulate a tiled unit cell containing the stated number of copies of the system. GPU properties are listed in Table 3.

Robust benchmarking that covers the range of relevant use cases and scenarios is hard. One important thing to note in Figure 9 is that, while the established MD codes are running calculations on a tiled system for the purposes of comparison, in this test STORMM’s kernels are required to treat each replica of the system as distinct, as would be the case in a replica exchange or weighted ensemble simulation (in Figure 8, all codes were operating on a single unit cell). The traditional codes are therefore able to see all atoms and the PME meshes as contiguous in memory, whereas STORMM is respecting barriers between systems and any undersized work units or disjoint memory spaces this may entail. A precise comparison is also elusive due to the high-level configuration of each engine. In the case of AMBER’s code, which runs the charge density mapping in the same stream as the rest of the force calculation when performing standard MD, the comparison to our test program driving the STORMM kernels is strong. However, OpenMM defaults to running its memory-intensive density mapping and FFT work in a separate stream from the compute-intensive particle-particle non-bonded work. Toggling this feature showed that the complete MD cycle benefits by as much as 15-20% on cards with plentiful L2, but on all cards the effect tends to diminish with system size and with limited L2 it can even become a detriment (data not shown). In any case, OpenMM’s streaming applies to the entire PME workflow, from mapping to force interpolation, whereas the present study seeks to examine the efficiency of the mapping in the limit of many atoms. The true cost of the density mapping in OpenMM might not be as high as indicated by Figure 9, but the OpenMM tests were repeated with and without a separate PME stream, making little change in the mapping times or trends (data not shown), and experience from programming the STORMM kernels as well as AMBER’s approach also suggests that the benefit of interleaving this particular part of PME would be minor.

Despite a minor handicap, STORMM’s mapping kernels perform very well. The “global accumulation” kernel’s timings include the required initialization and accumulator conversion steps as performed in AMBER and OpenMM. It is intermediate in speed between the two established codes although its throughput deteriorates the most rapidly as a function of the total number of atoms in play. On the A100, most approaches show a degree of improvement in their throughput as the number of atoms increases–larger workloads are easier to balance among the tens of thousands of threads that each card supports. However, for the RTX 2080-Ti and A40, all L2-based approaches show a degree of regression, e.g. they run ten replicas of the kinase system more than twice as fast as they run twenty. This reflects a critical hardware limitation on some cards: the size of the L2 cache. The RTX 2080-Ti and A40 have L2 cache sizes of 5.5MB and 6MB, respectively (see Table 3). The typical density in a protein-in-water molecular simulation is about 1 particle per 9.5 cubic Å ; for 4^th^ order interpolation this is a density of about one particle per ten mesh points, for 5^th^ order a density of about one particle per five mesh points, and for 6^th^ order a density of about one particle per three mesh points. With each accumulator being 4 bytes, the single replica of the kinase then has a mesh size of about 0.8 - 2.4MB. Adding a second replica may approach the limits of the cache; third and fourth replicas can cause it to start thrashing as threads working on different regions of the problem issue instructions to disparate memory addresses. The A100 (80GB PCIe), a late-generation model with an expansive 80MB L2 cache, can handle tens of replicas of the system with even the densest meshes before L2 is filled, and even then the massive resource will have a low rate of overturn: the A100’s powerful memory bus (which has 170% more bandwidth than the A40) can keep up. It is clear from Figure 9 that the design of the algorithm, in particular the pattern in which different particles are mapped, matters. While slowest overall, OpenMM’s approach, with its careful pre-sorting of particles to optimize the way it cascades across the mesh, shows less performance degradation than either of the other kernels working out of L2 on the A40 or RTX 2080-Ti. The fastest of these L2-based kernels, in AMBER, uses some aggressive rearrangement of the accumulators to optimize coherence of all threads’ write instructions. Whereas STORMM’s kernel is writing to *M* ^2^ cache lines per atom (and perhaps twice that many, if the mapping wraps across the periodic boundary), AMBER’s kernel is mapping to as few as *M* and at most 4*M* . There are also redundant calculations of the same *B* -spline coefficients in the STORMM kernel, assigning *M* threads to each atom rather than just 1, whereas the AMBER kernel assigns multiple threads to each atom but pools the computations of the *B*-spline coefficients. However, the optimizations in the AMBER kernel impose limitations of their own, and STORMM’s “global accumulation” kernel is meant to serve edge cases that users might request.

The pervasive degradation of performance with higher atom counts, a sort of digital Tragedy of the Commons where each thread going about its work starves the others of a critical resource, challenges the premise of gaining efficiency by stacking related simulations together. The same problem is likely seen in Figure 8, as OpenMM’s L2-focused atom staging and force accumulation begins to show regression on the same cards. To overcome this, STORMM’s “local work unit” kernel illustrated in Figure 10 again pulls the problem closer to the chip and provides the best performance in nearly all cases, except perhaps the smallest number of atoms and the lowest interpolation order, where AMBER is faster on older hardware. Even on the A100, the most favorable circumstances for the traditional L2-focused methods, STORMM quickly overtakes the performance of the AMBER engine running 4^th^ order interpolation with its own 5^th^ order interpolation. On other cards it is even be able to perform 6^th^ order interpolation in less wall time than AMBER.

**Figure 10:**
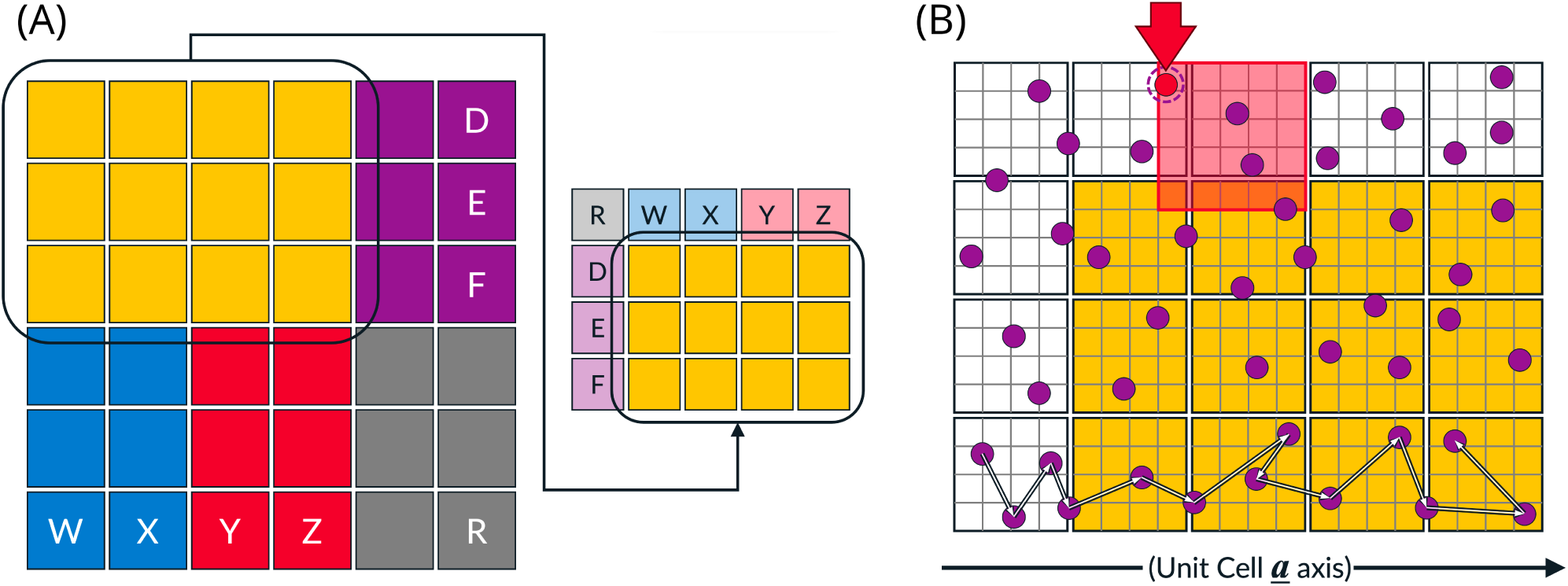
Two-dimensional illustration of work unit design for particle density mapping in periodic simulations. In panel A, a periodic unit cell is subdivided into colored work units. The PME mesh regions assigned to each work unit are non-overlapping, but particles in a “halo” region around each PME mesh subdivision must be imported by any given work unit. Wrapping in the periodic boundary conditions is encoded into work units as needed. The three-dimensional implementation in STORMM includes variable heights for each work unit. During setup, work units are arranged in decreasing order of size, so that the shortest work units can back-fill the GPU thread blocks and improve load balancing. Panel B details an individual work unit, composed of neighbor list partitions (black squares) which align the PME mesh (grey lines). Particles in neighbor list partitions are found in contiguous arrays stepping along the unit cell’s *a* axis (the Cartesian *x* axis in an orthorhombic unit cell), but within any given partition the atoms are expected to be randomized. The work units are designed such that GPU thread groups step along atoms as illustrated by the series of white arrows. The *B*-spline stencil due to one selected particle is marked in red: for an interpolation order of *M* , a work unit must import atoms in a halo region extending *M −* 1 PME mesh points from its assigned section of the PME mesh (colored gold in Panel B). STORMM will detect the card in use and tailor work units to utilize the available L1 cache space. STORMM seeks an *a* axis length of the PME mesh assignment that is not a multiple of 8, 16, or 32, to help distribute work across __shared__ memory banks.

New technology can help to a degree. The RTX 4090 has ample cache resources, and L2-based methods indeed show less regression even in very large systems, but none of the L2-based methods run much faster on the RTX 4090 than they did on the A100, or when kept within the L2 capacity of the A40. The L2 instruction bottleneck evident in Figure 3 may be bearing down. While OpenMM appears to show a modest improvement on this card, closer inspection suggests that most of the difference is in the sorting procedure, which takes 30 to 50 *µ*s per replica on A40, 30 to 60 *µ*s per replica on A100, and only 14 to 35 *µ*s per replica on RTX 4090. In contrast, STORMM’s “local work unit” implementation delivers more than double the density mapping rate on RTX 4090 as that seen on A40 or A100. The extreme throughput of this unique implementation, which was tuned on the Ampere cards prior to any tests on RTX 4090, suggests a dividend in “future-proofing.”

The “local work unit” kernel’s first strength is that it needs no initialization or conversion kernels to support it. The accumulators are initialized in __shared__ memory and converted to the float32_t result, which is then written to __global__ memory in its complete state. Separate initialization and conversion operations, which can take about 10*µ*s per replica of the kinase system, account for at least 20% of the run time of STORMM’s “global accumulation” implementation. More significant is that the “local work unit” kernel issues atomicAdd() instructions to __shared__ memory, which as shown in Figure 3 can be much faster than L2. Even with a spacious L2, the next fastest kernel in AMBER only gains about 25-35% throughput in very large systems, and with limited cache its throughput can degrade by more than half. In bypassing L2, STORMM’s “local work unit” kernel scales with the compute capacity of each card and can more than triple in throughput as more atoms are added.

While the L2 sensitivity of the particle mapping can be mitigated or avoided depending on the algorithm design, the L2 sensitivity of the next phase of the particle-mesh workflow, the FFT, is strong and unavoidable. The L2 cache is the lowest-latency coherent resource for the GPU, and while an extensive analysis of FFT performance and algorithm design is outside the scope of this paper Figures 11 and 12 support an informed discussion of the strategy within STORMM. There are two performance curves evident in Figure 11: the first, when the entire mesh fits in the chip’s L2, is dominated by the card’s clock speed. All of the GPUs tested become saturated, that is they can reach their asymptotic performance limit, with problems that would be associated with MD simulations of about 50,000 atoms. (AMBER’s 4th order interpolation used a mesh of 80 *×* 96 *×* 80, or 614400 points per replica, whereas 5^th^ order interpolation in STORMM and OpenMM takes place on meshes with about 350,000 mesh points.) If the problem begins to spill out of L2, there is a rapid transition to the second performance regime, one dominated by the global memory bus speed (compare Figure 11 to Table 3). Batching FFTs, as NVIDIA’s cuFFT library is able to do even for the three-dimensional case so long as all problems have the same dimensions, appears to improve the odds of reaching the asymptotic performance, not open up a new and better regime (data not shown). Furthermore, in addition to the sanctioned radices of 2, 3, 5, and 7, the inclusion of radix 11 (having a prime factor of 11 along any one dimension of the mesh) has low impact on FFT performance or the chance to reach the asymptotic limit. However, the combination of radices 7 and 11, that is to make any of the three mesh dimensions a multiple of 77, risks significant slowdown in the FFTs, worsened by using “in-place” methods. Another notable feature in all panels of Figure 11 is a patch of FFTs with under half a million points which are two to three times as expensive as others of similar sizes. These are meshes with 75 (prime factors 3, 5, and 5) as their largest dimension, which appears to raise the cost of FFTs by at least 100% and even more if the second or third side also has dimension 75. Otherwise, the performance of “in-place” versus “out-of-place” FFTs is within 2-3% on all cards, as shown in Figure 12, despite NVIDIA recommending the “out of place” approach in their documentation. The “out of place” FFT not only doubles the memory requirements for storing two instances of the PME mesh of any given molecular system, it also takes twice the L2 space, accelerating the transition to the slow performance regime on cards with limited L2 and favoring “in-place” methods by as much as 30% when this happens.

**Figure 11:**
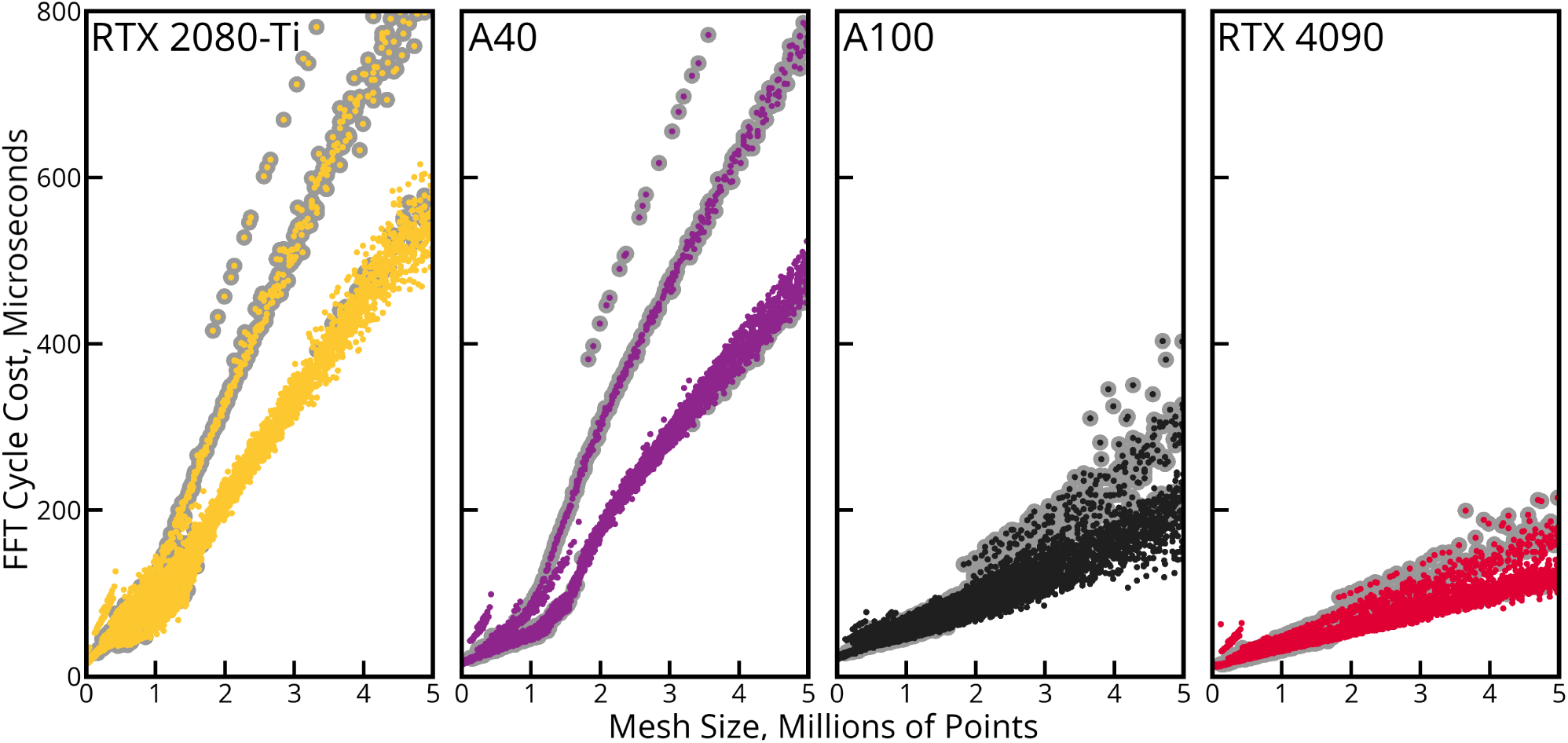
Cost of the FFT cycle in PME calculations on modern NVIDIA GPUs. FFT computation times were measured by transforming random data into frequency space and back with “in-place” methods. Sizes were selected from all combinations of mesh dimensions factoring into primes of 11 or less. Measurements are reported as the average of 200 such cycles, with the cost of re-normalizing the data removed. FFTs involving a combination of radices 7 and 11 along the same dimension are marked with grey borders. The inset legend applies to both panels.

**Figure 12:**
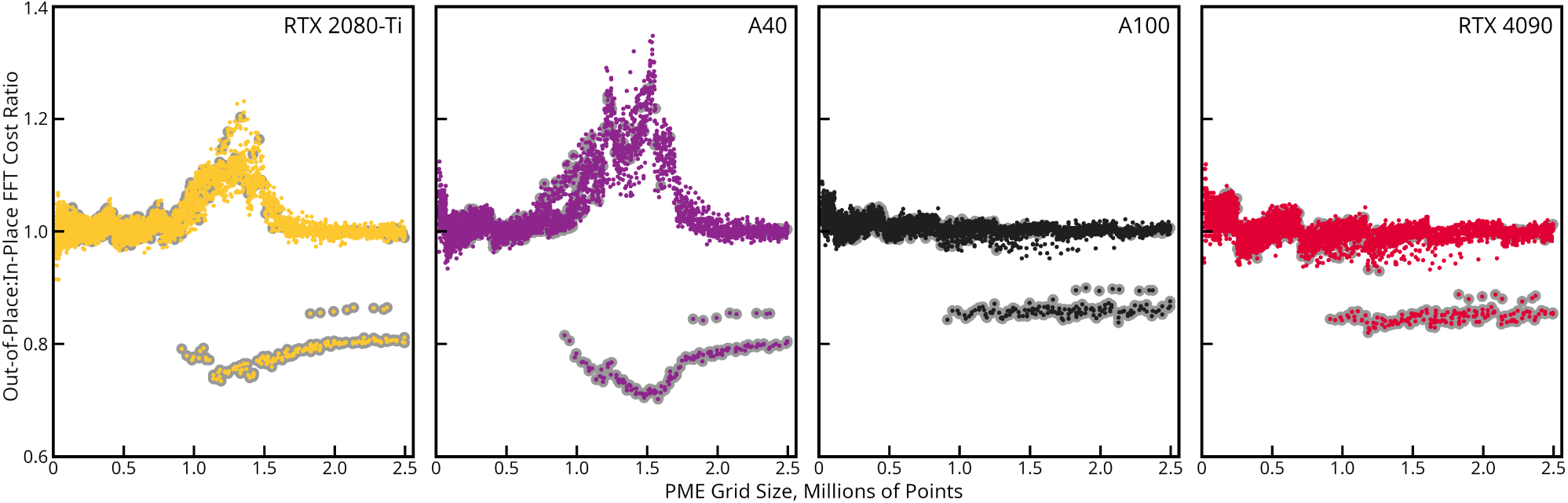
Comparison of the time cost of out-of-place to in-place FFTs on modern NVIDIA GPUs. A ratio greater than one indicates that the “out-of-place” FFT takes longer to compute than the “in-place” equivalent. As in Figure 11, PME meshes with dimensions incorporating radices 7 and 11 along the same axis are emphasized with grey borders.

For consistent performance, and to make room for more atoms and replicas on any given card, STORMM’s default FFT mode will be “in-place,” although the “out-of-place” option will also be implemented if future developments alter the landscape. When running multiple replicas or a mixture of periodic simulations, STORMM will detect the GPU’s L2 cache size and batch its FFTs in various groups or submit them in serial to keep the card working as close to the performance limit as possible without thrashing. Relief is at hand: Figure 13 shows the amount of L1 and L2 cache, per streaming multiprocessor, on various NVIDIA GPUs since compute capability 7.0. The amount of L2 cache burgeoned in various Ampere server-grade cards and continues to be sizeable in the H100 (“Hopper”) line. Copious L2 is now featured on commodity graphics cards in the Lovelace line, which helps to explain a number of performance jumps seen in very large (409k atom cellulose and 1067k atom virus capsid) systems on RTX 4090 benchmarks (see, for example, Table 5 and compare to https://ambermd.org and https://openmm.org): once FFTs and particle density mapping are unencumbered by coherent cache resources, the lower__ global__ memory bandwidth on these cards is less problematic.

**Figure 13:**
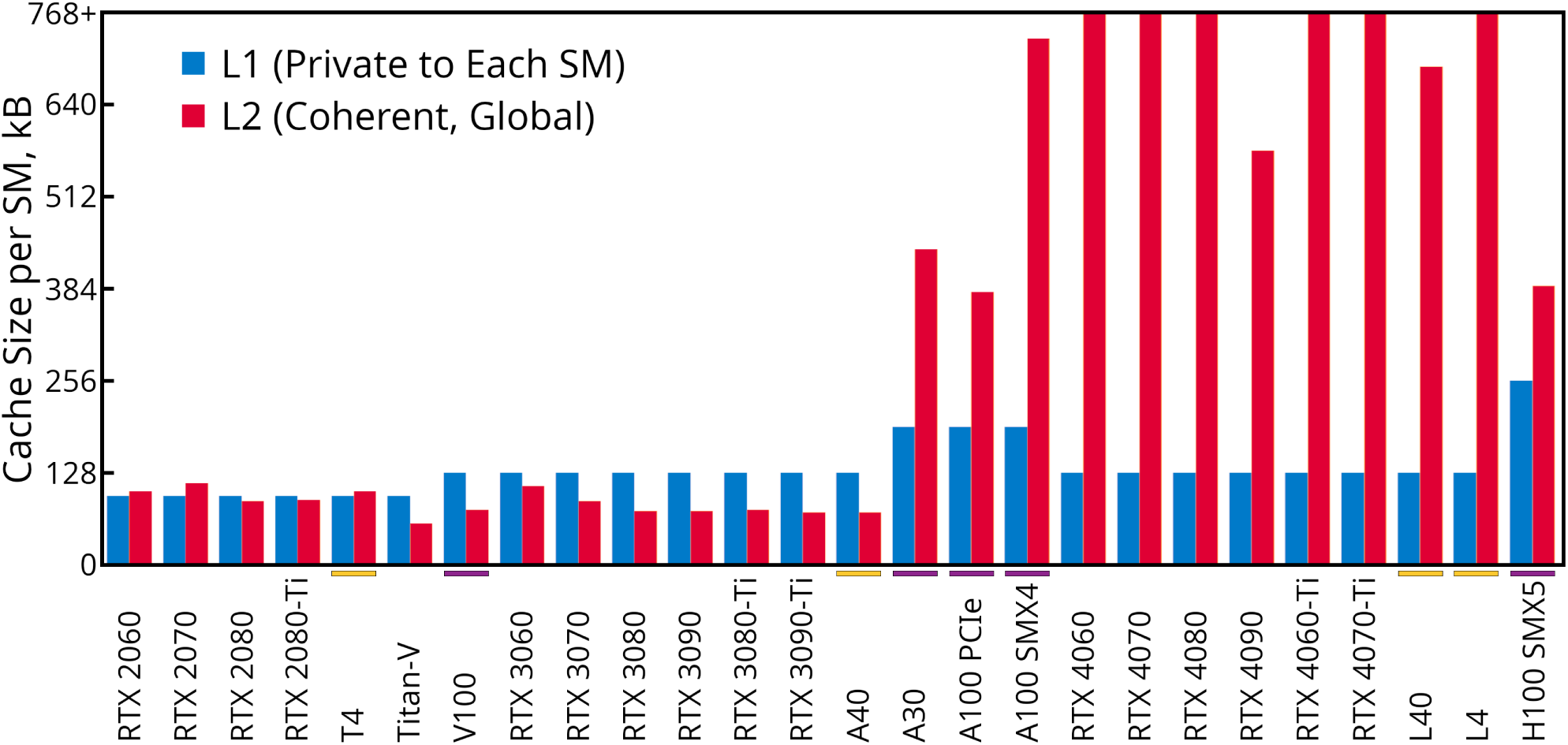
L1 and L2 Cache Sizes on NVIDIA GPUs. GPUs are listed left to right in approximate order of release date. L2 cache size per streaming multiprocessor is listed to give a better comparison with the amount of L1, which was as plentiful or more so until server-grade cards in the Ampere line, and also because the rate of overturn in L2 (thrashing) is a function of the number of threads attempting to utilize it. Cards which have high float64_t arithmetic throughput and are sanctioned for installation in rack servers are marked with a purple bar over the name. Other cards which have low float64_t throughput but are still sanctioned for installation in rack servers are marked with a gold bar over the name.

By raising the interpolation order, the mesh density can be reduced [30] while conserving the overall accuracy of the calculation. A good rule of thumb is that, to obtain forces of sufficient quality for a stable simulation, PME with 4^th^ order interpolation requires the mesh discretization to be at most 2/3 the *σ* width of the Gaussian in used in the splitting function (the Gaussian width is 1*/*(2*α*) in terms of the “Ewald coefficient” in Equation 2), PME with 5th order interpolation requires that the mesh discretization be at most 4/5 the *σ* width of the splitting function, and PME with 6^th^ order interpolation can use mesh a discretization as wide as the *σ* width itself. For a direct sum tolerance of 1.0 *×* 10*^−^*^5^, a Gaussian *σ* of 1.5 Å is obtained with a cutoff of about 8.346 Å. Longer cutoffs at the same direct sum tolerance will widen the *σ* and shorter cutoffs will shrink it in proportion. The default values of the AMBER simulation engines are built around this tradeoff: a default particle-particle interaction cutoff of 8 Å with code that tries to get the PME mesh discretization as close to 1 Å as possible, often landing at discretizations of 0.9 to 0.95 Å. It is an even trade between the interpolation orders: twice the mapping work for about half the overall FFT work, or just over three times the mapping work for a third of the FFT work. A similar analysis applies to the particle-particle cutoff: committing to three times the particle-particle work with a 50% larger cutoff would be another way to reduce the FFT work by a factor of three but without changing the amount of density mapping work. In the early days of GPU computing it was thought that this route might be taken. The fact that the old CPU parameters remain the typical (and fastest) choices for GPU molecular simulations is an indication that not enough has been done to adapt to the communication and memory characteristics of GPUs.

One possible complicating factor is that, with lower mesh densities and each particle issuing more density contributions, atomicAdd() instructions will collide on the same memory addresses in much higher rates. Global communication in the FFT is reduced at the price of intense local traffic in the mapping. However, Figure 9 suggests that, if anything, more intense calculations on a single atom are beneficial, at least up to 6^th^ order interpolation with appropriate mesh densities. In the limit of large atom counts, 5^th^ order interpolation by the “local work unit” kernel can be *faster* than 4^th^ order interpolation, and 6^th^ order interpolation has about the same efficiency as 5^th^ order in terms of the overall number of mapped points. The “local work unit” kernel is sensitive to the way the PME mesh aligns to the neighbor list decomposition: for an interpolation order *M* , all atoms needed to complete the density throughout one of the neighbor list cells can be found in that cell and those directly adjacent so long as there are at least (*M −* 1)^3^ mesh points per neighbor list cell. For the calculations in Figure 9 the combinations expected to produce forces of sufficient quality are 4^th^ order interpolation and 5^3^ mesh points per neighbor list cell, 5^th^ order interpolation and 4^3^ points per unit cell, or 6^th^ order interpolation and 3^3^ points per unit cell, with 5^th^ order hitting the “sweet spot.” In either of the other options, there are atoms in the outlying region that do not map anywhere in the work unit, and a higher proportion of atoms with stencils that have partial overlap with the work unit’s relevant mapping region. Further optimizations in the unique “local work unit” scheme which skip over some of these irrelevant atoms and stencil zones may be possible, although they would also benefit the case of 5^th^ order interpolation and 4^3^ mesh points per unit cell. Also, while a line determined by two points is not a trend, the RTX 4090 shows greater gains in the “local work units” mapping procedure than its FFTs relative to prior architectures. If future cards continue to deliver greater returns on __shared__ memory atomicAdd() relative to FFTs, 6^th^ order interpolation might become preferred.

Figure 9 compares the most similar modes of each code, in terms of the level of numerical detail and overall sizes of the meshes. The density mapping kernels in STORMM are always designed to run in bitwise reproducible, fixed-precision mode, but they are tuned to detect the number of fixed precision bits, examine the topologies at hand, and (with consideration to the particle and mesh densities) determine whether it is safe to forego the secondary int32 t accumulator. For a typical protein-in-water topology like the kinase system examined in this section, STORMM can accumulate with up to 29 fixed-precision bits after the decimal without the secondary accumulator. This is similar to the level of detail captured in the AMBER code, although we took OpenMM’s non-deterministic mode, accumulating in float32_t, as the closest match in terms of overall bandwidth requirements. Figure 14 shows the consequences of stepping into higher precision. Engaging the secondary accumulator to any degree, which STORMM does automatically for this system starting at 30 bits of fixed precision, requires waiting for and then processing of the return value from all atomic operations, and also that the secondary accumulators be stored somewhere: the “local work unit” kernel is designed to put them in__ shared__ memory at the cost of cutting the total mapped volume of any one work unit in half. Both factors are a significant hit to performance, often more than doubling the time taken to map particle density. Performance continues to degrade as the fixed precision model enters a regime where the secondary accumulator always accepts contributions. However, the performance remains very strong relative to the “global accumulation” kernel, which can crumble when faced with similar precision requirements, and the deterministic mapping procedure in OpenMM, which uses a 64-bit integer or a similar approach with paired uint32 t accumulators depending on the hardware. For their ability to handle multiple interpolation orders and also the user’s choice of precision model, the STORMM density mapping kernels are perhaps the most versatile of any to date.

**Figure 14:**
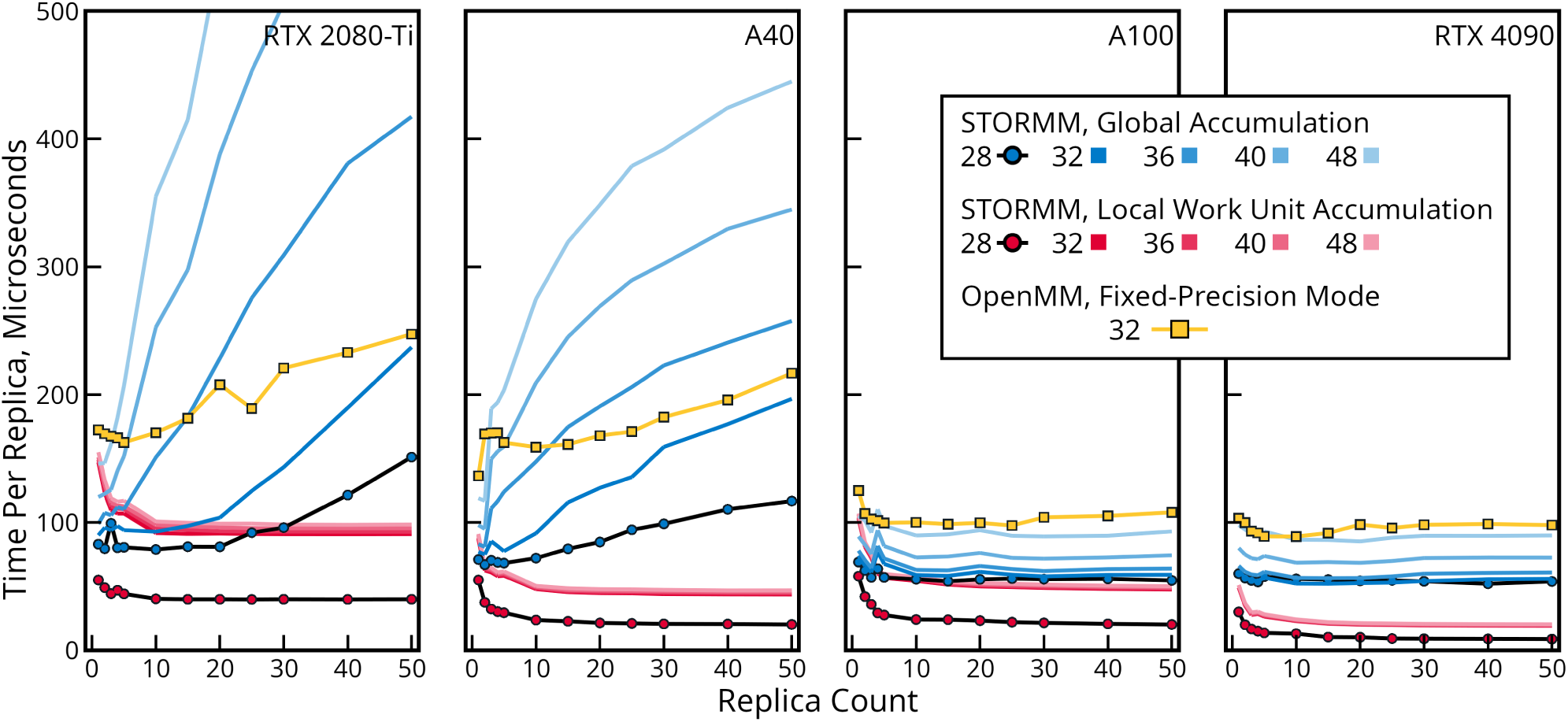
Comparison of fixed-precision particle density accumulation in STORMM and OpenMM 8.1. The AMBER GPU code also uses fixed-precision accumulation, but with 4^th^ order interpolation as opposed to the 5^th^ order methods compared above. STORMM can operate in user-specified bitwise precision, whereas all facets of OpenMM’s deterministic mode take 32 bits. STORMM calculations treat each replica of the 52,889 atom kinase simulation as distinct. For the purposes of comparison, OpenMM was tested with a tiled simulation box containing the equivalent number of replicas.

Major goals of the STORMM project include making molecular simulations more efficient and facilitating methodological advances by bringing many replicas and related systems together in the same run time instance. This requires a novel implementation, wherein lies an opportunity to reevaluate the basic allocation of effort and whether it is ideal for the evolving nature of modern GPUs. Molecular dynamics is sometimes said to scale as *O*(*P* log*P* ) in the number of particles *P* , but this reflects similar bias to statements about the problematic scaling of the FFT, a minor facet of the overall computing which exhibits weak logarithmic scaling. A more practical assessment would be that MD scales as the number of particles *P* times the cube of the particle-particle cutoff *R_C_* plus the cube of the particle-mesh interpolation order *M* , which can be counterbalanced by the cube of the inverse mesh discretization 1*/G*. The time cost explodes as any of *P* (*R_C_*)^3^ (the particle pairwise interactions), *PM* ^3^ (the particle-mesh mapping protocol), or (1*/G*)^3^ (the FFT and convolution). In practice, the valence interactions are a nontrivial but fixed cost based on the complexity of the force field and material in the system, also proportional to *P* . In the PME calculation, for any reasonable choice of *R_C_*, *M* , or 1*/G*, there is some combination of the other two parameters which can produce non-bonded forces of acceptable quality. This is the essence of STORMM’s unique coupling of the spatial decomposition to the PME mesh. Moreover, the combinations of *M* = 4 and 5*G ≈ R_C_/*2, *M* = 5 and 4*G ≈ R_C_/*2, or *M* = 6 and 3*G ≈ R_C_/*2 reflected in Figure 9 all deliver a similar level of accuracy, adequate for most chemical simulations, for a range of typical MD cutoffs (e.g. 8 Å to 12 Å). Making *R_C_* small reduces the scale of the spatial decomposition cells and drives up the mesh density, making it large drives down the mesh density: the splitting parameter controls the balance of work between particle-particle and particle-mesh interactions.

In the early days of HPC molecular simulations, demand for pairwise interactions was expected to grow based on the perception of needing lower communication bandwidth, but these involve a lot of “misses” when not all particles in one tile interact. In contrast, the particle density mapping and FFT are regular calculations where, in principle, GPUs excel. The STORMM particle-mesh setup requires that the spatial decomposition cells be cut very close to half the cutoff–if they become too large the mesh density could fall and degrade the quality of the calculated forces. Pursuant to that goal, a forthcoming paper will describe a fast method for taking the non-bonded migratory margin to zero and doing away with traditional GPU MD neighbor lists, instead updating the spatial decomposition at every step and checking the bonded exclusion status for every interacting pair. It is a radical departure from the traditional way to frame the MD problem on GPUs, but simpler in concept: at any given time, a particle will be located inside a minimal parallelepiped, with no need to consider diffusion or a margin by which each decomposition cell should be padded. There is no need to refresh a “phone book” of interacting tiles or exclusions: instead, it is all implied by the spatial decomposition and a static array of index-centered exclusion masks. This sets the stage for some surprising performance enhancements in the most memory-intensive facets of the workflow, and also straightforward methods for finding all particles in a particular vicinity for applications such as deep learning potential embedding or QM / MM boundaries.

## 6 Preliminary Applications

The original methods in STORMM have applications to relevant problems and the code base is a foundation for rapid development, but mature applications in conformer generation, docking, or molecular dynamics will take time to grow to a point where they compare to established programs and then go beyond the existing capabilities. While it is not yet possible to present mature applications, two features found in the alpha release of STORMM are detailed below. Both are built with the numerical methods and data structures detailed in previous sections. First, a large number of small molecules are subjected to combinatorial geometry permutations followed by energy-minimization *en masse*. Next, dynamics of a series of fast-folding proteins will be explored with equilibrium molecular dynamics in implicit solvent. In both cases, the GPU is hundreds to thousands of times more powerful than a single-threaded, CPU-based program, and STORMM is many times more powerful than other GPU-enabled MD programs, if they are useful to problems of such scales at all.

### 6.1 Mass small molecule geometry optimization

The energy surfaces of molecules determine the conformations that they will adopt in solution, as well as how they will interact with protein targets. While it is not in the scope of this work to compare the current methods in STORMM with an established docking or conformer generation program, the utility of solvent models and minimization are matters of debate when selecting conformers.[31, 32] Nonetheless, the relative cost of evaluating the solvation potential is much lower for a GPU than equivalent CPU code, due to an extreme increase in the relative cost of valence interactions (despite all of the progress evident in Figure 8). It is not until the number of particles *P* grows much larger that the *O*(*P* ^2^) scaling of the non-bonded calculation dwarfs the valence calculations and global reductions in the net forces required by energy minimization, as is the case for simulations of small proteins discussed in Sections 6.2 and 6.3.

Even without a strong interface to RDKit, STORMM is able to make intelligent decisions about the sampling spaces of small molecules by detecting their rotatable bonds (single bonds, not in ring systems) based on the input AMBER topologies. STORMM’s conformer application accepts inputs for the number of rotamers *N*_Rot_ about each bond to attempt to sample. While this would lead to an explosive (*N*_Rot_)*^K^* combinatorial possibilities for *K* rotatable bonds, the program can manage combinatorial sampling about individual bonds, pairs, or triplets of nearby rotatable bonds (randomizing other rotamer coordinates). It also accepts a hard upper limit to the number of rotamer combinations that should be attempted for any given molecule.

All of the selected conformations can then be submitted to energy minimization according to the chosen force field parameters (encoded in AMBER topology format), plus the options of restraints and a choice of implicit solvent models as discussed in Sections 4.4 and 3.7. STORMM uses a clash criterion to determine whether a conformation is worth subjecting to energy minimization, and to keep forces within the bounds of the fixed-precision implementation STORMM adds a “cooldown cycles” setting to the minimization namelist. As with AMBER, STORMM offers ncyc and maxcyc for the number of steepest descent and total line minimization cycles to attempt (the remainder performed by the conjugate gradient method), but cdcyc specifies the number of cycles over which to apply a softened potential, beginning with a user-specifiable degree of softening and dialing this back to zero (the natural potential is applied everywhere). There is no requirement that cdcyc be greater or smaller than ncyc. The electrostatic and van-der Waals potentials are softened by stipulating an absolute distance or some proportion of the pairwise Lennard-Jones *σ* parameter at which the potential makes a transition to a quadratic or quartic function, continuous in the first derivative, such that the softened potential reaches a maximum value for distances *r* = *−*1.0 (this can never be reached, but defines a tractable form for the softened potential). In this way, most non-bonded interactions, perhaps all of them for any reasonable configuration, behave according to the natural potential, and conflicts are resolved as the applicable range of the softened potential recedes.

Results for 93 drug-like molecules (containing 3-6 rotatable bonds, some with chiral centers or isomerizable double bonds) are summarized in Table 7, which is intended to assess the practicality of STORMM’s current features as a tool for automated generation of conformers and docked poses: the chemical model and details of the protocol will need revisions in order to produce competitive ensembles of compounds and candidates for any particular target. The FFEngine all-atom molecular mechanics force field was used to parameterize the ligands, along with different electrostatic effects. Ensembles of restraints were applied to prevent hydrogen bond donor and acceptor heavy atoms from coming closer than 2.8 Å to one another. With chiral centers or isomerizable double bonds and five or six rotatable bonds to sample, the coverage of the conformational space appears sparse, but the statistics reported in Table 7 are taken after filtering severe clashes from the initial states. Furthermore, energy minimization guides the various conformers into many of the same local minima, producing about one “unique” conformer for every three initial states. While many conformer generation protocols are said to cause candidate poses to collapse into conformations irrelevant to the state of the molecule in solution or in a binding pocket, energy-minimization under all solvent models causes a moderate increase in the radius of gyration across the test set. Restraints preventing internal hydrogen bonding appear to have a minor effect, boosting this result by about 2% across all compounds and electrostatic models (data not shown). None of the different methods show significant advantage in producing conformers closer to the bound state for each compound’s respective target ligand.

**Table 7:** Summary statistics from conformational permutations and mass energy minimizations of 93 drug candidates. Compounds had at least three rotatable bonds, sampled by computing all permutations of up to three proximate bonds with random sampling of other degrees of freedom, including double bonds outside of ring systems and chiral centers. As of this writing, STORMM does not load libraries of non-aromatic ring conformations, which would be an additional source of permutations. **(a)** The proprietary FFEngine was used to model the compounds, with electrostatics as listed. **(b)** The modified “Neck” Generalized Born model **(c)** Percentage (overall) of molecular conformations out of all possible permutations assembled by the CPU-based rotamer search, found to survive the severe clash check, and advance to energy minimization **(d)** Percentage of energy-minimized conformations distinguished by at least 0.5 Å heavy atom positional root mean-squared deviation (RMSD) **(e)** Percentage of energy-minimized conformations registering exceptionally favorable energies, i.e. because a polar hydrogen had collapsed into a neighboring atom with negative charge **(f)** Percentage of energy-minimized conformations registering very high (+1000.0 kcal/mol) scores, indicating intractable geometries such as conjoined or “stabbed” rings **(g)** Average (percentage) change in the radius of gyration *R*_gyr_ as a consequence of energy minimization. Error bars are standard deviations collected for conformations in each compound. **(h)** Minimum heavy-atom RMSD to the bound pose in each ligand’s respective target

| Potential<sup>(a)</sup> | Total Sampled Space (%)<sup>(c)</sup> | Unique Results<sup>(d)</sup> | Singularities<sup>(e)</sup> | Strained Conformations<sup>(f)</sup> | $\Delta R_{\text{gyr}}$ (%)<sup>(g)</sup> | Best RMSD<sup>(h)</sup> |
| --- | --- | --- | --- | --- | --- | --- |
| Generalized Born<sup>(b)</sup> | $1.6 \times 10^{-2}$ | 32.4% | 0.2% | 0.1% | $13.1 \pm 16.1$ | $0.83 \pm 0.36$ |
| Vacuum | $1.6 \times 10^{-2}$ | 36.3% | 0.2% | 0.2% | $14.6 \pm 18.3$ | $0.86 \pm 0.42$ |
| No Charges | $1.8 \times 10^{-2}$ | 32.6% | 0.0% | 0.1% | $12.4 \pm 14.9$ | $0.82 \pm 0.38$ |

The distances at which the softened non-bonded potentials engage may be specified in user input, but these trials underscored the importance of not softening the Lennard-Jones potential more than the electrostatic potential. Doing so can lead to singularities, in most cases involving polar hydrogens and their 1:4 neighbors or other atoms, which persist even after full restoration of the natural potential if the polar hydrogen has no Lennard-Jones character. Numerically, at the moment the forces or energies break the fixed-precision format, the minimizer can no longer make a decision on the direction of the move, leaving the system frozen in place as the ability to discern the difference between conformations at different points along the gradient line deteriorates more and more. To have the softened potentials begin at of 60% of the Lennard-Jones *σ* radii or 0.7 Å absolute distance between two charges appears to give reasonable results, but some configurations of some systems still fall into the trap. Restraints preventing internal hydrogen bonding were unable to mitigate a Lennard-Jones potential softened too far for its electrostatic counterpart (in some cases, the singularities involve polar hydrogens and 1:4 methyl carbons, which would not be protected by the restraints). The FIRE algorithm[33] or applying the RATTLE algorithm to forces would allow energy minimizations to proceed with rigid bond length constraints, which may broaden the range of acceptable onsets for the softened potentials. Both of these algorithms are planned for future development. It may also be beneficial to borrow the approach of thermodynamic integration, including its “softcore” methods, whereby the electrostatic potential remains softened until the van-der Waals potential is converged or close to its natural form.

The spread of energies and conformations obtained from this exercise does not yield many insights into the best way to obtain conformations closest to each ligand’s bound state, as illustrated in Figure 15. The FFEngine all-atom molecular mechanics force field[34] used to parameterize the ligands puts conformations of disparate geometries at similar energy levels and, given the strong anti-correlation of solvation energies with electrostatic internal energies, it may be preferable to omit both and solve conformations *in vacuo* with the van-der Waals potential and bonded interactions alone. However, to improve the conformer generator and expand into docking studies, STORMM has a nascent capability for mapping receptor potentials and shapes to different three-dimensional meshes. The capability is similar to mesh generation in the Particle-Mesh Ewald calculations described in Section 5.4, but uses separate data structures and tricubic spline interpolation rather than *B* -spline mapping. With a detailed representation of the receptor’s non-bonded potential and perhaps some mobile receptor side chains, solvation potentials and other details such as hydrogen bonding may become useful. STORMM’s multi-system energy minimization will enable high-throughput calculations within a flexible docking workflow.

**Figure 15:**
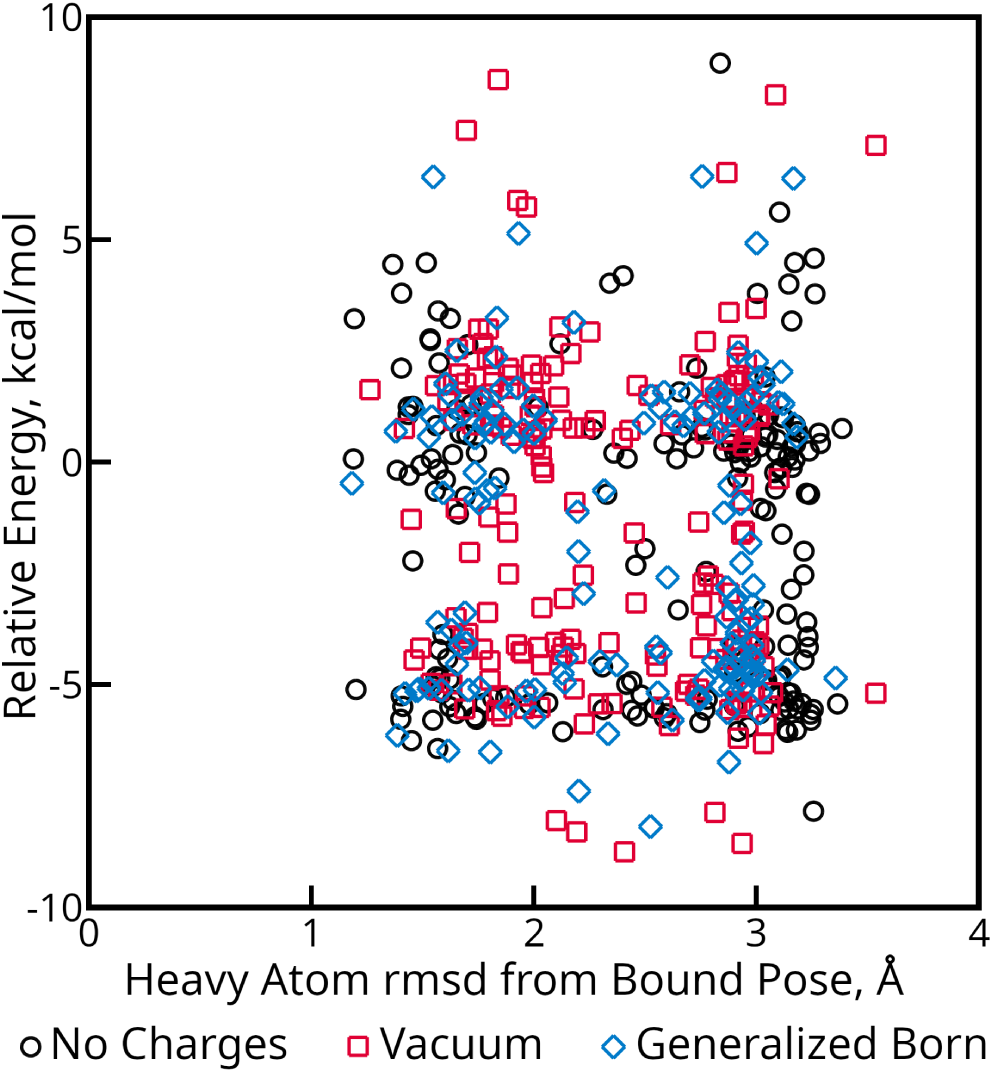
Relative energies for optimized conformations of an example drug compound. The compound in question is taken from PDB 5BVE and has six rotatable bonds, as well as some isomerization centers which bifurcate the energy surface along two apparent “tiers” in the plot.

### 6.2 Validation of a Molecular Dynamics Engine in STORMM

Molecular dynamics (MD) is a fundamental tool for sampling the conformational spaces of biomolecules, estimating binding free energies between proteins and small molecule ligands, and analyzing transport properties of bulk chemicals.[35] The latter two applications most often require condensed-phase simulations with periodic boundary conditions, but the core of an MD engine includes force calculations, an integrator for transforming forces into motion, and a means of regulating the overall amount of energy (hence, the temperature) of the system. Adding a velocity-Verlet integrator and a versatile Langevin thermostat to STORMM’s sophisticated force calculations enabled simple molecular dynamics simulations of any number of systems with isolated boundary conditions. Additional details of the implementation may be found in the Supporting Information. Applications and further developments that elaborate the dynamics engine will be published in subsequent studies.

One critical validation of any molecular dynamics integrator is its ability to conserve energy, although this may not be so simple a concept as it seems,[36] and here the notion of energy conservation is used as a means of finding errors or as a catalyst for discussion rather than an absolute metric of code quality. Simulations on six protein systems ranging in size from 219 to 918 atoms, detailed with the ff14SB force field[37] and solvated with the “Neck” GB model[20], were simulated in STORMM, OpenMM, and AMBER. The original “neck” GB model was chosen, rather than a newer variant,[38] because simulations with the newer model (“GBn2” in OpenMM nomenclature) displayed somewhat erratic energy trends in OpenMM, which this research helped trace to a bug in the GBn2 implementation for sulfur atoms. (The issue is being investigated as of the time of this writing.) Run parameters were chosen to provide each engine with favorable conditions for executing the simulations with minimal drift in the total energy: a time step of 0.5 fs, no truncation of pairwise interactions or the distance at which atoms could influence one another’s Born radii, and a constraint tolerance of 1.0 *×* 10*^−^*^7^ (close to the tightest that might be achievable with float32_t arithmetic). While many investigators have noted that constraints complicate assessment of energy conservation, this work focuses exclusively on constrained dynamics to mimic the conditions in most MD studies and also because a rigorous constraint implementation improves energy conservation by mitigating discretization errors. To further reduce high-frequency motions, the masses of atoms in the proteins were repartitioned to give hydrogens three times their natural values, despite the short time step already in place. In isolated boundary conditions, all forces are conservative and Newton’s third Law would seem to guarantee that any system will retain its net translational and rotational momentum from the beginning of the simulation, but for the fact that floating point accumulation is non-associative. Removal of net momentum can thus drain the overall energy of the system, even though it helps to prevent gradual drift in space such that its particles’ positions become less and less precise (which may accelerate energy drift due to less accurate forces). Momentum removal, and the coupled procedure of returning the system’s center of mass to the origin, were set to take place every 50 ps in AMBER and STORMM simulations, and any dynamics prior to the first momentum purge (which trims excess momentum due to the Maxwell velocity kick-start) were discarded from the analysis. The net translational momentum was removed from OpenMM simulations and at each step the system was placed back at the origin (in OpenMM’s single-precision production mode, guarding the coordinates against roundoff errors is essential). In addition, STORMM’s unique positional and velocity fixed-precision numerics (which carry units of Å and Å / fs, respectively), were given generous numbers of bits after the point to capture minute details of the step-by-step updates and constraint adjustments. Simulations were repeated in octuplicate to improve statistics.

A naive measurement of the energy conservation, the slope of the trendline for the simulation’s total energy, is presented in Table 8. As shown in Figure 16, taking such conservative run parameters, this one number is a strong indicator of the energy conservation in all three codes. In any simulation, the energy fluctuates on a short time scale and there can be stochastic events which affect the total energy. Higher-precision implementations, small time steps, constraints, and mass repartitioning to reduce the frequencies of the most rapid oscillations shift the balance further towards the systematic errors inherent to the run parameters: the naive measurement of energy conservation becomes a better description. However, the analyses may still contain hidden cancellation of errors. The simulations presented in the lower panel of Figure 16 would appear to produce better energy conservation than any of the cases in the upper panel. However, the simulation conditions are substantially worse, taking twice the time step, foregoing mass repartitioning, and using the lower-precision version of AMBER. One other detail of the upper panel in Figure 16 stands out: one of the eight trajectories simulated with STORMM’s “single-precision” settings shows a rapid jump in the total energy just after 6 ns. There is a second transition with about a quarter of this magnitude, not visible on the plot, in the STORMM “single-precision” simulations, and another in the OpenMM “single-precision” simulations. This sort of behavior helped sniff out problems with the force calculations in one GB model of OpenMM, but in that case such events were much more common and found with every OpenMM precision variant. The largest event lies in the eighth STORMM trajectory and occurs within the space of one 0.1 ps sample. While it is 12,200,000 steps into the calculation, all of STORMM’s trajectories have proven bitwise reproducible across cards of the same hardware line (e.g. all A40s, all A100s), and sometimes even across multiple card lines (data not shown). A new Watcher class is in development to monitor simulations for anomalous events subject to user specifications, which will help to characterize this and other needles in the vast haystack.

**Figure 16:**
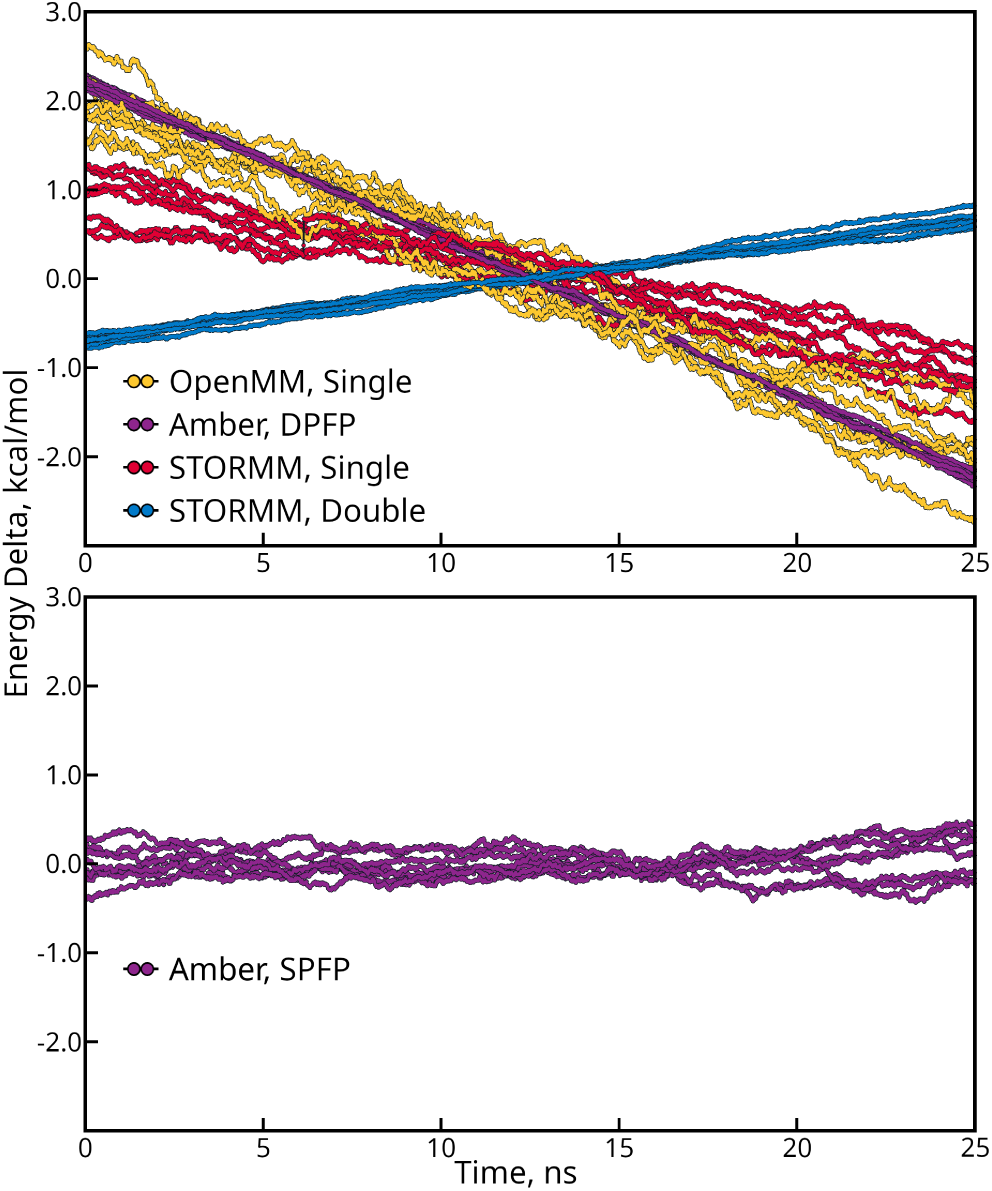
Total energy for eight trajectories of the 36-residue protein villin, produced with different codes. The “neck” GB implicit solvent model and AMBER’s ff14SB-“SC only” force field describe the system. Bond lengths to hydrogen atoms were held rigid with a tolerance of 1.0 *×* 10*^−^*^7^. Each plotted points reflects the average of 100 samples taken over a 10 ps window. In the upper panel, a 0.5 fs time step was used and masses of hydrogen atoms were repartitioned to three times their natural value. In the lower panel, a 1.0 fs time step and the natural hydrogen masses were used. As indicated in the legend, each plot is layered with a line and colored donuts: if the donuts are spaced closely enough, they fuse to take the appearance of a continuous fat line, but if the thin line beneath is ever exposed this indicates very rapid fluctuations in the total energy.

**Table 8:** Energy conservation measured as the slope of the trend line for total energy over 25 ns. Generalized Born simulations used a 0.5fs time step, no cutoffs, and a constraint tolerance of 1.0 *×* 10*^−^*^7^. Energy drift is reported as average heating or cooling (in Kelvin) based on each system’s estimated heat capacity at 300K. Error values reflect standard deviations over eight trials. In any given simulation, the instantaneous system temperature can vary, but the heat capacity is an intrinsic property that normalizes the energy drift across all system sizes. **(a)** Heat capacity is computed by a five-point stencil of the mean total energies of simulations conducted with OpenMM’s mixed-precision mode and held at 290K, 295K, 305K, or 310K with a Langevin thermostat mimicking collisions at 1.0 ps^-1^. These isothermal simulations extended for 50 ns, with the first 5 ns discarded for equilibration. Reported values and error bars reflect the mean and standard deviation of leave-two-out cross validation on eight independent stencil estimates. **(b)** Particle positions and velocities are stored in float32_t. All arithmetic is performed in float32_t. **(c)** Particle positions and velocities are stored in float64_t. Most arithmetic is performed in float32_t. Critical evaluations such as constraints and integration are carried out in float64_t. **(d)** Particle positions and velocities are stored in float64_t. All arithmetic is performed in float64_t. **(e)** Particle positions are stored to a precision of 1 part in 2^36^ of an Å, velocities to a precision of 1 part in 2^44^ of one Å-fs^-1^. Forces were accumulated with a precision of 1 part in 2^32^ of a kcal/mol-Å. All arithmetic is performed in float32_t. **(f)** Particle positions, velocities, and forces are stored to a precision of 68, 76, and 64 fixed-precision bits after the point, with the units in **(e)**. All arithmetic is performed in float64_t. STORMM does not have “single-” and “double-” precision variants: there is one executable with a variety of options available through user input.

| Protein | Conotoxin | Trp-cage,<br>tc5b | WW Do-<br>main | Villin | Protein B | Homeodomain |
| --- | --- | --- | --- | --- | --- | --- |
| Atom Count | 219 | 304 | 562 | 596 | 737 | 919 |
| Hydrogen<br>Count | 99 | 150 | 275 | 301 | 375 | 453 |
| Heat Capacity,<br>kcal/mol-K<sup>(a)</sup> | 1.28 ± 0.15 | 1.78 ± 0.17 | 2.75 ± 0.15 | 3.65 ± 0.17 | 4.14 ± 0.32 | 3.88 ± 0.23 |
| OpenMM,<br>Single<sup>(b)</sup> | -0.038 ±<br>0.011 | -0.043 ±<br>0.011 | -0.050 ±<br>0.009 | -0.044 ±<br>0.007 | -0.058 ±<br>0.008 | -0.059 ±<br>0.008 |
| OpenMM,<br>Mixed<sup>(c)</sup> | -0.037 ±<br>0.002 | -0.051 ±<br>0.002 | -0.051 ±<br>0.002 | -0.048 ±<br>0.001 | -0.062 ±<br>0.002 | -0.065 ±<br>0.002 |
| OpenMM,<br>Double<sup>(d)</sup> | -0.038 ±<br>0.002 | -0.051 ±<br>0.002 | -0.052 ±<br>0.001 | -0.050 ±<br>0.001 | -0.063 ±<br>0.000 | -0.065 ±<br>0.001 |
| AMBER,<br>SPFP<sup>(c)</sup> | -0.040 ±<br>0.005 | -0.047 ±<br>0.006 | -0.054 ±<br>0.002 | -0.047 ±<br>0.003 | -0.061 ±<br>0.003 | -0.064 ±<br>0.005 |
| AMBER,<br>DPFP<sup>(d)</sup> | -0.038 ±<br>0.001 | -0.049 ±<br>0.001 | -0.053 ±<br>0.001 | -0.049 ±<br>0.001 | -0.062 ±<br>0.001 | -0.065 ±<br>0.001 |
| STORMM,<br>Single<sup>(e)</sup> | -0.021 ±<br>0.011 | -0.025 ±<br>0.007 | -0.032 ±<br>0.007 | -0.022 ±<br>0.006 | -0.027 ±<br>0.006 | -0.037 ±<br>0.006 |
| STORMM,<br>Double<sup>(f)</sup> | 0.017 ±<br>0.002 | 0.016 ±<br>0.002 | 0.018 ±<br>0.002 | 0.015 ±<br>0.002 | 0.015 ±<br>0.001 | 0.018 ± 0.001 |

In the simulations presented in this section, the systematic errors may include discretization (high-frequency motions gather error under a finite time step), inexact computation of the constraints, and imprecise accumulation of forces. The tight tolerance of 1.0 *×* 10*^−^*^7^, near the limits or possibly beneath what is achievable in OpenMM’s “single-precision” mode due to rounding error in the float32_t coordinates, produces a weak signal but appears to be still evident in Table 8. Often, the constraints appear to impart a cooling effect on the system, almost linear in the time step for small time steps. The rate of total energy loss is well correlated (*R* = 97%) to the number of hydrogen atoms and thus constrained bonds in each system. Furthermore, the effect appears to be counterbalanced by discretization errors if the hydrogen masses are not repartitioned to reduce the frequency of hydrogen atom oscillations as shown for the villin system in Figures 16 and 18. (Note that when the bonds to hydrogen are held rigid by SHAKE or RATTLE, the next-highest frequency motion of the system is not, then, bonds between heavy atoms. As has been noted,[39] hypotheses include high-frequency collisions as well as the fact that the hydrogens can still make a rocking motion according to various bond angle force constants: three independent bond angles may affect the motion of a hydrogen atom on a typical tetrahedral carbon atom, and the sum of their effects is somewhat stiffer than any one term.) Loosening the constraint tolerance, to the default value of 1.0 *×* 10*^−^*^5^ taken by OpenMM or AMBER or higher, results in a dramatic increase of the systematic error (a rapid cooling effect), as shown again for villin in Figure 16. An aggressive 4 fs time step, which can be employed if bonds to hydrogen atoms are held rigid and the masses of those hydrogens are repartitioned, imparts stochastic energy drift but not on the order of the systematic errors arising from a loose constraint tolerance. Combining the two conditions results in a much larger energy drain, such that any of the systems studied would cool by 10 to 30 K/ns without a thermostat to inject energy from step to step (data not shown). For the marginal cost of a tighter constraint tolerance, many MD simulations taking aggressive run parameters can be conducted with much less energy escaping due to approximations and being replaced by the thermostat. However, even this precaution is of questionable worth given results presented in Section 6.4. All thermostats using typical coupling constants are capable of heating or cooling the system towards the target temperature at rates much greater than 30 K/ns, and stochastic thermostats in particular are always replacing a great deal of the system’s kinetic energy with their own.

STORMM passes a crucial test of its underlying numerics and force calculations with high marks. Its production-quality simulations display about half the energy drift of AMBER and OpenMM, even when the other codes are using double-precision arithmetic. It is notable that STORMM uses a velocity-Verlet integrator, with single-precision arithmetic in its “single”-precision mode, whereas AMBER and OpenMM’s default configuration both use the ”leapfrog” algorithm. When propagating dynamics on a system with bond length constraints, a velocity-Verlet integrator also constrains velocities of particles to eliminate movement of particles along the rigid bonds connecting them, which as shown in Table 9 will affect the conservation properties of the simulation. It would be a significant effort to graft a velocity-Verlet integrator onto AMBER, but the OpenMM package allows users to script a customized integrator with high efficiency. (A leapfrog integrator, which is simpler, faster, and more stable for the case of a constant step size, is planned for future STORMM development.) Comparing Tables 8 and 9 suggests that, even when running in complete double-precision mode, OpenMM’s velocity-Verlet integrator exacerbates the energy drift seen with the leapfrog-Verlet integrator. The velocity-Verlet integrator can also exaggerate the energy drift due to a loose constraint tolerance or an aggressive time step (data not shown).

**Table 9:**
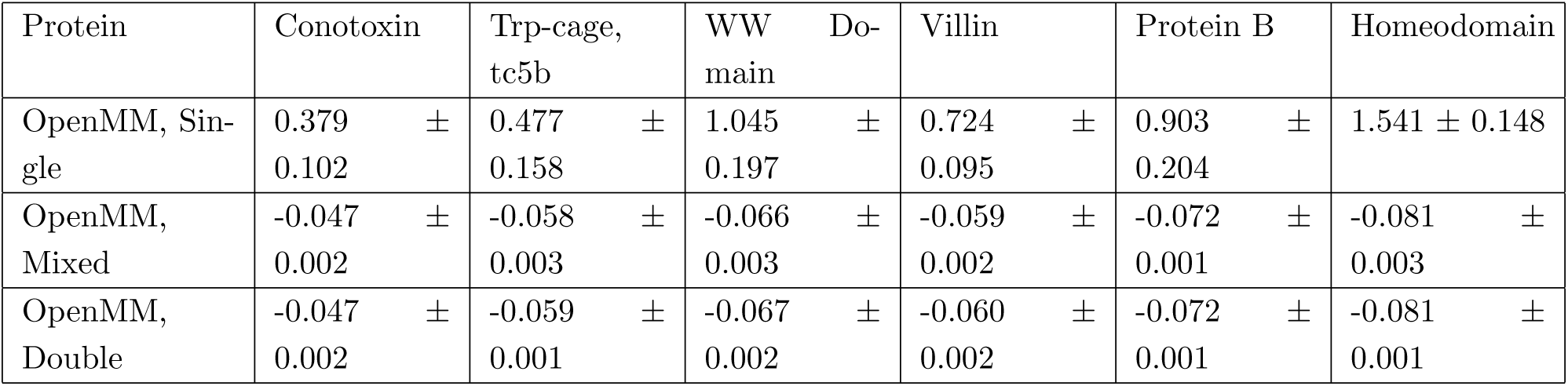
Energy conservation obtained with OpenMM 8.1 running a velocity-Verlet integrator. Energy drift in each system is expressed as a function of the heat capacity (see Table 8), in K/ns. Simulations used identical setup parameters to those in the previous table.

Other numerical issues arising from finite precision have been noted with both velocity- and leapfrog-Verlet integrators,[40] where the identified pathology of lower-precision numbers was a rapid cooling effect in the context of analytic, non-iterative SETTLE constraints (up to 170 K/ns, depending on the nuances of the integrator). Here, without high-precision integration, velocity-Verlet heats the system at ten to twenty times the rate of cooling seen in leapfrog-Verlet, even under a very conservative 0.5 fs time step, although the overall energy drift is up to 100 times smaller than what was reported in the previous study. OpenMM cannot constrain bond lengths or velocity vectors below a tolerance allowed by the underlying floating point numbers for its particle positions, which for “single-precision” mode and a protein the size of villin is about 1.0 *×* 10*^−^*^6^. Figure 17 suggests that, with a velocity-Verlet integrator, it is detrimental to try. However, OpenMM appears to be mitigating most of the consequences of float32_t positions and velocities.

**Figure 17:**
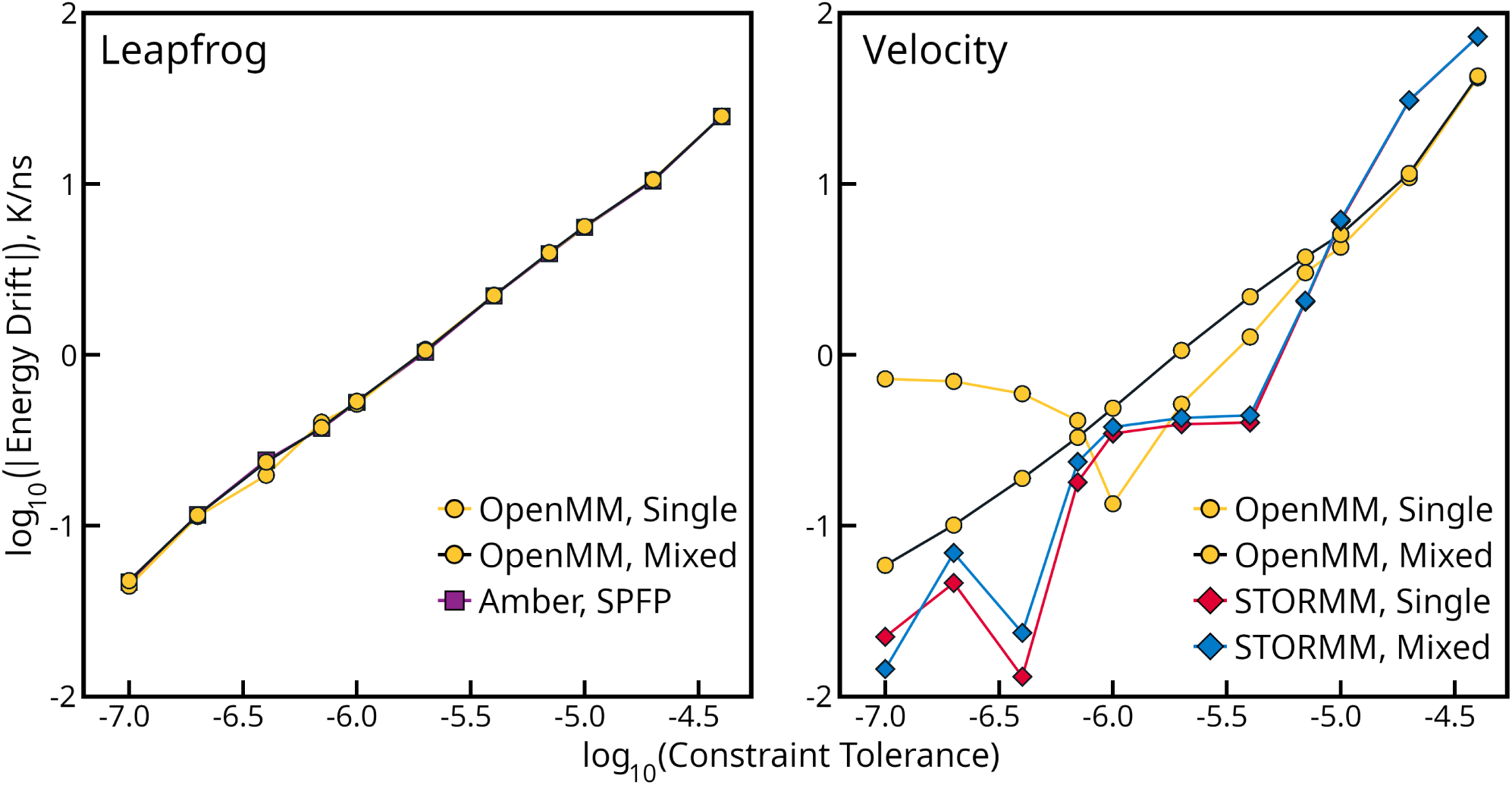
Energy drift as a function of the constraint tolerance in the villin system. In almost all cases, the energy drift is negative, but its absolute magnitude is taken to plot the result on two logarithmic axes. Results for different codes and precision models are indicated in the inset legends. Each panel presents one of the two common Verlet integrator schemes. STORMM’s “single-precision” settings follow those in Table 8. A “mixed-precision” STORMM mode was created by combining float64_t arithmetic for constraints and bonded interactions with float32_t arithmetic for non-bonded calculations using the &precision input namelist. Positional, velocity, and force accumulation in this mode were carried out with 52, 64, and 40 bits after the point, respectively.

As implied by the lower panel of Figure 16 and the right-hand panel of Figure 18, it is possible that STORMM’s “improved” energy conservation in Table 8 is due to some fortuitous cancellation of errors, such as a weak heating effect of the velocity Verlet integrator counterbalancing constraint cooling. However, when running with double-precision arithmetic and a fixed-precision model capable of capturing billions of times the level of detail one might take in production simulations, STORMM’s energy conservation shows further moderate improvement with a reversal in the sign of the drift. More significant is that STORMM’s unique method of constraining multiple bonds connected to the same central atom (described in the Supporting Information) does not produce the same cooling effect that appears to be inherent in the common method used by AMBER and OpenMM. While most codes implement “hub and spoke” constraints by moving the central atom in response to each individual bond’s update for every iteration spanning the whole group, this may bias the results towards whichever bond comes first in the topology. In contrast, STORMM pools the moves over the central atom proposed by all constraints in the group and adjusts this atom once per iteration. While it appears to yield better energy conservation for a variety of constraint tolerances up to the AMBER and OpenMM default value (see Figure 17), the cooling effect’s linearity in the constraint tolerance is broken, and this is not a consequence of performing the iterations in float32_t arithmetic: switching to float64_t arithmetic and boosting the precision of all accumulators yields the same erratic result.

**Figure 18:**
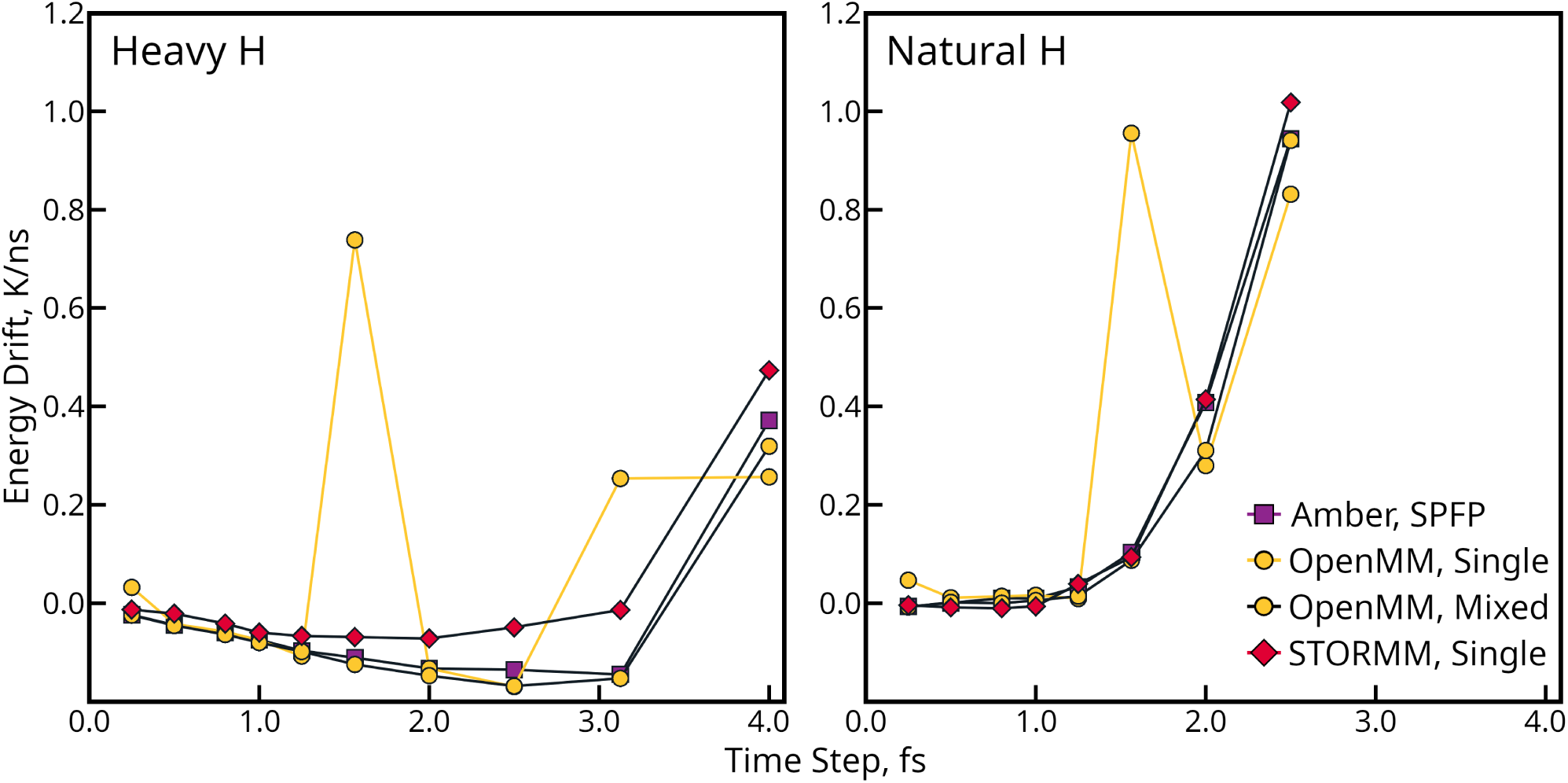
Effect of the time step size on energy drift in the villin system. Simulation protocols follow from Tabel 8, with the time step varying as indicated and, in the right hand panel, the natural hydrogen masses being used. The inset legend applies to both figures.

While pooling the moves over the central atom is the exclusive approach in STORMM’s GPU code, the CPU code implements both approaches, allowing a direct comparison. The CPU dynamics code performs all of its calculations on a single core using float64_t for arithmetic, coordinates (a PhaseSpace object holds the system, see Table 1), and most accumulations: it is not the level of precision achievable on the GPU but enough to distinguish the two constraint methods. Running the smallest conotoxin and tc5b systems in “neck” GB solvent with OpenMM and AMBER’s constraint approach leads to energy drifts of *−*0.152 *±* 0.013 K/ns and *−*0.146 *±* 0.011 K/ns, respectively, whereas the STORMM constraint approach produces energy drift at about 60-70% of that rate (*−*0.089 *±* 0.008 K/ns and *−*0.106 *±* 0.012 K/ns). There is indeed a difference between the schemes, adding weight to the proposition that STORMM’s constraint approach is the source of the improved energy conservation. However, the overall magnitudes of the drift on the CPU are higher in both cases, suggesting that differences between the CPU and GPU precision models also play a role and complicate questions as to how the sign of the drift might have flipped under STORMM’s “double-precision” settings.

To boost the signal from -RH_2_ and -RH_3_ hub-and-spoke constraint groups, a series of arginine, leucine, phenylalanine, and valine oligomers were constructed and run using the CPU code with either constraint approach, again for 25 ns in octuplicate. All N-terminal residues have the amino -RH_3_ group, while all main-chain backbones contain amide, -RH. Whereas valine residues contain dual methyl groups (-RH_3_), each arginine residue contains five -RH_2_ groups. Leucine contains both -RH_2_ and -RH_3_, and while phenylalanine contains one -RH_2_ its side chain also includes five -RH. There is some noise and perhaps non-linearity in the data, but a weighted sum of contributions from each type of group (-RH, -RH_2_, -RH_3_) yields the coefficients shown in Table 10 and fits the data very well for several common settings of the constraint tolerance. Weights for the -RH groups, which are always handled the same way, are similar in the two models for any given constraint tolerance, indicating that the regression captures the correct behavior. The coefficients for each class of constraint group suggest that, with the tight 1.0 *×* 10*^−^*^7^ tolerance, AMBER and OpenMM would display even more energy drift were it not for cancellation of errors: in these codes, a heating effect from -RH_3_ groups counteracts the cooling effect of -RH_2_ groups, but it is not as strong and in natural proteins -RH_2_ groups are more numerous, hence AMBER and OpenMM show more rapid cooling in Table 8. For various constraint tolerances, STORMM’s unbiased method almost always results in -RH_2_ and -RH_3_ groups producing much lower energy drift. While the regression is clear for a 0.5 fs time step when the dominant source of drift is the constraints, discretization errors begin to play a significant role at larger time steps and would complicate the analysis.

**Table 10:** Energy drift in oligomers of Arg, Leu, Phe, and Val decomposed by constraint groups. A linear model is proposed whereby the observed drift in each system’s total energy *E*_Total_ is expressed as a weighted sum of the number *N* of each type of SHAKE / RATTLE constraint groups: (*δ/δt*)*E*_Total_ = (*N*_H_)*α*_H_ + (*N*_H2_)*α*_H2_ + (*N*_H3_)*α*_H3_. The weights *α*_Hx_ are given in units of kcal/mol-ns. Different models are fitted for each method of applying the constraints and constraint tolerance setting, and all apply at a time step of 0.5 fs. The coefficient of determination for each model is given to the right of its weights.

| Constraint Tolerance | AMBER / OpenMM Approach |  |  |  | STORMM Approach |  |  |  |
| --- | --- | --- | --- | --- | --- | --- | --- | --- |
| | $\alpha_H$ | $\alpha_{H2}$ | $\alpha_{H3}$ | $R^2$ | $\alpha_H$ | $\alpha_{H2}$ | $\alpha_{H3}$ | $R^2$ |
| 1.0e-7 | -0.0039 | -0.0034 | 0.0026 | 0.998 | -0.0038 | -0.0003 | -0.0004 | 0.997 |
| 1.0e-6 | 0.0842 | -0.0340 | -0.1140 | 0.995 | 0.0773 | -0.0329 | -0.0531 | 0.999 |
| 1.0e-5 | -0.0390 | 0.3917 | 0.8283 | 0.983 | 0.0015 | -0.1028 | 0.2394 | 0.991 |

It could still be theorized that a cancellation of errors is taking place with STORMM’s method, particularly under the tightest constraint tolerance (1.0 *×* 10*^−^*^7^). Why would -RH_2,3_ produce a lower energy drift than -RH? The hydrogens tend to be symmetric about their central atom: perhaps STORMM’s approach counterbalances inaccuracies from the connected constraints once they are solved to high enough precision. In situations where the -RH groups are the largest problem, the issue would seem to be that the constraint algorithm does not engage *if the positions of the particles in the SHAKE group or projection of particle velocities along the bond in the RATTLE group are already within the specified tolerance*. Enforcing even one or two iterations in the -RH case, for negligible cost given the parallel processing of more complex groups, would deliver better geometries. This might stamp out a source of error and further reduce the energy drift due to constraints under very tight tolerances, although -RH does not appear to be a major source of energy drift under loose tolerances.

Energy conservation is not a sufficient criterion, or sometimes even necessary, to conduct a valid simulation. Most production simulations do not conserve energy, and to report “energy is conserved” may neglect cancellation of errors. However, given thoughtful analysis and precise run conditions, it can be fruitful to study energy conservation both as a validity check on new code and as a means for improving the underlying algorithms. The flexibility and exquisite control of numerics available in STORMM may yield other insightful research on constraints, multiple time steps, and other facets of MD integrators.

### 6.3 Performance of Implicit Solvent Molecular Dynamics in STORMM

The *O*(*N* ^2^) complexity of typical Generalized Born calculations, which either take no cutoffs or are of a size that most of their interactions fall within the cutoff anyway, implies poor scaling for small problems to the tens of thousands of threads on modern GPUs. STORMM’s capability to stack multiple systems into a single runtime process obtains excellent GPU utilization with tens, hundreds, or even thousands of concurrent simulations, as shown in Figure 19. Throughput may require many system replicas to “top out” on modern GPUs with a hundred thousand or more active threads in the major kernels’ launch grids. Maximum throughput can be better understood in terms of the total non-bonded work in these Generalized Born problems, and STORMM’s non-bonded work units offer a seamless way for systems to flow together on the GPU. Each valence work unit described in Section 5.3 is exclusive to one system, but the non-bonded work units described in the Supporting Information may incorporate tiles from multiple systems as the atom import space permits.

**Figure 19:**
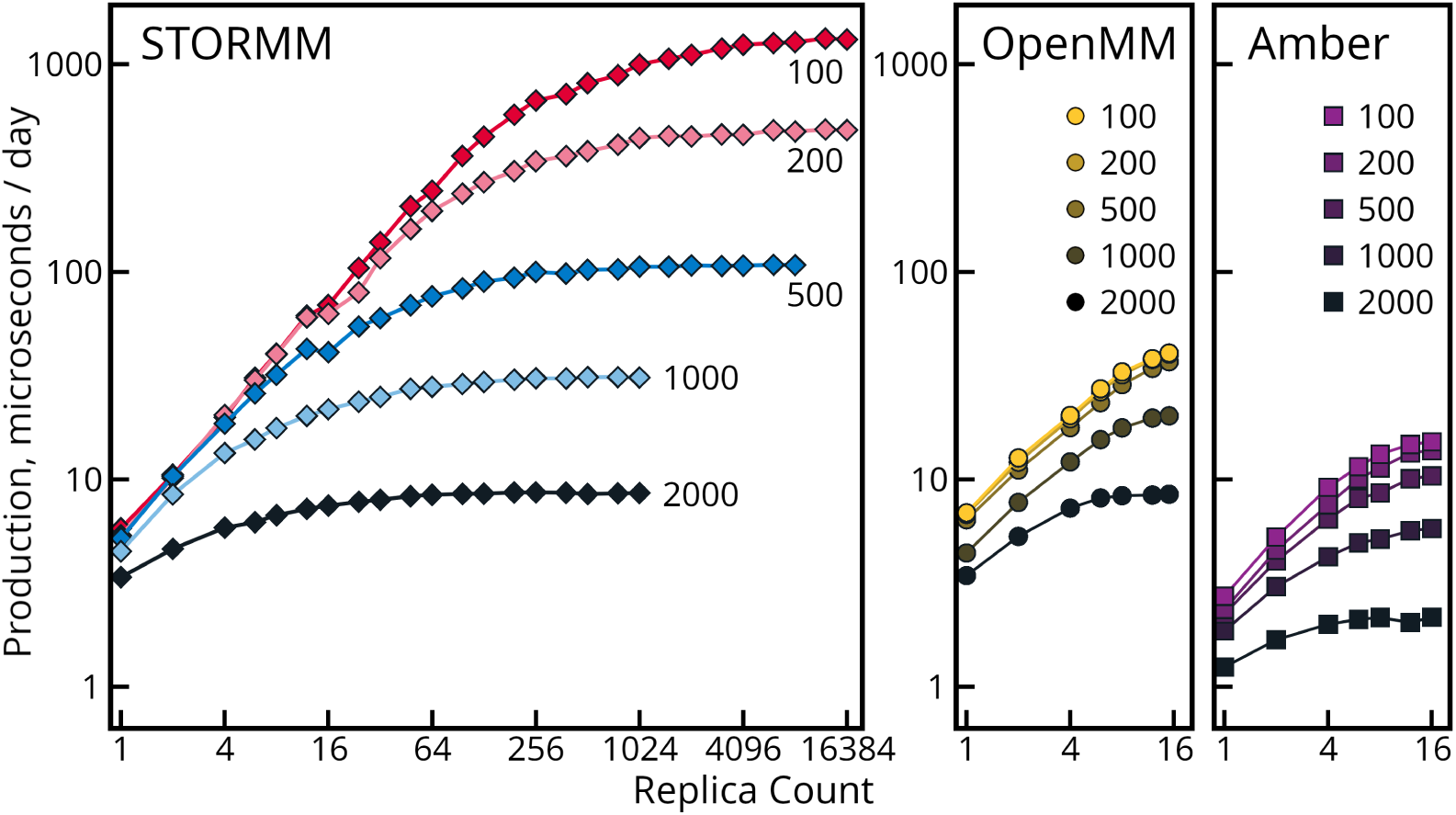
Performance running multiple replicas of small implicit solvent simulations. The “neck” GB model (igb = 7 in AMBER, GBn in OpenMM) was used to model solvent effects. Peptide chains were taken from fragments of large proteins in the PDB (see Figure 20 for systems of other sizes) and modeled with the ff14SB-“Side Chains Only” force field. Hydrogen masses were repartitioned to three times their natural value. A 4 fs time step was used with a Langevin integrator (coupling constant 1.0 / ps). System sizes are indicated on the plots or by inset legends. Plots compare the “production” modes of each code: STORMM with appropriate single-precision settings, single-precision OpenMM, and AMBER’s SPFP mode. Results for RTX 4090 are presented: other cards’ results differ in the overall scale of the throughput, which follows from the total compute capacity. STORMM calculations pertain to one runtime instance. AMBER and OpenMM calculations ran multiple concurrent executables using the NVIDIA MPS daemon.

The card’s memory is the hard limit on the number of independent systems that STORMM can run in parallel on a GPU. With NVIDIA’s **M**ulti-**P**rocess **S**ervice (MPS), AMBER and OpenMM can also run multiple systems on a single card, although in separate runtime instances. MPS supports up to sixteen concurrent processes (any more will begin to overload MPS lanes and conflict as would two concurrent jobs on the same GPU without the MPS daemon to interlace them). OpenMM throughput was found to peak at 15 jobs per GPU, whereas 16 AMBER jobs can run on one GPU through MPS. In the established codes, MPS is a means to get 5- to 6-fold throughput increases on small implicit solvent simulations. However, they do not keep up with the throughput available by running the same numbers of systems in a single STORMM runtime instance, or approach the throughput that STORMM can get on much larger collections of small problems. One additional advantage of STORMM, not shown in Figure 19, is that the program can run separate topologies and build the work units to push all simulations forward at the same rate, so many systems of 200, 500, and 1000 atoms could all be submitted as part of the same batch and all would finish at the same time, with no extra effort to backfill a job queue running under MPS to ensure optimal utilization of the card.

The scaling on single simulations is also competitive, as shown in Figure 20. In terms of numerics, STORMM’s production mode falls somewhere between OpenMM’s single-precision mode, applying constraints with float32_t arithmetic, and OpenMM’s mixed-precision mode, storing its coordinates in 64-bit values (only when higher-precision settings are engaged does STORMM access the extra 32-bit accumulators in the PhaseSpaceSynthesis). Its performance follows suit for a range of common GB problem sizes, although at higher atom counts OpenMM runs 20-25% faster as it uses a more straightforward approach whereas STORMM takes extra measures to make its non-bonded calculations more accurate and spread the calculations across more threads at low particle counts (see the Supporting Information). AMBER has the poorest performance at small atom counts, but catches up on RTX-2080Ti as the system size grows to allow more thread engagement. On Ampere and later generation cards, however, AMBER’s “Neck” GB implementations appear to deteriorate, scaling worse than *O*(*N* ^2^) even as high atom counts should permit complete thread utilization in the cards.

**Figure 20:**
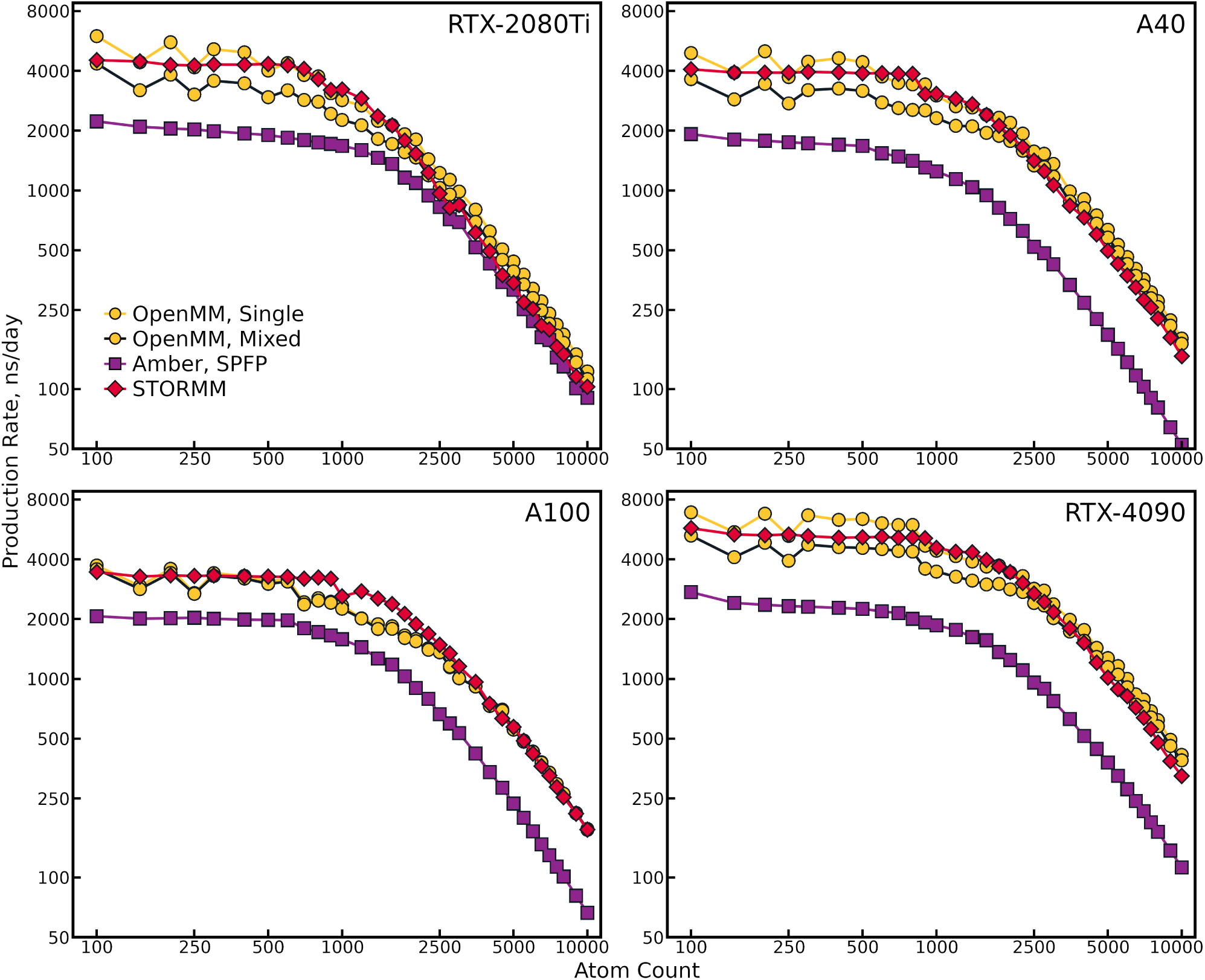
Performance of implicit solvent simulations in three different codes. Simulations were guided by the AMBER ff14SB-“Side Chains Only” force field and “Neck” GB implicit solvent. Hydrogen mass repartitioning (HMR) and a 4 fs time step were used with a Langevin thermostat (coupling constant 1.0 / ps) in AMBER and STORMM and a LangevinMiddleIntegrator in OpenMM. The inset legend of the top left panel applies to all panels.

Implicit solvent simulations of a few hundred particles could be a powerful tool for many applications in drug discovery, such as enhancing flexible docking calculations and predicting small molecule properties (e.g. solubility, aggregation, strain energy, and conformational ensembles). For general purpose molecular simulations, explicit water molecules and periodic boundary conditions are standard.

### 6.4 Comparing Trajectories from STORMM and AMBER

To assess STORMM’s molecular dynamics performance against an established code (AMBER22), the Trp-cage (tc5b) system was run using the modified “Neck” GB solvent model, a 4.0 fs time step, and temperature regulation by a Langevin integrator with a collision frequency of 1.0 ps^-1^. Hydrogen masses were repartitioned and the constraint tolerance on their rigid bonds was set to 1.0 *×* 10*^−^*^5^, following the protocol in a prior study.[38] Simulations were carried out for a total of 9.6 ms, hundreds of times longer than the original study, with either code running hundreds of separate trajectories started from either the native state (the first NMR structure in PDB 1L2Y) or an extended conformation. The first 1 *µ*s of each trajectory was discarded to decorrelate the independent simulations. Results collected with both AMBER and STORMM are in quantitative agreement with those of the prior study, as shown in Figure 21. The protein folds and unfolds many times during the simulations, and the highest populated conformation lies at about 0.8 Å root mean-squared deviation (RMSD) in the C*_α_* positions of residues 3 through 18 relative to the native state. Secondary peaks at 3.6 and 5.0 Å RMSD are also seen, again in agreement with the earlier results. Simulations initiated from both the native state and the extended conformation appear to converge to the same profile in either code, and the distributions obtained by STORMM and AMBER show no differences reaching statistical significance after subdividing each set of trajectories into sixteen parts and submitting the block averages to a two-tailed, unpaired Student’s *t* -test (*p <* 0.05). In this comparison, a “significant difference” was defined as a patch of values in the RMSD histograms wherein STORMM and AMBER trajectories show different populations when started from both native and extended conformations but neither code shows a significant difference between its own trajectories initiated from either starting point. To note, with only 300 *µ*s of simulation time, there was a difference when looking at the mean frequencies for each RMSD value, thereby warranting the more extensive simulations described above, which indicated that the distributions are in fact the same.

**Figure 21:**
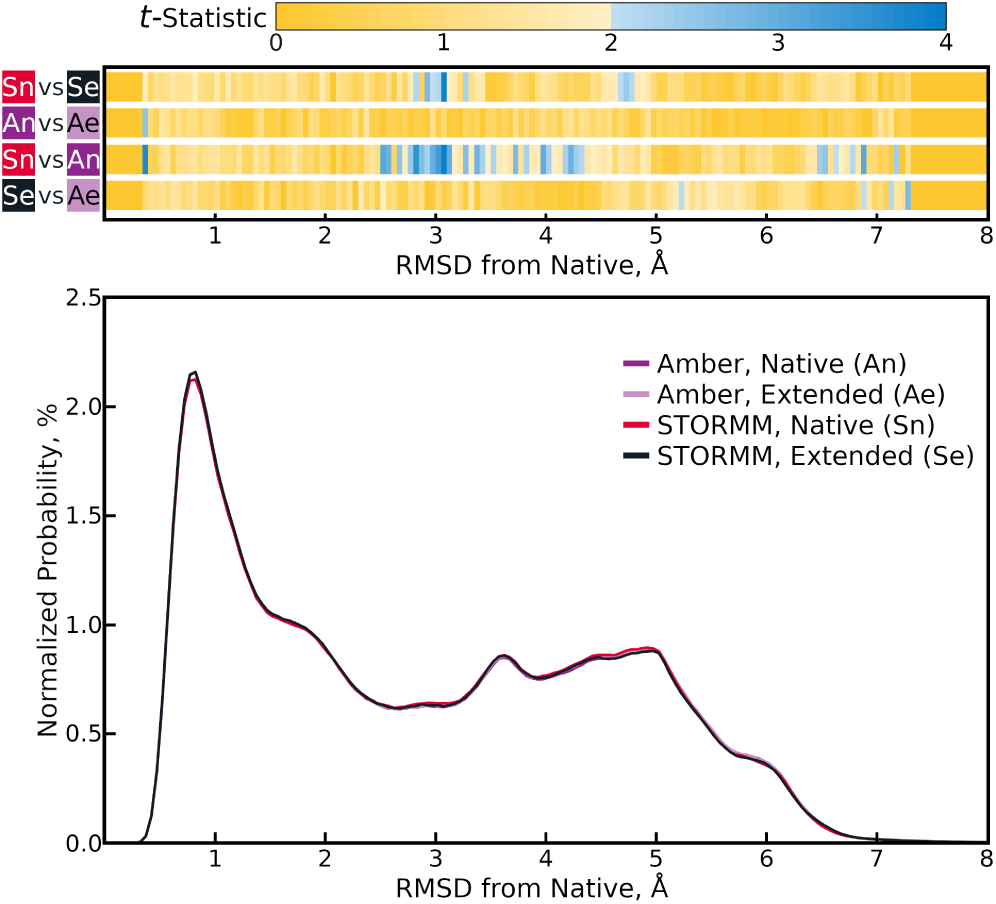
C*_α_* RMSD distributions for Trp-cage (tc5b) protein relative to the native state collected over four aggregated trajectories. Each pooled trajectory spans 2.4 ms. Histograms are normalized by the overall number of data points but not the estimated volume of states at a given RMSD (i.e. normalized by the squared value). The t-statistics obtained for comparisons between various pooled trajectories are shown in the upper panel, with the color scheme on the ordinate axis matching that in the inset legend of the lower panel. Thus, the lower two bands show the statistical significance of deviations between the abundance of structures with any given RMSD to the native state when comparing pooled trajectories generated by AMBER and by STORMM. The upper two bands provide context by showing the statistical significance of differences between pooled trajectories obtained with the same code but starting from different initial states (the “extended” or “native” tc5b conformations). To achieve 95% confidence that populations differ based on 32 independent measurements (separating any of the aggregated trajectories into sixteen blocks and comparing block averages from two trajectories), the *t* -statistic must exceed 2.037. A difference between STORMM and AMBER results would have been declared had the lower two bands both showed significance (shades of blue in the heat map) at some RMSD value while the upper two bands did not (yellow or gold in the heat map).

While STORMM’s energy conservation characteristics and the matching trajectory distributions are encouraging, questions linger from the analysis in Section 6.2: does the energy drain created by a loose constraint tolerance and an aggressive time step have any observable effect on the outcome of a simulation, and if so, do STORMM’s unique constraint iterations make any difference? To address these questions, the above protocol for tc5b was repeated once more, collecting another 4.8 ms of trajectory with STORMM for identical run conditions but taking a constraint tolerance of 1.0 *×* 10*^−^*^7^. While the resources were not available to repeat the test with AMBER, the constraint solutions are expected to converge in the limit of a tight tolerance no matter the method of iteration. Block averaging over the sixteen groups of equivalent simulations with either tolerance or starting configuration suggests that the equilibrium structural distributions are not influenced by the accuracy of the constraints: the distributions of C*_α_* RMSDs show no differences of statistical significance, nor do *χ*_1_ or *χ*_2_ distributions for any of the residues differ in any way that approaches even 95% confidence. (Following from the comparison above, the experiment sought a modest arc of the rotamer range where different constraint tolerances produce different outcomes but alternate starting conditions do not, for *p <* 0.05. No residue of tc5b satisfied these criteria.) Furthermore, the autocorrelation time associated with each variable, a basic assessment of the dynamics, suggests that there are still no significant consequences to the choice of constraint tolerance. These results, in turn, suggest that STORMM’s approach to SHAKE and RATTLE does not affect the simulation results.

The massive amount of simulation also provided a valuable “stress test” of the code base. One of the 640 STORMM trajectories taking a 1.0 *×* 10*^−^*^5^ tolerance did encounter some event, after about 1.6 billion steps, involving a very large velocity update from which the dynamics could not recover. This may be some other instance of the event noted in Figure 16, larger and more menacing in the context of production time steps and looser constraints, but it is unlikely to be indicative of a systematic problem that influences structural distributions. The event will be investigated with a range of precision settings and the forthcoming Watcher class over the course of STORMM’s alpha release, which begins with this publication.

## 7 Conclusions

The **S**tructure and **TO**pology **R**eplica **M**olecular **M**echanics (STORMM) simulation engine has been written from the ground up to best utilize powerful new GPU architectures that dominate the research computing landscape. While still in early development, STORMM already incorporates many useful features, an extensible code base, and several demonstrable improvements in the precision and performance of molecular simulations on modern GPU architectures. The paradigm of CPU-optimized work units and matching layouts of memory on the CPU and GPU is conducive to rapid development with safe handling of intricate algorithms. The versatility and control provided by a rich C++ code base is also essential for managing kernel launch decisions based on user requests and the available hardware, as simply stacking more problems together is not always beneficial to the overall throughput. Through careful algorithm design and automated arrangement of the workload, STORMM can handle many systems and simulations in the same run time process, tending towards the asymptotic performance limits of every graphics card.

Future development priorities in STORMM include a complete periodic molecular simulations capability with replica exchange[41] and weighted ensemble[42] features to allow for molecular dynamics and free energy simulations. There will also be further elaboration of the conformer generation capability to include representations of rigid and, later, semi-rigid receptors. This effort will set up a continuum of methods between conformer generation and flexible docking with a central goal being a three-dimensional modeling and pose refinement system to balance the plethora of compounds created by generative modeling approaches.[43, 44] Integral to each effort will be refinement and elaboration of the underlying coordinate and topology classes, e.g. elaboration of the coordinate objects to harness all of the Hybrid objects’ data modes, to curate a vast coordinate “reservoir” in the host memory for sequential swaps with a smaller but still substantial array of “active” molecular systems driven by a synthesis of common topologies. As planned, the periodic simulation capability will manage its neighbor list in a manner that will make it simpler to determine any and all neighbors of a given atom, whereas in most other codes the pair list only stores half the list of neighbors as an implicit guard against double-counting. Such organization will facilitate the incorporation of advanced features like subdomains treated with quantum mechanics[45] or machine learning potentials.[46] More fluid organization within STORMM will also be supported by stronger integration with external molecular modeling environments through Python bindings. The modular design of molecular objects coupled with scalable algorithms in STORMM will support a range of simulation methods for managing many independent calculations with a common theme.

## Supporting information

Additional benchmarking data for several molecular dynamics (MD) codes, plus implementation details for STORMM's MD feature

## Acknowledgment

The authors thank Peter Eastman, primary developer of OpenMM, for helpful conversations, as well as Peng Wang (NVIDIA Corporation) for technical expertise and advice. Ana Silveira (Psivant, LLC) contributed a set of drug candidates to test the conformer generator. Francesco Pontiggia (Harvard University, formerly Psivant, LLC) and Thomas Schultz (Psivant, LLC) assisted with logistical aspects of cluster resources and preparing the STORMM documentation. The authors declare no competing interests. No government funding played a part in the development of STORMM.

## References

[1] Zoe Cournia, Bryce Allen, and Woody Sherman. Relative binding free energy calculations in drug discovery: Recent advances and practical considerations. Journal of Chemical Information and Modeling, 57(12):2911–2937, 2017. PMID: 29243483.

[2] J. Harry Moore, Christian Margreitter, Jon Paul Janet, Ola Engkvist, Bert L. de Groot, and Vytautas Gapsys. Automated relative binding free energy calculations from SMILES to ΔΔ*G*. Nature Comm. Chem., 6(1), 2023. Article Number: 82.

[3] Duan Ni, Yaqin Liu, Ren Kong, Zhengtian Yu, Shaoyong Lu, and Jian Zhang. Computational elucidation of allosteric communication in proteins for allosteric drug design. Drug Discovery Today, 27(8):2226–2234, 2022.

[4] Alexios Chatzigoulas and Zoe Cournia. Rational design of allosteric modulators: Challenges and successes. WIREs Computational Molecular Science, 11(6):e1529, 2021.

[5] Andreas Krämer, An Ghysels, Eric Wang, Richard M. Venable, Jeffery B. Klauda, Bernard R. Brooks, and Richard W. Pastor. Membrane permeability of small molecules from unbiased molecular dynamics simulations. J. Chem. Phys., 153:124107, 2020.

[6] Shakhawath Hossain, Aleksei Kabedev, Albin Parrow, Christel A.S. Bergström, and Per Larsson. Molecular simulation as a computational pharmaceutics tool to predict drug solubility, solubilization processes and partitioning. European Journal of Pharmaceutics and Biopharmaceutics, 137:46–55, 2019.

[7] Chen Yang, Tong Geng, Tianqi Wang, Rushi Patel, Qingqing Xiong, Ahmed Sanaullah, Chunshu Wu, Jiayi Sheng, Charles Lin, Vipin Sachdeva, Woody Sherman, and Martin Herbordt. Fully integrated fpga molecular dynamics simulations. In *Proceedings of the International Conference for High Performance Computing, Networking*, Storage and Analysis, pages 1–31, 2019.

[8] Matt J Harvey, Giovanni Giupponi, and G De Fabritiis. Acemd: accelerating biomolecular dynamics in the microsecond time scale. Journal of chemical theory and computation, 5(6):1632–1639, 2009.

[9] Kevin J Bowers, Edmond Chow, Huafeng Xu, Ron O Dror, Michael P Eastwood, Brent A Gregersen, John L Klepeis, Istvan Kolossvary, Mark A Moraes, Federico D Sacerdoti, John K. Salmon, Yibing Shan Shan, and David E. Shaw. Scalable algorithms for molecular dynamics simulations on commodity clusters. In Proceedings of the 2006 ACM/IEEE Conference on Supercomputing, pages 84–es, 2006.

[10] Ádám Jàsz, Ádám Rák, István Ladjánszki, and György Cserey. Classical molecular dynamics on graphics processing unit architectures. WIREs Computational Molecular Science, 10(2):e1444, 2020.

[11] Ji Ding, Shidi Tang, Zheming Mei, Lingyue Wang, Qinqin Huang, Haifeng Hu, Ming Ling, and Jiansheng Wu. Vina-gpu 2.0: Further accelerating autodock vina and its derivatives with graphics processing units. Journal of Chemical Information and Modeling, 63(7):1982–1998, 2023. PMID: 36941232.

[12] Stefan Seritan, Christoph Bannwarth, Bryan S. Fales, Edward G. Hohenstein, Christine M. Isborn, Sara I. L. Kokkila-Schumacher, Xin Li, Fang Liu, Nathan Luehr, James W. Snyder Jr., Chenchen Song, Alexey V. Titov, Ivan S. Ufimtsev, Lee-Ping Wang, and Todd J. Martínez. TeraChem: A graphical processing unit-accelerated electronic structure package for large-scale ab initio molecular dynamics. WIREs Computational Molecular Science, 11(2):e1494, 2021.

[13] Nicholas Nethercote and Julian Seward. How to Shadow Every Byte of Memory Used by a Program. In VEE ’07: Proceedings of the 3rd International Conference on Virtual Execution Environments, New York, NY, USA, 2007. Association for Computing Machinery.

[14] RDKit Contributors. RDKit: Open-source cheminformatics. https://www.rdkit.org.

[15] Ivan D. Welsh and Jane R. Allison. Automated simultaneous assignment of bond orders and formal charges. Journal of Cheminformatics, 2019. PMID: 30840171.

[16] W. Zeng and R. L. Church. Finding shortest paths on real road networks: the case for A*. International Journal of Geographical Information Science, 23(4):531–543, 2009.

[17] Dylan M. Anstine and Olexandr Isayev. Generative models as an emerging paradigm in the chemical sciences. Journal of the American Chemical Society, 145(16):8736–8750, 2023. PMID: 37052978.

[18] Ulrich Essmann, Lalith Perera, Max L. Berkowitz, Tom Darden, Hsing Lee, and Lee G Pedersen. A smooth particle mesh Ewald method. Journal of Chemical Physics, 103:8577–8593, 1995.

[19] Mark James Abraham, Teemu Murtola, Roland Schulz, Szilárd Páll, Jeremy C. Smith, Berk Hess, and Erik Lindahl. Gromacs: High performance molecular simulations through multi-level parallelism from laptops to supercomputers. SoftwareX, 1-2:19–25, 2015.

[20] John Mongan, Carlos Simmerling, J. Andrew McCammon, David A. Case, and Alexei Onufriev. Generalized Born model with a simple, robust molecular volume correction. J. Chem. Theory Comput., 3(1):156–169, 2007. PMID: 21072141.

[21] Alexei Onufriev, Don Bashford, and David A. Case. Exploring protein native states and large-scale conformational changes with a modified generalized born model. Proteins, 55(2):383–394, 2004.

[22] Gregory D. Hawkins, Christopher J. Crame, and Donald G. Truhlar. Pairwise solute descreening of solute charges from a dielectric medium. Chem. Phys. Lett., 246(1):122–129, 1995.

[23] Peter Eastman, Jason Swails, John D. Chodera, Robert T. McGibbon, Yutong Zhao, Kyle A. Beauchamp, Lee-Ping Wang, Andrew C. Simmonett, Matthew P. Harrigan, Chaya D. Stern, Rafal P. Wiewiora, Bernard R. Brooks, and Vijay S. Pande. OpenMM 7: Rapid development of high performance algorithms for molecular dynamics. Public Library of Science Computational Biology, 13(7):e1005659, 2017.

[24] Tai-Sung Lee, David S. Cerutti, Dan Mermelstein, Charles Lin, Scott LeGrand, Timothy J. Giese, Adrian Roitberg, David A. Case, Ross C. Walker, and Darrin M. York. GPU-Accelerated Molecular Dynamics and Free Energy Methods in Amber18: Performance Enhancements and New Features. Journal of Chemical Information and Modeling, 58(10):2043–2050, 2018. PMID: 30199633.

[25] Matthias Buck, Sabine Bouguet-Bonnet, Richard W. Pastor, and Alexander D. MacKerell. Importance of the cmap correction to the charmm22 protein force field: Dynamics of hen lysozyme. Biophysical Journal, 90(4):L36–L38, 2006.

[26] Roger W. Hockney and James W. Eastwood. Computer Simulation Using Particles. Taylor & Francis, 1988.

[27] David J. Hardy, Zhe Wu, James C. Phillips, John E. Stone, Robert D. Skeel, and Klaus Schulten. Multi-level summation method for electrostatic force evaluation. Journal of Chemical Theory and Computation, 11(2):766–779, 2015. PMID: 25691833.

[28] W. Michael Brown, Axel Kohlmeyer, Steven J. Plimpton, and Arnold N. Tharrington. Implementing molecular dynamics on hybrid high performance computers – Particle–particle particle-mesh. Comput. Phys. Comm., 183(3):449–459, 2012.

[29] David S. Cerutti and David A. Case. Multi-level ewald: A hybrid multigrid/fast fourier transform approach to the electrostatic particle-mesh problem. Journal of Chemical Theory and Computation, 6(2):443–458, 2010. PMID: 22039358.

[30] David S. Cerutti, Robert E. Duke, Thomas A. Darden, and Terry P. Lybrand. Staggered Mesh Ewald: An Extension of the Smooth Particle-Mesh Ewald Method Adding Great Versatility. Journal of Chemical Theory and Computation, 5(9):2322–2338, 2009. PMID: 20174456.

[31] Andrew T. McNutt, Fatimah Bisiriyu, Sophia Song, Ananya Vyas, Geoffrey R. Hutchison, and David Ryan Koes. Conformer generation for structure-based drug design: How many and how good? Journal of Chemical Information and Modeling, 63(21):6598–6607, 2023.

[32] Paul C. D. Hawkins, A. Geoffrey Skillman, Gregory L. Warren, Benjamin A. Ellingson, and Matthew T. Stahl. Conformer Generation with OMEGA: Algorithm and Validation Using High Quality Structures from the Protein Databank and Cambridge Structural Database. Journal of Chemical Information and Modeling, 50(4):572–584, 2010.

[33] Erik Bitzek, Pekka Koskinen, Franz Gähler, Michael Moseler, and Peter Gumbsch. Structural relaxation made simple. Phys. Rev. Lett., 97:170201, Oct 2006.

[34] Sheenam Khuttan, Solmaz Azimi, Joseph Z. Wu, Sebastian Dick, Chuanjie Wu, Huafeng Xu, and Emilio Gallichio. Taming multiple binding poses in alchemical binding free energy prediction: The beta-cyclodextrin host–guest SAMPL9 blinded challenge. Phys. Chem. Chem. Phys., 25:24364–24376, 2023.

[35] Edward J. Maginn, Richard A. Messerly, Daniel J. Carlson, Daniel R. Roe, and J. Richard Elliot. Best Practices for Computing Transport Properties 1. Self-Diffusivity and Viscosity from Equilibrium Molecular Dynamics. Living Journal of Computational Molecular Science, 1(1):6324, 2018.

[36] Peter Eastman and Vijay Pande. Energy Conservation as a Measure of Simulation Accuracy. BioRxiv, 2015.

[37] James A. Maier, Carmenza Martinez, Koushik Kasavajhala, Lauren Wickstrom, Kevin E. Hauser, and Carlos Simmerling. ff14sb: Improving the accuracy of protein side chain and backbone parameters from ff99sb. Journal of Chemical Theory and Computation, 11(8):3696–3713, 2015. PMID: 26574453.

[38] Hai Nguyen, Daniel R. Roe, and Carlos Simmerling. Improved generalized Born solvent model parameters for protein simulations. J. Chem. Theory Comput., 9(4):2020–2034, 2013. PMID: PMC4361090.

[39] Ron Elber, A. Peter Ruymgaart, and Berk Hess. SHAKE parallelization. Eur. Phys. J., 200(1):211–233, 2011.

[40] Ross A. Lippert, Kevin J. Bowers, Ron O. Dror, Michael P. Eastwood, Brent A. Gregersen, John L. Klepeis, Istvan Kolossvary, and David E. Shaw. A common, avoidable source of error in molecular dynamics integrators. The Journal of Chemical Physics, 126(4):046101, 2007.

[41] Rafael C. Bernardi, Marcelo C.R. Melo, and Klaus Schulten. Enhanced sampling techniques in molecular dynamics simulations of biological systems. Biochimica et Biophysica Acta (BBA) - General Subjects, 1850(5):872–877, 2015.

[42] Daniel M. Zuckerman and Lillian T. Chong. Weighted Ensemble Simulation: Review of Methodology, Applications, and Software. Annu. Rev. Biophys., 46:43–57, 2017. PMID: 28301772.

[43] Vendy Fialková, Jiaxi Zhao, Kostas Papadopoulos, Ola Engkvist, Esben Jannik Bjerrum, Thierry Kogej, and Atanas Patronov. Libinvent: Reaction-based generative scaffold decoration for in silico library design. Journal of Chemical Information and Modeling, 62(9):2046–2063, 2022. PMID: 34460269.

[44] John B. Ingraham, Max Baranov, Zak Costello, Karl. W. Barber, Wujie Wang, Ahmed Ismail, Vincent Frappier, Dana M. Lord, Christopher Ng-Thow-Hing, Erik R. Van Vlack, Shan Tie, Vincent Xue, Sarah C. Cowles, Alan Leung, João V. Rodrigues, Claudio L. Morales-Perez, Alex M. Ayoub, Robin Green, Katharine Puentes, Frank Oplinger, Nishant V. Panwar, Fritz Obermeyer, Adam R. Root, Andrew L. Beam, Frank J. Poelwijk, and Gevorg Grigoryan. Illuminating protein space with a programmable generative model. Nature, 623:1070–1078, 2023.

[45] C.E. Tzeliou, M.A. Mermigki, and D. Tzeli. Review on the QM/MM Methodologies and Their Application to Metalloproteins. Molecules, 27(9):2660, 2022. PMID: 35566011.

[46] Peter Eastman, Raimondas Galvelis, Raúl P. Peláez, Charlles R. A. Abreu, Stephen E. Farr, Emilio Gallicchio, Anton Gorenko, Michael M. Henry, Frank Hu, Jing Huang, Andreas Krämer, Julien Michel, Joshua A. Mitchell, Vijay S. Pande, João PGLM Rodrigues, Jaime Rodriguez-Guerra, Andrew C. Simmonett, Sukrit Singh, Jason Swails, Philip Turner, Yuanqing Wang, Ivy Zhang, John D. Chodera, Gianni De Fabritiis, and Thomas E. Markland. Openmm 8: Molecular dynamics simulation with machine learning potentials. The Journal of Physical Chemistry B, 128(1):109–116, 2024. PMID: 38154096.

