## Additional benchmarking data for several molecular dynamics (MD) codes, plus implementation details for STORMM's MD feature for "STORMM: Structure and TOpology Replica Molecular Mechanics for chemical simulations"

https://www.overleaf.com/project/64d021ed1bcc1a91c05f549c
ALT TITLE: STORMM chemical simulation software

### Supporting Information: Further details of molecular dynamics in Amber, OpenMM, and STORMM

David S. Cerutti[1], Rafal Wiewiora[1], Simon Boothroyd[1], and Woody Sherman[1]

[1]Psivant Therapeutics, Boston, Massachusetts 02210, United States

March 27, 2024

#### 1 Some known benchmarks

Most MD codes publish numbers to demonstrate their performance on a handful of systems. In Table 1, the Amber benchmarks DHFR (referred to as "JAC" on the Amber website), FactorIX, Cellulose, and STMV were all run in the constant volume, constant temperature ensemble with a 4fs time step typical of production MD for simple equilibrium simulations. These runs are the source of the "$\mu$s per step" figures reported in the main text, by simple unit conversion. In the main text, lower time per step is better, whereas here, higher ns/day is better.

Amber22 was used to compile these benchmarks. The version of OpenMM used to compile these benchmarks is a post-release modification of the 8.1 code, developed by Peter Eastman after this research revealed that some pair list block optimizations in 8.1 were sensitive to the topological layout of the system. As a result, the 8.1 release sees slowdowns of up to 30% on the FactorIX benchmark and 10-20% on the STMV benchmark. A new version will be released with the corrections tested herein, and was available to the public but not official as of the time of this writing. The changes do not affect the time taken for valence computations or charge density mapping detailed in the main text.

#### 2 A closer look

The four systems shown in Table 1 offer only a glimpse of the performance, even in typical equilibrium MD. The systems span a large range of sizes with few data points, and the contents of each system are inconsistent, as hinted at in the table's caption.

For a more complete picture, the DHFR benchmark was tiled up to 100 times, using the most compact arrangement possible for each number of tiles, to make super-boxes and simulate these systems of up to 2.36 million atoms. In whatever degree of replication, the DHFR system is 90% water, putting it towards the low end of protein content for protein-in-water simulations. The DHFR topology in the original Amber benchmark was created some 20 years ago and its exact force field was difficult to trace, let alone reproduce in the current builders. The tiled systems were therefore modeled in the contemporary ff14SB force field,[1] with the same atom-for-atom content. In addition, a protein crystal of a scorpion venom toxin adapted from earlier work [2] was tiled to create a super-cell of four crystallographic unit cells, a total of 24,674 atoms in all. Again, the protein content was modeled with ff14SB while acetate and ammonium ions were modeled with parameters developed in the earlier work (also available on the Amber website Tutorials section, "Simulation of Protein Crystal").

| System | DHFR$^{(a)}$ | FactorIX$^{(b)}$ | Cellulose$^{(c)}$ | STMV$^{(d)}$ |
|---|---|---|---|---|
| RTX 2080-Ti | | | | |
| AMBER ns/day | 835 | 301 | 65.4 | 22.6 |
| OpenMM ns/day | 1166 | 313 | 54.2 | 17.4 |
| A40 | | | | |
| Amber ns/day | 890 | 367 | 83.5 | 29.5 |
| OpenMM ns/day | 1455 | 452 | 64.9 | 21.7 |
| A100 | | | | |
| Amber ns/day | 1108 | 501 | 145.7 | 51.8 |
| OpenMM ns/day | 1350 | 496 | 128.5 | 42.2 |
| RTX 4090 | | | | |
| AMBER ns/day | 1383 | 671 | 185.6 | 69.1 |
| OpenMM ns/day | 2135 | 825 | 199.0 | 65.6 |

Table 1: Timings for AMBER22 and a post-release modification of OpenMM 8.1 running four AMBER benchmark MD systems. Run conditions: 4fs time step, constant volume, temperature maintained at 300K by Langevin thermostat, 9.0 Å cutoff on short-ranged electrostatic and Lennard-Jones interactions, TIP3P water. PME was applied only to electrostatics, with each program's default behavior determining the grid density.
**(a)** Dihydrofolate reductase, 23558 atoms (including 21069 in water)
**(b)** Factor IX, 90906 atoms (including 85074 in water)
**(c)** Cellulose fibers (in water), 408609 atoms (including 317565 in water)
**(d)** Satellite Tobacco Mosaic Virus, 1067095 atoms (including 900159 in water)

While comparable in size to the basic unit of DHFR tiling, 65% of this system's atoms were found in protein and polyatomic ions, matter bearing dihedral interactions and other valence terms. Like the DHFR unit, the 24,674 atom block was replicated up to 100 times to create simulations of up to 2.47 million atoms. These two cases bracket the material content of biomolecular simulations in terms of high and low water content: while rigid TIP3P water molecules, used throughout this study, require cheaper computations, water has a higher diffusion coefficient and can thus affect pair list rebuilding times. The degree to which these two effects balance each other is uncertain, but the toxin crystal's particle density is one per 9.94 Å$^3$ in comparison to the DHFR simulation's density of one particle per 9.12 Å$^3$, implying a 30% increase in the number of pairwise interactions to compute for the same 9.0 Å cutoff. The content as well as the size of the simulation both have bearing on its cost.

Figure 1 shows the relative speeds of each each code when given many systems sampling the range of sizes on a finer (and broader) scale than the four benchmarks in Table 1. The isolated examples become curves, and it is apparent that each code requires a certain number of atoms, given the GPU and the system content, to saturate the compute capacity. The first major result, contrary to much perception in the community, is that OpenMM is now faster than Amber22, at least in terms of equilibrium MD, for the most useful system sizes (e.g. 150,000 atoms or less) on NVIDIA's consumer-grade gaming cards. Only on the data-center grade A100, where the memory bandwidth is much higher, does Amber excel at lower atom counts, and even then OpenMM and Amber remain competitive up to 150,000 atoms when Amber finally wrests the lead. In all cases, Amber wins as the systems get very large, but OpenMM appears to have made considerable strides in recent years and the crossover point grows with the size of the card: for the most recent RTX 4090, OpenMM is the clear winner for systems as large as 250,000 or even 500,000 atoms. Amber's valence interaction computations are likely a major contributor, as discussed in the main text. The other basic result is that both codes display a degree

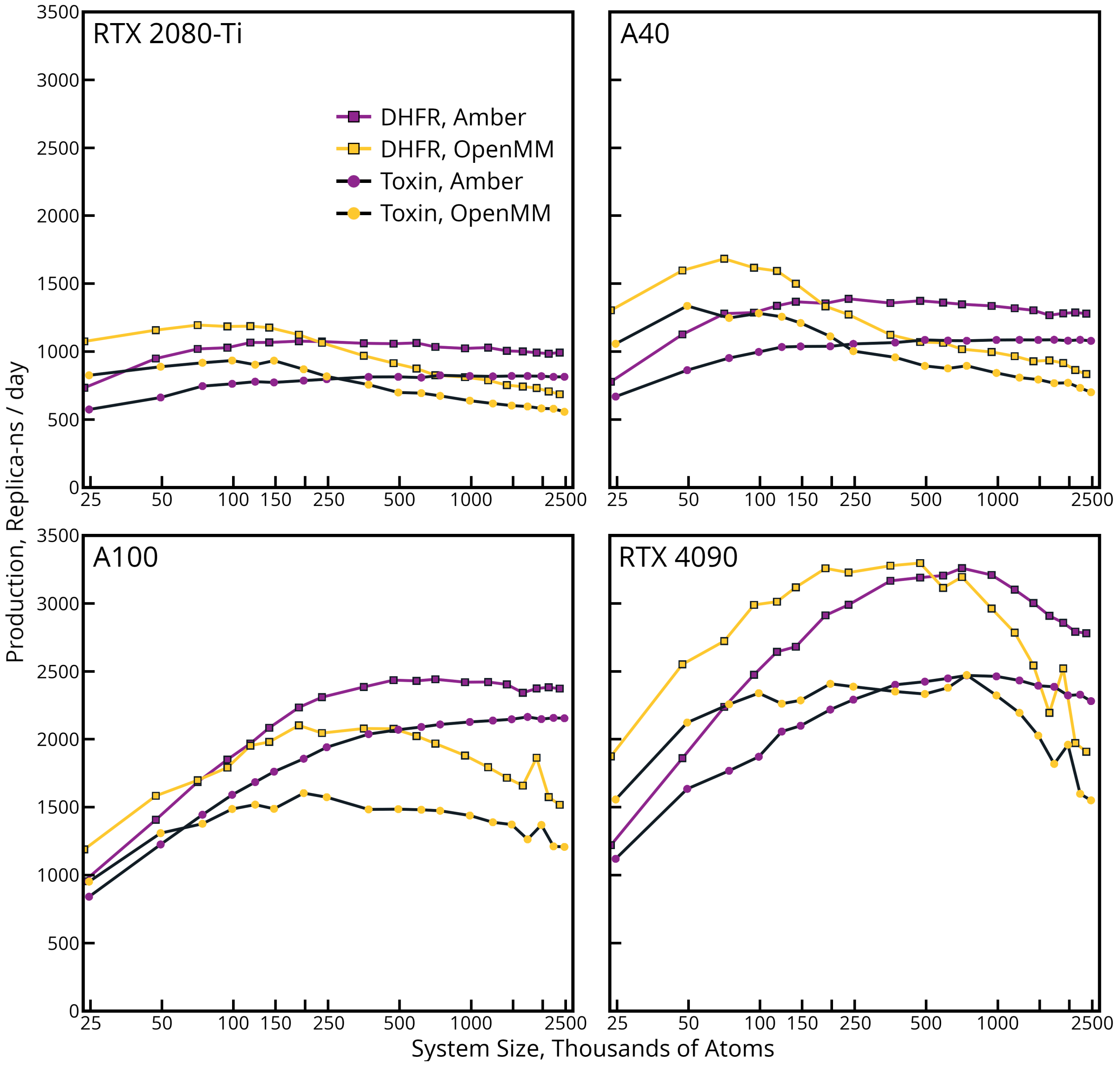


Figure 1: **Amber22 and a post-release modification of OpenMM 8.1 running on systems of varied content and a range of sizes.** The $x$-axis is plotted on a logarithmic scale. The $y$-axis shows the production per replica-time, i.e. the overall production (ns/day) of a simulation with 47,116 atoms and two replicas of DHFR is multiplied by two, to present all simulations in a convenient metric.

of performance regression after reaching their peak throughputs. Amber does so to much less of a degree than OpenMM, but it nonetheless runs into memory bottlenecks or becomes overtaken by nonlinear effects such as the degradation in particle density mapping discussed in the main text or the cost of pair list rebuilding (Amber and OpenMM both see the cost rise, but a mechanistic understanding of each code suggests that the reasons differ).

The DHFR benchmark may not be representative of most MD simulations, for a number of reasons. First, the protein has a charge of +11 a.u., though this is not balanced by any counterions. The math is fine: PME clips the excess charge by convention. The periodic boundary conditions imply that there might be pressure for the independent protein copies in the tiled systems to shift into a new packing arrangement but not repel each other in a significant way. However, it is still not representative of most simulation setups, and lacking counterions probably helped this system avoid performance regression in OpenMM 8.1 (the problems arose based on the Amber builder's ordering of the topology, in particular water and ions). Also, close inspection of the results in Figure 1 for the single replica case, which is the DHFR benchmark but taking a different protein model, shows that Amber22 simulates the system 6-9% slower and OpenMM simulates it 2-7% slower when the protein is modeled with ff14SB as opposed to the dated topology used in the official benchmark. This effect is seen on all GPUs tested. The number of dihedral interactions increases some 40% in the newer model. It might be explained by the Amber valence work units running through an extra round of dihedral interactions for a given group of atoms and OpenMM having to issue more atomics to the same atoms for similar reasons. Whatever the cause, the fact that two independent codes show similar and significant effects based on the choice of chemical model suggests that these aspects of a benchmark are important, and that the common DHFR benchmark needs revision.

As always, benchmarking comes with caveats that must be communicated and understood. Most significant here is that, while OpenMM and Amber are both running in their default production modes (single-precision with non-deterministic charge density accumulation for OpenMM, single-precision with fixed-point determinism for Amber), the precision models are not the same. OpenMM probably has an overall advantage given the way it handles coordinates in this mode, and its "mixed" precision model, where most computations are carried out in `float32_t` but accumulation is mediated by `float64_t`, might be a better comparison. However, the effect is probably not enough to erase OpenMM's advantage except near the crossover points in Figure 1. Furthermore, most users will opt for the fastest available option, and both codes are able to manage simulations with reasonable energy conservation. Another feature that may have helped Amber gain a perceived advantage in years past has also fallen away in recent versions: when running on Volta and Turing GPUs with CUDA 10 and below, Amber was compiled with instructions for compute capability 6.x combined with whatever architecture-specific code optimizations (e.g. `-gencode arch=compute_60,code=sm_86` for an A40). This trick, which turns off independent thread rescheduling, was offered to give old codes a grace period for porting to CUDA's new "Simultaneous Instruction, Multiple Threads" paradigm and instead retain the older warp-synchronous programming model. While the grace period *continues to be extended*, it was thought that it was to expire in CUDA 11 and therefore Amber stopped taking this compile option. OpenMM, in contrast, has always adopted current NVIDIA compiling recommendations, so the two codes are, since 2020, on equal footing in this regard. For well-written code with well-coalesced warps, however, foregoing independent thread rescheduling on NVIDIA hardware can confer 10 to 15% improved performance. This has been seen in STORMM (data is not shown, and the optimization is not taken), and was noted by Amber users switching to CUDA 11 on Turing and Volta-generation hardware. The temporary use of this compiling option may have given Amber an undocumented advantage between 2017 and 2020. Further progress in OpenMM, however, makes its developmental version the clear winner for most standard MD applications.

### 3 Compiling STORMM's unique capabilities into a novel MD implementation

The complete STORMM molecular dynamics implementation has several innovations that are described in this supplement so that the main article could focus on the most noteworthy and exemplary features. While the dynamics implementation is in its infancy (written in the three months before this work was submitted), it maintains the versatile and modular design of the underlying components. Attention is given to each detail of the non-bonded calculation, particle movement, and thermo-regulation to ensure that the high standards of the basic libraries translate into reliable simulations.

In its all-to-all non-bonded loops, STORMM uses an unusual method for splitting NVIDIA's 32-lane warps in half to process a $16 \times 16$ tile as illustrated in Figure 2. The approach is to have the 16 lowest-index thread lanes read atoms along one edge of the tile, for which they then have sole ownership. The 16 highest-index lanes read atoms along the other tile edge, for which they also have sole ownership. Pairwise interactions are evaluated by having each thread of the low or high lanes read atom data from registers of a corresponding thread in the high or low lanes (this is the `__shfl()` instruction in CUDA). Each thread then stores the computed force affecting its own atom, but the equal and opposite force does not go to waste: threads cannot write to one another's registers, but they can calculate the index of the thread lane that read their own atom, and then query the force computed there. In this manner, 32 threads in the CUDA warp compute all interactions in the $16 \times 16$ tile in eight interaction cycles. Calculating the thread lane from which to draw coordinates, and afterwards the lane from which to extract forces, requires a degree of integer arithmetic, whereas all commodity cards tested in this paper are built to deliver more floating-point arithmetic. However, the strategy reduces register pressure by eliminating extra copies of the atom coordinate and property information that would otherwise have to be passed around as the tile cycles. Devoting each warp to a $16 \times 16$ tile also applies up to four times as many threads to a given number of pairwise interactions as a scheme using $32 \times 32$ atom tiles, but at the expense of higher bandwidth to and from `__global__` memory. As with other aspects of the force calculation, STORMM produces work units to import collections of atoms and then evaluate all possible tiles among them, which mitigates the bandwidth issue but imposes its own cost in terms of synchronization across the thread blocks. We set the thread block size at eight warps to balance between the cost and benefit.

As explained in the Table 1 of the main text, STORMM stores all of its positions and particle velocities in fixed-precision with a user-specified number of bits after the point. The central motivation is to ensure that displacements between particles can always be calculated to a given precision, regardless of the floating-point type used for real-valued arithmetic. While consumer-grade graphics cards tend to have poor throughput on `float64_t` arithmetic, all NVIDIA cards have good throughput on addition or multiplication with `int64_t` numbers. While it would be most accurate to calculate the displacements between every interacting pair in fixed-precision and then convert these numbers to floating point, the conversion is expensive. STORMM's design entails a compromise for the non-bonded tiles: the average value of all particles' positions in the tile is reduced across the warp so that the interacting tile's center of geometry can be moved to the origin, minimizing the floating point values of all positions to conserve bits in their mantissas. Some tiles, e.g. those with atoms on opposite sides of a large protein, will have sprawling configurations, but because the atoms along each side of the tile are taken from consecutive topological entries, these tiles are likely to contain two close-knit groups of particles very far from one another. Their pair interactions will be weak. The adjustment adds a cost to the overall dynamics cycle, but may be a plank in STORMM's excellent energy conservation.

Numerical precision, reproducibility, and stability are paramount in STORMM, and these criteria are also reflected in the design of the random number generators. STORMM's class of integrators comes with built-in thermostating capabilities, and in the case of a Langevin or Andersen thermostat a separate Xoshiro256++

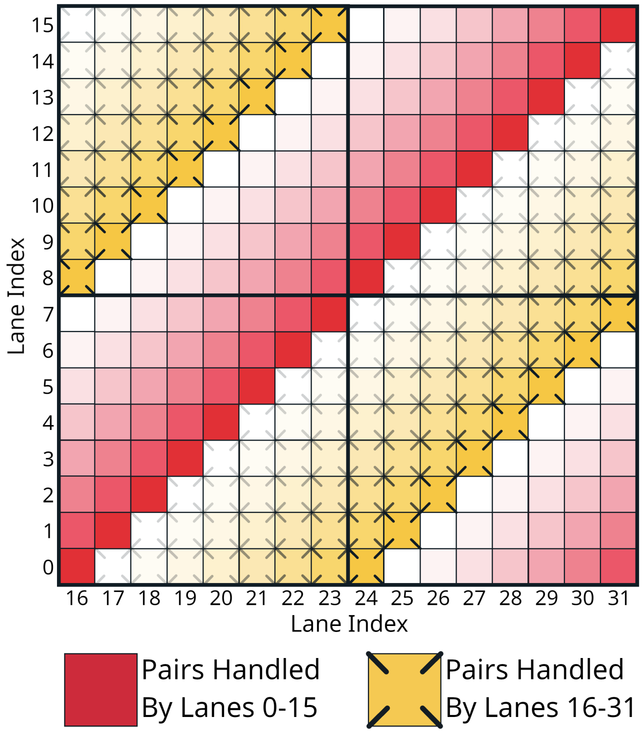


Figure 2: **Order of pair interactions in a tile with half the vector processing width.** The illustration applies to a $16 \times 16$ tile appropriate for NVIDIA's 32-threaded "warp." A similar strategy could create a $32 \times 32$ tile for the 64-threaded "wave front" in AMD's consumer-grade cards. Red and yellow crossed squares indicate the pairs handled by threads in the first and second halves of the thread group, respectively, with the intensity of the color fading in subsequent iterations.

generator[3, 4] state is allocated for each atom in the synthesis of all systems. The initial states for these generators are taken by jumping one seeded state forward by the "long stride" of $2^{192}$ cycles, ensuring that the random sequences affecting each atom will never overlap. STORMM also implements the Xoroshiro128+ generator, the 128-bit states of which require only half the memory of the 256-bit Xoshiro256++ engine, but Xoroshiro128+ fails the BigCrush test[5]. Furthermore, it is not recommended to run more than a few thousand separate streams of Xoroshiro128+ and presume them free of correlations. Each of these generators create 64-bit random strings, the high bits of which are of the highest random quality. Uniform random numbers are then generated in `float32_t` or `float64_t` by taking the high 24 or 53 bits of the string, respectively, and creating a floating-point number in the range [0, 1). To generate random numbers on a normal distribution, the Box-Muller transform[6] is applied, but computing the logarithm of $1.0 - x$ for $x$ taken from a uniform distribution in `float64_t` precision even if subsequent square root or trigonometric operations are carried out in `float32_t` precision to return a `float32_t` result. Careful treatment of the logarithm improves signal in the tails of the distribution, for which there must be a probability of sampling the uniform distribution at values much smaller than the $\epsilon = 2^{-24}$ for a `float32_t` in the range [0, 1). Computing $1 - x$ flips the range to (0, 1] to preclude an undefined logarithm if $x = 0$, but also limits the logarithmic argument to the aforementioned $\epsilon$. Furthermore, there must be a provision for sampling values of $1 - x$ very close to 1: $log(1 - x) \rightarrow x$ as $x \rightarrow 0$, hence $\mathrm{logf}(1 - \epsilon)$ for `float32_t` yields a value of $\epsilon$ and the square root of this, the closest available number to zero other than zero itself, is rather large ($\sim$ 0.00034527). By working in `float64_t` until the logarithm is evaluated, the most likely values of the distribution receive smooth sampling as well. Uniform random numbers with only 24 bits of detail always provide a coarse sampling of the center of a normal distribution and, depending on how they are generated, can also be inadequate for sampling the tails. It may be possible to adapt the methodology in Section 5.2 ("Logarithmic Indexing for Tabulated Functional Forms") of the main text to develop spline tables for the Box-Muller transform based on uniform random numbers computed with 32 to 40 bits while performing all arithmetic in `float32_t` for cards that do not support 64-bit logarithms, or in situations where speed is of the essence, but the limited application of "double-precision" math in our Box-Muller approach does not appear to make a noticeable change to the MD timings as a whole on consumer-grade cards. The built-in XOR-shift generators and their vectorization also enable the CPU to predict the random sequence that any atom will encounter on a GPU, although more effort will be needed before CPU-based simulations can perfectly track GPU simulations.

The XOR-shift generators are so efficient that the problem is communication rather than computing the numbers themselves. Both OpenMM and Amber launch a dedicated kernel to cache arrays of random numbers as needed for all atoms, but this is more about simplicity and, in Amber's case, the fact that the CUDA random libraries are leveraged for production runs. STORMM also pre-computes random numbers in one kernel and stores them for use in another, but in larger quantities if the user specifies it: to pull each atom's random state vector out of memory and then replace the modified state to continue on the sequence amounts to 32 bytes read and written back to the card's RAM, likely for the purpose of creating random `float32_t` values for each Cartesian axis (a total of only 12 bytes). By storing more numbers and making less frequent checkouts of the generators, the memory traffic can be enriched in random number products rather than the generator states, although this can overwhelm the memory bus or engage too few threads at a time if not woven into some other kernel with low bandwidth requirements of its own. It is a similar strategy to OpenMM's separate streams for the memory-intensive PME convolutions and compute-intensive pairwise interactions, but implemented in the code rather than with card-level features. For implicit solvent MD, the best options were kernels for GB radii computations or, in vacuum simulations, the non-bonded pairwise tiles. For the forthcoming PME implementation, random number generation may be assigned to the particle-mesh mapping kernel described in Section 5.4 ("Particle-mesh interactions in periodic simulations") of the main text. Assignments for refreshing the random number cache arrays are written into the non-bonded work units that STORMM creates for the system at startup, alongside directives to initialize force accumulators for subsequent steps. This setup works well in combination with the integrator being fused to the valence interaction kernel, as illustrated in Figure 6 of the main text: vacuum dynamics with a Langevin or Andersen thermostat can be carried out with just two kernel launches for the entire MD step, while implicit solvent simulations involve only four kernel launches. Rescaling approaches such as Berendsen or Nose-Hoover thermostats, which require a reduction over all particles, demand additional kernel launches (it is not guaranteed that one valence work unit will subsume all particles), more round-trips of particle details to and from `__global__` memory, and would likely take longer.

One other unique facet of STORMM's MD implementation is its handling of constraints. While the SETTLE algorithm[7] will be implemented to handle rigid water molecules once periodic dynamics are implemented for systems in the condensed phase, the RATTLE algorithm[8] is implemented for the release version at the time of this writing. Codes such as Amber and OpenMM handle constraints of multiple particles all bonded to a central atom but not to one another ("hub and spoke" groups) by assigning one group to each thread. STORMM assigns one thread to each constrained bond and incorporates information into its work units indicating which groups of threads are participating in the same hub-and-spoke group. While Amber and OpenMM iterate by constraining each bond of the group in sequence, moving the distal atoms and more massive central atom with each adjustment, in STORMM threads move each of the distal atoms and then cooperate with other threads working on the same hub-and-spoke group (all of which are guaranteed to be in the same warp by the design of the instruction sets) to reduce and broadcast their proposed moves of the central atom. In this manner, there is no dependence on the order in which the bonds appear in the constraint instruction, although the Amber and OpenMM approach converges over slightly fewer iterations as shown by CPU-based testing code (see Table 2). Better parallelism is likely to be a speed bonus of its own, despite the extra communication surrounding the central atom update, and delegating one thread to each constrained bond enables any number of rigid bonds to connect to a single atom without raising the kernel's register pressure. This approach may be the most amenable to GPUs, as matrix-based methodologies such as $P$-SHAKE,[9] while they also move all atoms based on common reference positions with each iteration, can entail very high register pressure. In typical MD applications, bond length constraints are executed as part of the fused valence work unit kernel, along with other aspects of the integration. In addition, STORMM's GPU implementation makes use of the fixed-precision coordinate representation in the `PhaseSpaceSynthesis` to capture minute adjustments in the constrained positions or velocities when running calculations in `float32_t`. In contrast, OpenMM's "single-precision" production mode cannot converge constrained bonds as tightly if they are further from the coordinate origin, and Amber relies on its `float64_t` reference coordinates to implement constraints.

| SHAKE | | | | | | |
|---|---|---|---|---|---|---|
| | STORMM Approach | | | Typical Approach | | |
| Tolerance | 1 Bond | 2 Bonds | 3 Bonds | 1 Bond | 2 Bonds | 3 Bonds |
| 1.0e-7 | 6.71 ± 0.75 | 8.28 ± 1.09 | 9.70 ± 0.88 | 6.71 ± 0.75 | 7.29 ± 0.62 | 7.70 ± 0.57 |
| 1.0e-6 | 5.31 ± 0.78 | 6.39 ± 0.96 | 7.59 ± 0.79 | 5.31 ± 0.78 | 5.80 ± 0.58 | 6.21 ± 0.56 |
| 1.0e-5 | 3.85 ± 0.75 | 4.55 ± 0.81 | 5.48 ± 0.72 | 3.86 ± 0.75 | 4.32 ± 0.58 | 4.75 ± 0.52 |
| RATTLE | | | | | | |
| | STORMM Approach | | | Typical Approach | | |
| Tolerance | 1 Bond | 2 Bonds | 3 Bonds | 1 Bond | 2 Bonds | 3 Bonds |
| 1.0e-7 | 6.87 ± 0.79 | 8.37 ± 1.12 | 9.75 ± 0.87 | 6.86 ± 0.79 | 7.37 ± 0.62 | 7.73 ± 0.59 |
| 1.0e-6 | 5.45 ± 0.78 | 6.49 ± 0.99 | 7.63 ± 0.79 | 5.45 ± 0.78 | 5.88 ± 0.60 | 6.24 ± 0.58 |
| 1.0e-5 | 4.02 ± 0.78 | 4.64 ± 0.83 | 5.52 ± 0.73 | 4.02 ± 0.79 | 4.40 ± 0.60 | 4.78 ± 0.54 |

Table 2: **Iterations of SHAKE or RATTLE required to converge hub-and-spoke constraint groups in the Protein B system.** Iterative algorithms for STORMM's unique approach and the "typical" approach found in Amber and OpenMM were carried out using STORMM's CPU implementation in `float64_t` arithmetic and coordinates. The number of samples for each test (840,000 to 2.54 million) is large enough that all comparisons, save those between iteration methods with constraint groups with only one bond, are statistically significant. Executing RATTLE consistently takes 0.03 to 0.17 more iterations than SHAKE on the same groups of atoms, and STORMM's iteration method for groups with two or three bonds takes 0.23 to 2.02 more iterations than the typical approach, the largest differences appearing in triple-bonded groups with very tight tolerances.

### References


[1] James A. Maier, Carmenza Martinez, Koushik Kasavajhala, Lauren Wickstrom, Kevin E. Hauser, and Carlos Simmerling. ff14sb: Improving the accuracy of protein side chain and backbone parameters from ff99sb. *Journal of Chemical Theory and Computation*, 11(8):3696–3713, 2015. PMID: 26574453.

[2] David S. Cerutti, Peter L. Freddolino, Robert E. Jr. Duke, and David A. Case. Simulations of a protein crystal with a high resolution x-ray structure: Evaluation of force fields and water models. *The Journal of Physical Chemistry B*, 114(40):12811–12824, 2010.

[3] David Blackman and Sebastiano Vigna. Scrambled linear pseudorandom number generators. *ACM Trans. Math. Softw.*, 47(4), sep 2021.

[4] Sebastiano Vigna. It is high time we let go of the mersenne twister, 2019.

[5] Daniel Lemire and Melissa E. O'Neill. Xorshift1024*, xorshift1024+, xorshift128+ and xoroshiro128+ fail statistical tests for linearity. *Journal of Computational and Applied Mathematics*, 350:139–142, 2019.

[6] George Edward Pelham Box and Mervin Edgar Muller. A note on the generation of random normal deviates. *Annals of Mathematical Statistics*, 29:610–611, 1958.

[7] Shuichi Miyamoto and Peter A. Kollman. Settle: An analytical version of the SHAKE and RATTLE algorithm for rigid water models. *J. Comput. Chem.*, 13(8):952–962, 1992.

[8] Hans C. Andersen. Rattle: A "velocity" version of the shake algorithm for molecular dynamics calculations. *Journal of Computational Physics*, 52(1):24–34, 1983.

[9] Pedro Gonnet. P-shake: A quadratically convergent shake in o(n2). *Journal of Computational Physics*, 220(2):740–750, 2007.
